# Genomic prediction of heterosis, inbreeding control, and mate allocation in outbred diploid and tetraploid populations

**DOI:** 10.1101/2024.07.01.601581

**Authors:** Jeffrey B. Endelman

## Abstract

Breeders have long appreciated the need to balance selection for short-term genetic gain with maintaining genetic variance for long-term gain. For outbred populations, the method called Optimum Contribution Selection (OCS) chooses parental contributions to maximize the average breeding value at a prescribed inbreeding rate. With Optimum Mate Allocation (OMA), the contribution of each mating is optimized, which allows for specific combining ability due to dominance. To enable OCS and OMA in polyploid species, new theoretical results were derived to (1) predict mid-parent heterosis due to dominance and (2) control inbreeding in a population of arbitrary ploidy. A new Convex optimization framework for OMA, named COMA, was developed and released as public software. Under stochastic simulation of a genomic selection program, COMA maintained a target inbreeding rate of 0.5% using either pedigree or genomic IBD kinship. Significantly more genetic gain was realized with pedigree kinship, which is consistent with previous studies showing the selective advantage of an individual under OCS is dominated by its Mendelian sampling term. Despite the higher accuracy (+0.2–0.3) when predicting mate performance with OMA compared to OCS, there was little long-term gain advantage. The sparsity of the COMA mating design and flexibility to incorporate mating constraints offer practical incentives over OCS. In a potato breeding case study with 170 candidates, the optimal solution at 0.5% inbreeding involved 43 parents but only 43 of the 903 possible matings.

## INTRODUCTION

Plant and animal breeders have long appreciated the need to balance selection for short-term genetic gain with maintaining genetic variance for long-term gain. For outbred populations, inbreeding is an effective proxy for the loss of genetic variance over time. A simple expression to capture this idea exists for random mating populations: for inbreeding coefficient *F*, the additive genetic variance is 1–*F* times the base population variance (Falconer and Mackay 1996). Even when plant varieties are released as inbred lines or F1 hybrids (e.g., wheat, maize), long-term management of genetic diversity by inbreeding control is still relevant with respect to an outbred parental population, which may be defined theoretically or used in practice (Gaynor et al. 2017).

Because individuals with high genetic merit are often related, truncation selection leads to matings between close relatives. This phenomenon is exacerbated with selection on pedigree or genomic BLUP, and because of its long-term use in animal breeding, there is a significant body of research on optimizing selection under restricted inbreeding (Wray and Thompson 1990; Toro and Pérez-Enciso 1990; Wray and Goddard 1994; Meuwissen 1997; Grundy et al. 1998). In the standard methodology known as optimum contribution selection (OCS), the decision variables are the genetic contribution of each parental candidate to the next generation, and the objective is to maximize genetic merit at a target inbreeding rate, typically 0.5–1.0% (Sonneson et al. 2012; Woolliams et al. 2015). Inbreeding rate represents the probability of “new” inbreeding per generation and is inversely related to the effective population size (Falconer and Mackay 1996).

OCS has several well-known limitations. It assumes random mating, but practical mating plans often require constraints based on the reproductive biology of the species, such as the number of offspring per mating or the number of matings per individual. Because these constraints can make it difficult to realize the optimum contributions, there is an advantage to optimizing the contributions and mating plan simultaneously, which has been called optimum mate allocation (OMA) (Kinghorn 2011; Akdemir and Sánchez 2016; Gorjanc and Hickey 2018). Shifting from OCS to OMA also allows the specific combining ability of each mating to be considered when computing genetic merit (Toro and Varona, 2010).

A practical motivation for the present study was to implement OMA in a tetraploid potato (*S. tuberosum*) breeding program. Several new theoretical results were needed to realize this goal. Diploid formulas for inbreeding rate are not correct for autopolyploids because the inbreeding coefficient of an individual is not equal to parental kinship; it also depends on the parental inbreeding coefficients (Kerr et al. 2012). There is also no published formula for the genomic prediction of mate performance (GPMP) in polyploids, defined as the F1 progeny mean with directional dominance; previous studies have used look-up tables, limiting theoretical insight (Labroo et al. 2023).

From a computational perspective, an attractive feature of OCS is the *convex* nature of the problem, which implies the global optimum can be found efficiently (Boyd and Vandenberghe 2004). This feature has been exploited in previous OCS studies (Pong-Wong and Woolliams 2007; Wellman 2019), but OMA implementations have not been explicitly convex, relying instead on evolutionary optimization algorithms (Holland 1992). I report here on the formulation of Convex OMA and associated software, named COMA. This new methodology was evaluated using stochastic simulation of a simple breeding program, as well as real potato breeding data.

## THEORY

### F1 progeny mean

Our single-locus, bi-allelic model is based on the decomposition of genotypic value into the population mean plus additive and digenic dominance values (Endelman 2023). (Higher order dominance for polyploids is neglected.) The additive (*u*_*i*_) and dominance (*v*_*i*_) values for individual *i* are the products of regressors and regression coefficients:

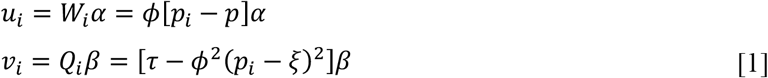

where 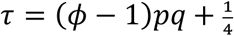 and 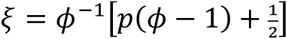

Additive regressor *W*_*i*_ is a linear function of the difference between the individual allele frequency *p*_*i*_ and the population allele frequency 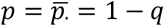. The proportionality constant, or slope, is the ploidy *ϕ* (Fig. 1A). From standard quantitative genetics, the additive regression coefficient *α* is the allele substitution effect at Hardy-Weinberg (i.e., panmictic) Equilibrium, HWE (Falconer and Mackay 1996).

**Figure 1.**
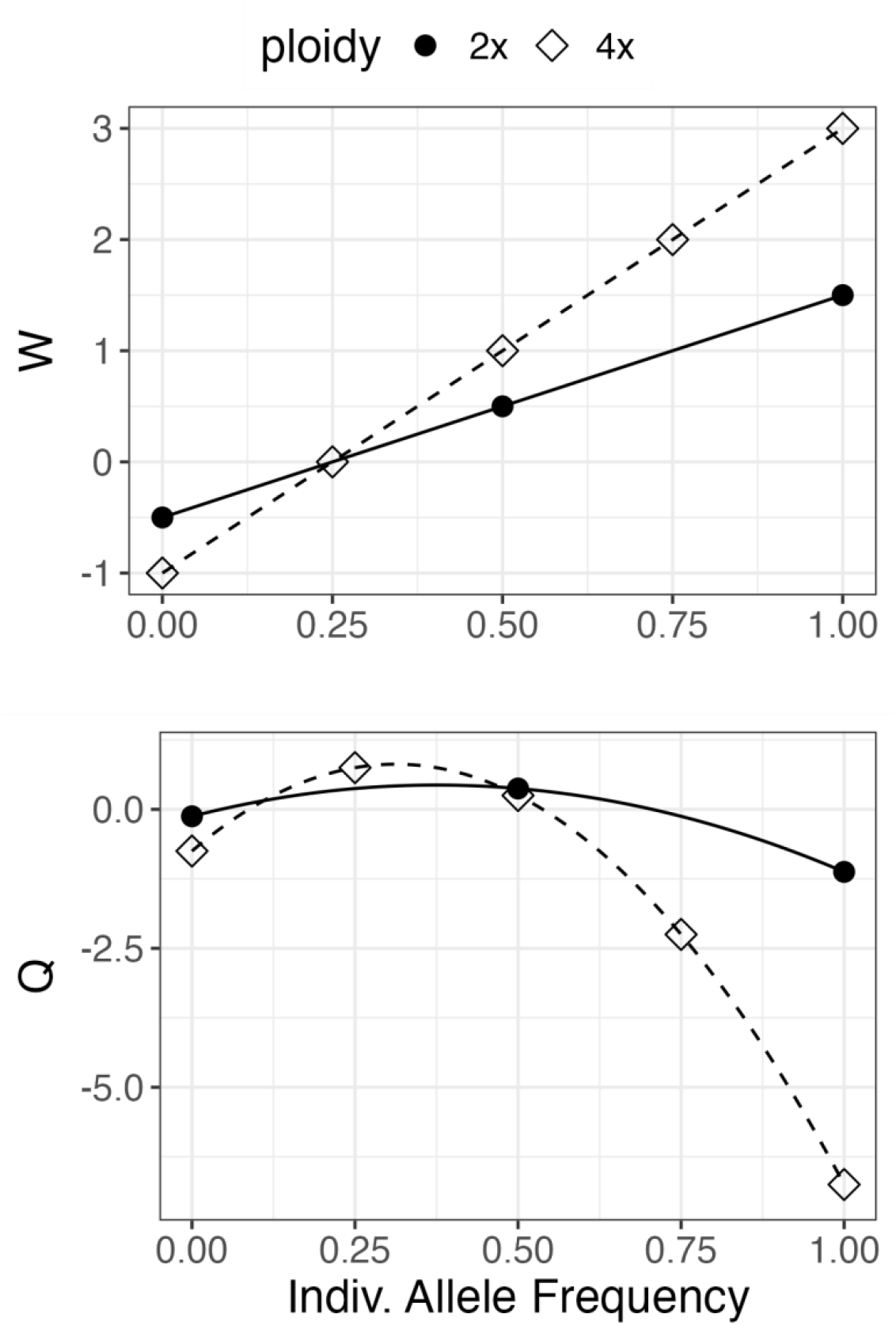
The additive (W) and dominance (Q) regressors as linear vs. quadratic functions, respectively, of individual allele frequency (dosage/ploidy). The population frequency is ¼.

Dominance regressor *Q*_*i*_ is a quadratic function of the difference between *p*_*i*_ and *ξ*, a parameter that depends on *p* and *ϕ*. From the inverted parabolic shape of *Q*, homozygotes have lower dominance value than heterozygotes, as expected (Fig. 1B). The curvature of the parabola equals ploidy squared. The dominance regression coefficient *β* is the digenic substitution effect at HWE; in other words, it is the difference between the effect of the heterozygous diplotype and the mean effect of the homozygous diplotypes (Endelman 2023). For diploids, Eq. 1 reduces to the classical “breeding value” parameterization with *β* = *d* (Vitezica et al. 2013). The symbol *β* is used to emphasize it is the extension of *α* to digenic effects.

Consider F1 progeny of the mating between individuals 1 and 2 with allele frequencies *p*_1_ and *p*_2_, respectively. Because additive values are linear functions of allele dosage, the mean progeny additive value equals the mid-parent additive value: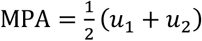. The contribution of dominance effects is the sum of the mid-parent dominance value (MPD) and mid-parent heterosis (MPH), where

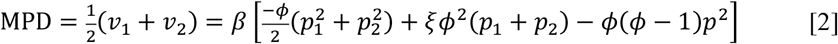

This expression follows from Eq. 1 and is a quadratic function of two variables, which from geometry represents a conic section (e.g., hyperbola, ellipse, or parabola) in the (*p*_1_, *p*_2_) plane. The general form of the quadratic equation for conic sections is typically written as

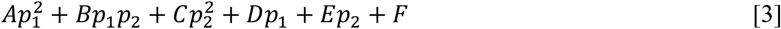

Comparing Eq. 2 and 3 allows for identification of the *A* through *F* coefficients (Table 1). Because *A*=*C* and *D*=*E* in this case, the centered, matrix form of the equation is

**Table 1.**
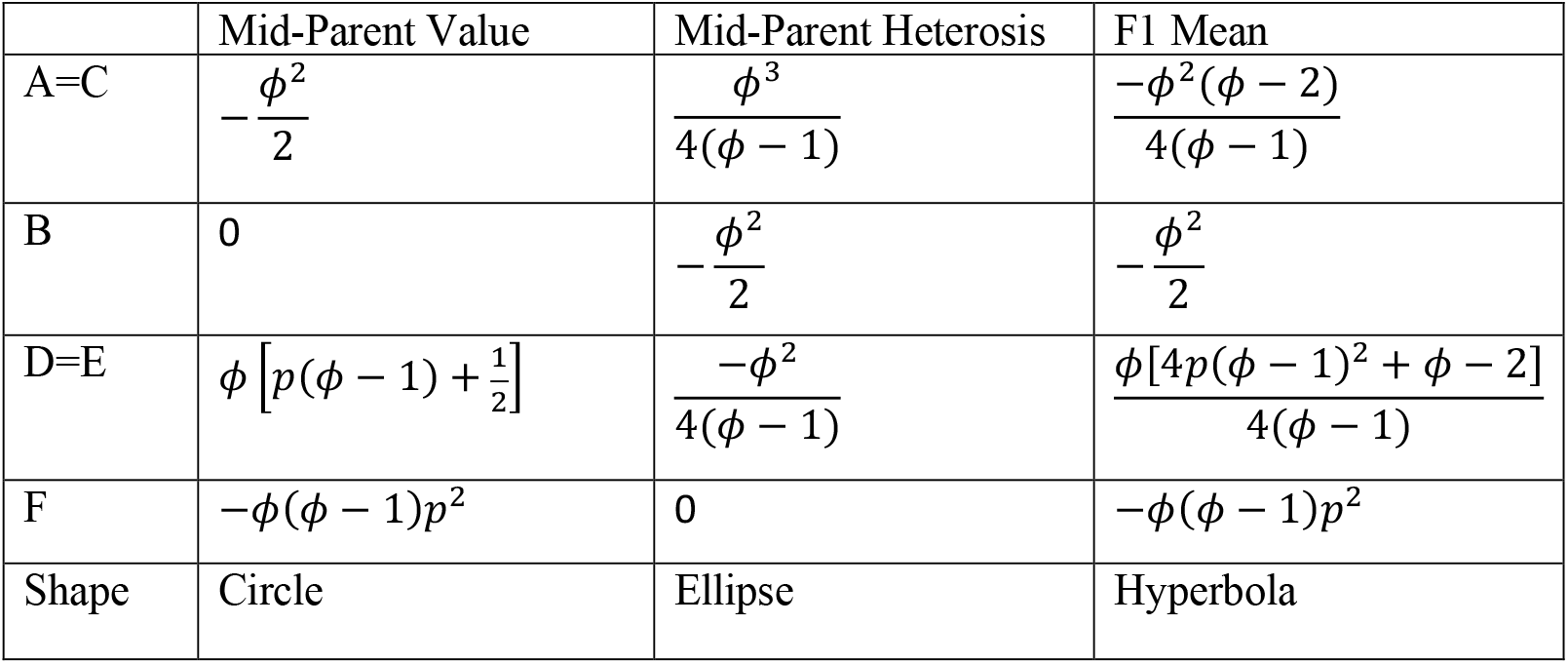
Parameters of the quadratic equation for dominance values,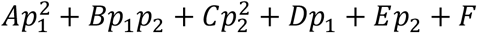.

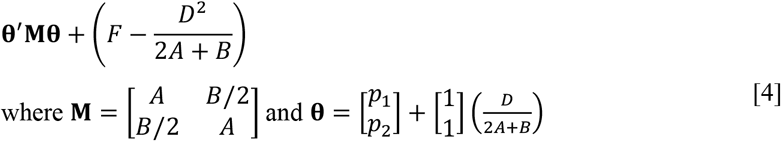

The sign of the determinant, |**M**| = *A*^2^ − *B*^2^/4, determines whether the conic section is a hyperbola (negative), ellipse (positive), or parabola (zero). The level curves of MPD are circles centered on (½, ½) when *p* = ½, but in general the center depends on *p* (Fig. 2A).

**Figure 2.**
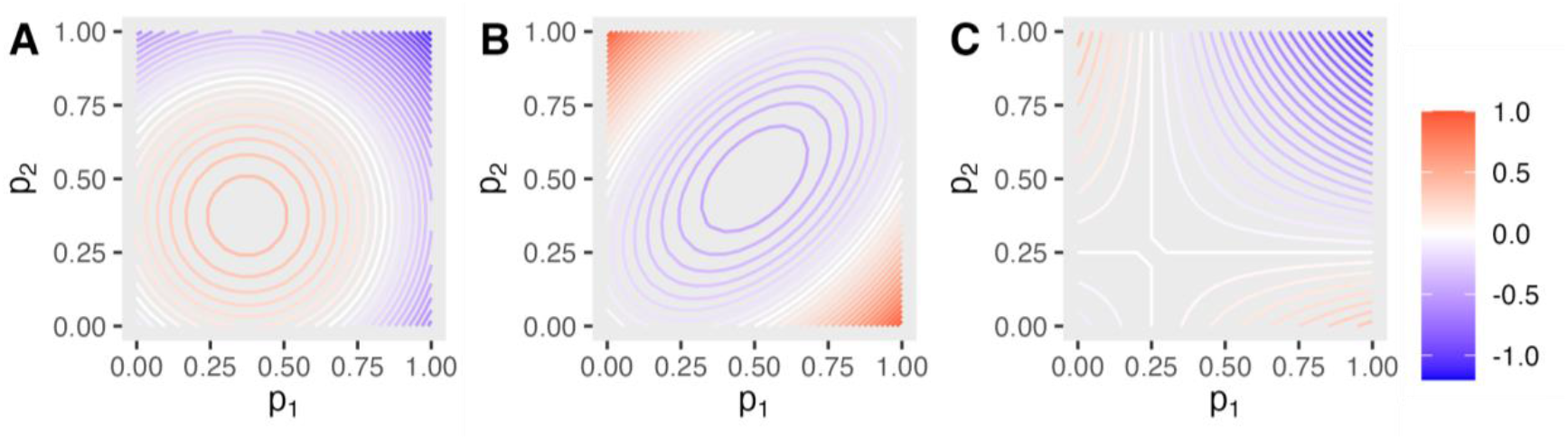
Level curves for mid-parent dominance value (A), mid-parent heterosis (B), and their sum (C), in diploids. The population frequency is ¼.

The solution for MPH was determined by first deriving the F1 mean under a random bivalents model of meiosis, which is mathematically equivalent to random chromosome segregation (Gallais 2003). Gametic dosage *γ* follows the hypergeometric distribution, which describes the probability of choosing *γ* copies of the allele in *ϕ*/2 draws without replacement from a parent with *ϕ* chromosomes and zygote dosage *z*. The first and second moments of this distribution are (Ibe 2014)

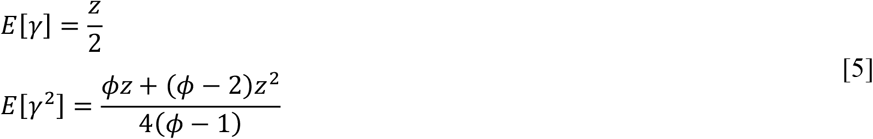

Allele dosage *z*_12_ of the F1 progeny from parents 1 and 2 is the sum of random variables *γ*_1_ and *γ*_2_, and the moments follow from Eq. 5:

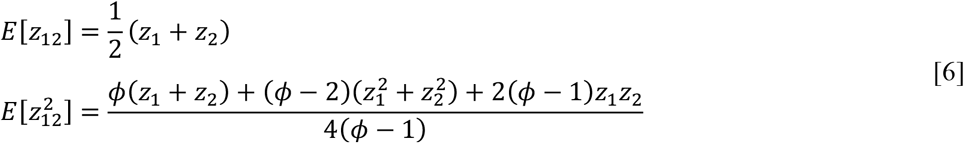

From Eq. 1, the expected dominance value of the F1 progeny equals *E*[*Q*_12_]*β*, where

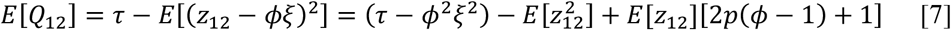

The expression *τ* − *ϕ*^2^*ξ*^2^ simplifies to conic parameter *F* in Table 1. The other conic parameters follow by substituting Eq. 6 into Eq. 7 and recognizing that *z*_*i*_ = *ϕp*_*i*_.

The solution for MPH follows by subtracting the conic parameters for MPD from the conic parameters for the F1 mean. Whereas MPD depends on allele frequency, MPH does not: it has elliptical level curves at a 45° angle to the coordinate axes that are always centered at (½, ½) (Fig. 2B). The eccentricity of the level curves, which is the ratio between the major and minor axes of the ellipse, increases with ploidy according to 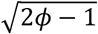 (derivation in File S1).

Viewed in three dimensions, MPD is a concave paraboloid, with its maximum value at the center point. MPH is a convex paraboloid, with its lowest value at the center, and its highest value when the two parents are homozygous for different alleles. Their sum is a hyperbolic paraboloid, also known as a saddle surface, which increases with distance from the saddle point in some directions but decreases in other directions (Fig. 2C). The hyperbola asymptotes are only perpendicular for diploids; the area of the plane where the dominance contribution is positive decreases with ploidy (Fig. S1 in File S2).

### Maximizing genetic merit

The proposed objective for OMA is to maximize the F1 progeny mean:

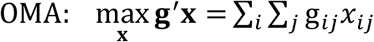

*g*_*ij*_ is the mean genotypic value (relative to the population mean) of progeny from female parent *i* and male parent *j*, and *x*_*ij*_ denotes the allocation for this mating. Unless otherwise indicated, the sum index is from 1 to *n* parents. The *n*^2^ x 1 column vectors **g** and **x** can be viewed as vectorized *n* x *n* matrices. The mate allocation variables are proportions that sum to 1. The total contribution (*y*) of an individual is the average of its female (*y*_*f*_) and male (*y*_*m*_) contributions, allowing for hermaphroditic species. (Some authors define *y* as a sum instead of average, which does not affect the results.) The sex-specific contributions are computed by summing over the other index of the mate allocation variables:

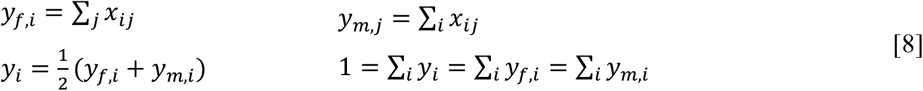

In the previous section, the F1 progeny mean was expressed as MPA+MPD+MPH. Alternatively, it can be expressed as the sum of general and specific combining abilities (GCA and SCA, respectively): *g*_*ij*_ = GCA_*i*_ + GCA_*j*_ + SCA_*ij*_. Breeding value (BV) is twice the GCA at HWE. In the diploid single locus model, breeding value equals additive value, which identifies SCA as MPD+MPH. For polyploids, BV includes a proportion 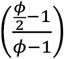 of the dominance value (Gallais 2003). If SCA is neglected, the OMA objective reduces to maximizing the average breeding value of the parents, weighted by their contributions (derivation in File S1), which is the typical objective for OCS:

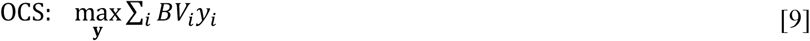

### Inbreeding control

The inbreeding coefficient *F*_*t*_ in generation *t* is the probability that two genes at one locus are identical by descent (IBD). For polyploids, the genes are randomly sampled without replacement (Gallais 2003). Using the law of total probability, we can condition on whether the two genes are derived from the same gene in the previous generation, which occurs with probability Δ*F*:

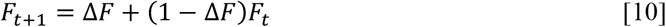

Eq. 10 is a recurrence equation between the inbreeding coefficients from adjacent generations, and Δ*F* is the inbreeding rate (Falconer and Mackay 1996). Iterating one more generation produces

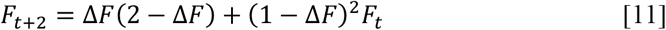

and an explicit time equation follows from the formula for a finite geometric series:

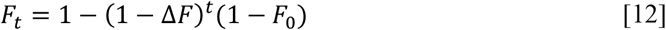

In the OCS literature, the constraint on inbreeding rate is implemented using the quadratic form **y**^**′**^**ky**, which is called group coancestry, or kinship (**y** is the vector of contributions, and **K** is the kinship matrix). To derive the formula for any ploidy, let *F*_*ij*_ denote the inbreeding coefficient of an individual in generation *t*+1 with parents *i* and *j*, which have inbreeding coefficients *F*_*i*_ and *F*_*j*_, respectively. Under random chromosome segregation, the relationship between inbreeding and kinship across one generation is (Kerr et al. 2012)

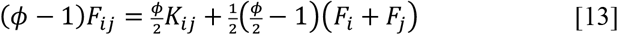

The average inbreeding coefficient of generation *t*+1 is the average over all possible matings, weighted by their allocation:

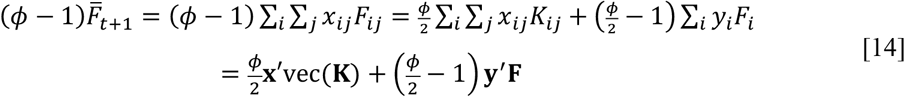

The second equality in Eq. 14 follows from the relationship between mate allocations and individual contributions (Eq. 8), and the third equality introduces vector notation. The symbol 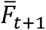 has the same meaning as *F*_*t*+1_ in Eq. 10, but the overbar (for average) is introduced to differentiate the two kinds of subscripts: time vs. individuals. Under random mating, mate allocation is the product of the female and male contributions (*x*_*ij*_ = *y*_*f,i*_*y*_*m,j*_), which leads to

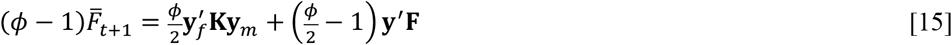

For hermaphroditic individuals with equal female and male contributions (**y**_*f*_ = **y**_*m*_ = **y**), the expression further simplifies to

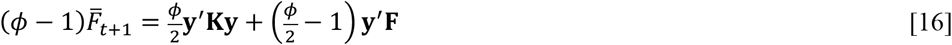

Analogous to Eq. 13, the relationship between inbreeding and kinship between generations *t*+2 and *t*+1 is

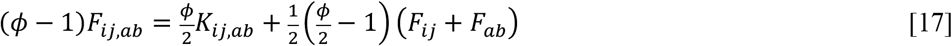

*F*_*ij,ab*_ denotes the inbreeding coefficient of an individual in generation *t*+2 with female parent *ij* in generation *t*+1 and maternal grandparents *i* and *j* in generation *t*. The male parent is *ab*, with paternal grandparents *a* and *b*. Using Eq. 13 and that parental kinship is the average of four grandparental kinships, inbreeding in *t*+2 can be related to the parameters of generation *t*:

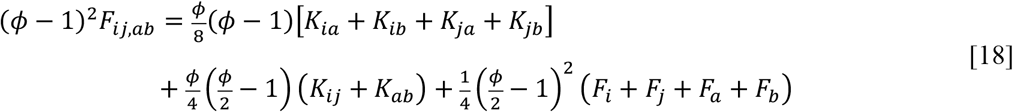

The average inbreeding coefficient of generation *t*+2 is the average over matings in *t*+1, weighted by their allocation. Since mate allocation in generation *t*+1 is not yet determined, random mating is assumed as a conservative bound, which leads to the following result (derivation in File S1):

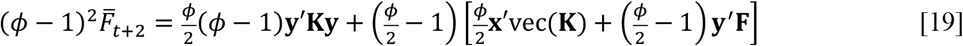

## METHODS

### Convex Optimization

The convex OMA problem is

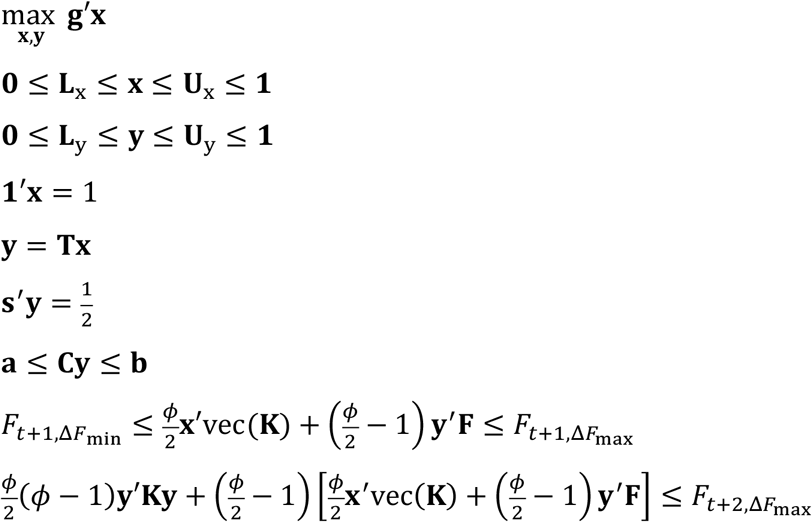

The genetic merit **g** for each mating was the sum of MPA, MPD, and MPH across all markers. The first two constraints represent lower (**L**) and upper (**U**) bounds on the mate allocation (**x**) and contribution (**y**) variables. The incidence matrix **T** in constraint **y** = **Tx** captures the relationship between the variables in Eq. 8. Constraint 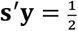 is only used with separate sexes, and the elements of **s** are 0 for females and 1 for males to normalize the sex-specific contributions. Equation **a** ≤ **Cy** ≤ **b** represents optional linear constraints on the contributions; an example is given for the potato dataset. The constraint involving *F*_*t*+1_ combines Eq. 10 and 16, putting lower and upper bounds on the inbreeding coefficient of the progeny based on lower and upper bounds on the inbreeding rate: Δ*F*_min_ ≤ Δ*F* ≤ Δ*F*_max_. The last constraint combines Eq. 11 and 19, putting an upper bound on the inbreeding coefficient of the grand-progeny; **K** must be positive semidefinite for convexity.

If ***δ*** denotes the vector of breeding values, the convex OCS problem is

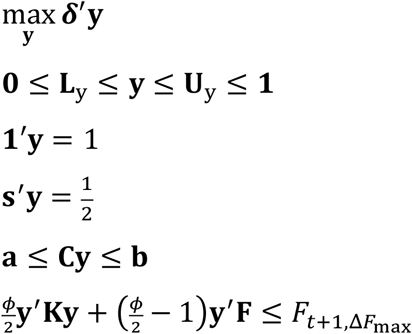

The most appropriate inbreeding constraint, Eq. 15, is not convex, so the hermaphroditic form, Eq. 16, is used instead. This is the traditional OCS approach (Woolliams et al. 2015), but as Eq. 19 illustrates, it is a tighter bound than strictly needed for diploid species with separate sexes.

R package COMA (https://github.com/jendelman/COMA) uses R package CVXR (Fu et al. 2020) to solve convex formulations of OCS and OMA (File S3). CVXR transforms high-level descriptions of convex optimization problems into the format required of convex solvers. The group kinship constraints identify OCS and OMA as second-order cone programs, and the ECOS solver that comes with CVXR is the default in COMA. During the breeding program simulation described below, ECOS sometimes returned with status “optimal_inaccurate” instead of the desired result “optimal;” this issue was resolved by using the commercial solver MOSEK, which must be installed separately (MOSEK ApS 2024).

From the optimal solution, the average inbreeding coefficients of the progeny and grand-progeny are calculated according to Eq. 16 and 19, respectively. The average inbreeding rates for the progeny, Δ*F*_1_, and grand-progeny, Δ*F*_2_, are then computed by inverting Eq. 12:

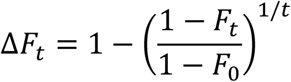

### Potato example

Marker effects were computed from the 2015–2019 data of the R/StageWise package (Endelman 2023), which contained three traits—total yield, fry color, and vine maturity—and 12K SNP markers for 700 tetraploid clones. Two-stage analysis of the multi-year trial followed Vignette 3 of the package (File S5). Genotype means (BLUEs) were estimated per environment (*StageWise::Stage1*) and then used as the response variable in the multi-environment mixed model (*StageWise::Stage2*), using ASReml-R v4.2 (Butler et al. 2023) to estimate variance-covariance (var-cov) parameters. The mixed model includes additive *u*_*ik*_ and dominance *v*_*ik*_ values for trait *k* of clone *i*, the fixed effect *E*_*jk*_ of trait *k* in environment *j*, a random effect *s*_*ijk*_ for the Stage 1 error, and the residual *gE*_*ijk*_ is the genotype x environment interaction:

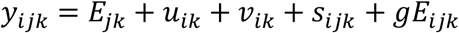

The Stage 1 error is multivariate normal, with var-cov equal to the direct-sum of the var-cov matrices of the multi-trait BLUEs from Stage 1. The genetic and residual effects were also multivariate normal with separable covariance for clones vs. traits:

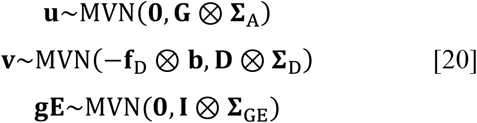

The trait var-cov matrices (**Σ**) were unstructured (i.e., full rank). The additive **G** and dominance **D** genomic relationship matrices follow the multi-locus extension of Eq. 1, using allele frequency *p*_*k*_ at locus *k* for the current population (Endelman 2023):

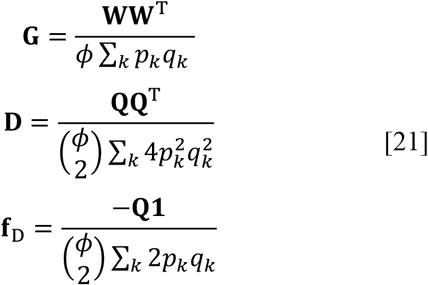

The **W** and **Q** matrices contain the additive and dominance regressors, respectively (Eq. 1). The elements of covariate **f**_D_ are genomic inbreeding coefficients, and the corresponding vector of regression coefficients **b** (Eq. 20), one for each trait, represents inbreeding depression, aka baseline heterosis (Labroo et al. 2021).

Empirical BLUPs for marker effects were based on a multi-trait index using R/StageWise, with the contribution of inbreeding depression included in the dominance effects (Endelman 2023). Yield and fry color have similar importance as breeding objectives, while later maturity is undesirable. Initially, a restricted index was calculated, with equal merit for yield and fry color and zero gain for maturity (File S5). Due to the unfavorable correlation between yield and late maturity, the yield response for this index was only 65% of the response for fry color. As an alternative, a desired gains index was calculated (Pesek and Baker 1969), based on equal gains for yield and fry color and zero gain for maturity. The restricted index methodology was described in Endelman (2023); from Eq. 24 of that publication, if ***δ*** denotes the desired gain vector, then the index coefficients of the multi-trait BLUPs are **c** = **B**^−1^***δ***, where **B**^−1^ is the inverse var-cov matrix of the BLUPs. The realized response is proportional to the desired gains, and from Eq. 25 of Endelman (2023), the proportionality constant is 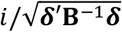 for selection intensity *i*. Imposing equal gains for yield and fry color only reduced total merit by 1% (File S5), and given the uncertainty of the true merit coefficients, this index was selected.

Estimates of dominance variance and inbreeding depression for the multi-trait index were expressed relative to the additive variance and standard deviation, respectively (File S5, Table S1 in File S2). If **v** denotes the multi-trait vector of dominance values (Eq. 20), the vector of dominance index values is (**I⨂c′**)**v**, with mean **μ** = −(**c′b**)**f**_D_ and variance **V** = (**c′Σ**_D_**c**)**D**. The dominance variance of the population is therefore 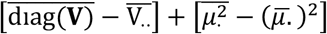 (Legarra 2016; Endelman 2023), and the additive variance was computed analogously.

The PVY resistance data are based on KASP marker snpST00073 for Ry_adg_ (Kante et al. 2021), which is co-dominant but not reliable at differentiating the three possible heterozygotes in tetraploid potato. Based on the low frequency of Ry_adg_ in our germplasm, a frequency of ¼ (i.e., single copy) was assumed for clones testing positive for the marker.

### Breeding simulation

A mass selection program was simulated with AlphaSimR (Gaynor et al. 2021), using a constant population size of 500 hermaphroditic individuals and non-overlapping generations. R/SimPlus (https://github.com/jendelman/SimPlus) was created to standardize breeding and analysis modules across the treatments (File S4).

For each of the ten replicate simulations per ploidy (2x and 4x), a founder population (N=500) in mutation-drift equilibrium was created using the coalescent, with mutation rate 10^−8^, effective population size 100, and a genome of ten 1 Morgan chromosomes. These parameters generated approximately 15K segregating sites in the 2x founders and 30K in the 4x founders. Founder haplotypes were tracked across generations using setTrackRec=TRUE.

AlphaSimR uses the (*a,d*) parameterization for QTL effects, where 2*a* is the difference between the homozygotes and *a*+*d* is the difference between the heterozygote and recessive homozygote (Falconer and Mackay 1996). An “ADG” trait was simulated with the following settings: nQtlPerChr=1000, mean=0, var=1, varDD=0.5, meanDD=0.5, useVarA=FALSE. This implies a normal distribution for the dominance degree (DD) parameter, *d*/|*a*|, with mean 0.5, variance 0.5, and effects scaled to create unit genetic variance in the founders. The average ratio of dominance to additive variance in the founder population was 0.12 for diploids and 0.25 for tetraploids, compared to 0.38 for the potato dataset (Table S1 in File S2). The average baseline heterosis, scaled by the additive genetic standard deviation, was 4.3 and 7.3 for the diploid and tetraploid simulations, respectively, compared to 7.1 for the potato dataset. Non-genetic variance (parameter varE) was fixed at 2 throughout the simulation.

10 generations of phenotypic selection were simulated using OCS at 0.5% inbreeding (pedigree kinship), followed by 25 generations of genomic selection using different methods. (Generation 0 designates the beginning of GS.) Each GS cycle consisted of (1) phenotyping the candidates, (2) calculating marker effects, (3) developing the mating plan by OCS or OMA, (4) and generating new progeny. To reduce computing time, the training population for predicting marker effects was limited to phenotype and genotype data for the selection candidates and two previous generations. The phenotypes and complete genotype data (including QTL) were written to file and analyzed with R/StageWise to predict the additive and dominance marker effects.

Selection accuracy was estimated each generation based on the correlation between the true and predicted means for 100 randomly selected matings, using 50 progeny per mating. The OCS prediction is based on the sum of the GCAs, while OMA also includes SCA.

Due to the large number of mate allocation variables (125,250) for 500 candidates, the candidate pool was reduced to 200 individuals by OCS before applying OMA. A lower bound *N* on the number of selected parents can be achieved using an upper limit 1/*N* on the parental contributions. When there was no solution at the target rate (i.e., problem infeasible), the upper bound was increased in increments of 0.5% (software parameter *dF*.*step*).

Three different kinship models were tested and named based on the corresponding relationship matrix. (Kinship equals relationship divided by ploidy.) Method A is pedigree kinship based on the probability of IBD and was computed with R/AGHmatrix (Amadeu et al. 2016). Method G_IBD_ is genomic kinship and extends method A to capture Mendelian sampling; it is equivalent to an allele matching coefficient using founder alleles, which were obtained in AlphaSimR with *pullIbdHaplo*. Method G is Method 1 of VanRaden (2008), using founder allele frequencies as the reference population (Eq. 21). Computation of the **G**_IBD_ and **G** matrices was implemented as function *G_mat* in R/polyBreedR (Endelman et al. 2024). Inbreeding coefficients were computed from the diagonals of the kinship matrix according to *f*_*A,i*_ = (*ϕK*_*ii*_ − 1)/(*ϕ* − 1) (Gallais 2003).

The mating plan from COMA was realized in AlphaSimR using *SimPlus::sim_mate*, which generates progeny according to the vector of mate allocations **x**, interpreted as a discrete probability distribution. The mating plan under OCS was a diallel with mate allocation equal to the product of the parental contributions.

## RESULTS

### Constraints and kinship

COMA limits inbreeding in the progeny under the optimized mating plan, and in the grand-progeny assuming random mating of the progeny. The grand-progeny constraint is needed to maintain the target inbreeding rate; without it, the optimal solution would be 100% allocation to the mating with the highest merit for unrelated diploid parents. The grand-progeny constraint is quadratic and therefore, to remain convex, is only an upper bound (as with OCS). The progeny constraint is a linear inequality, which allows for both upper and lower bounds on the inbreeding rate. Initially, only an upper bound was used, based on the hypothesis that if the solution with maximum genetic merit involved less inbreeding, this would be advantageous for long-term gain.

But this hypothesis was contradicted in a simulated genomic selection program. Comparing COMA selection with upper bound (UB) vs. equality (EQ) constraints at 0.5% inbreeding, genetic gain with the UB strategy was slightly higher after one generation (Fig. 3A, Fig. S2 in File S2). This is expected from the principles of mathematical optimization: since all possible mate allocations under EQ are also allowed under UB, the optimal UB solution cannot be lower and will typically be higher. But when COMA was applied in generation 1, there was more gain with EQ than UB in generation 2; this does not violate any mathematical principle because the candidates are different. When inbreeding was measured by pedigree IBD kinship (A), the long-term gain advantage of EQ over UB was small but statistically significant at the 0.05 level (Fig. 3A, Fig. S2 in File S2). For genomic IBD kinship (G_IBD_), the rate of genetic gain with EQ was much higher.

**Figure 3.**
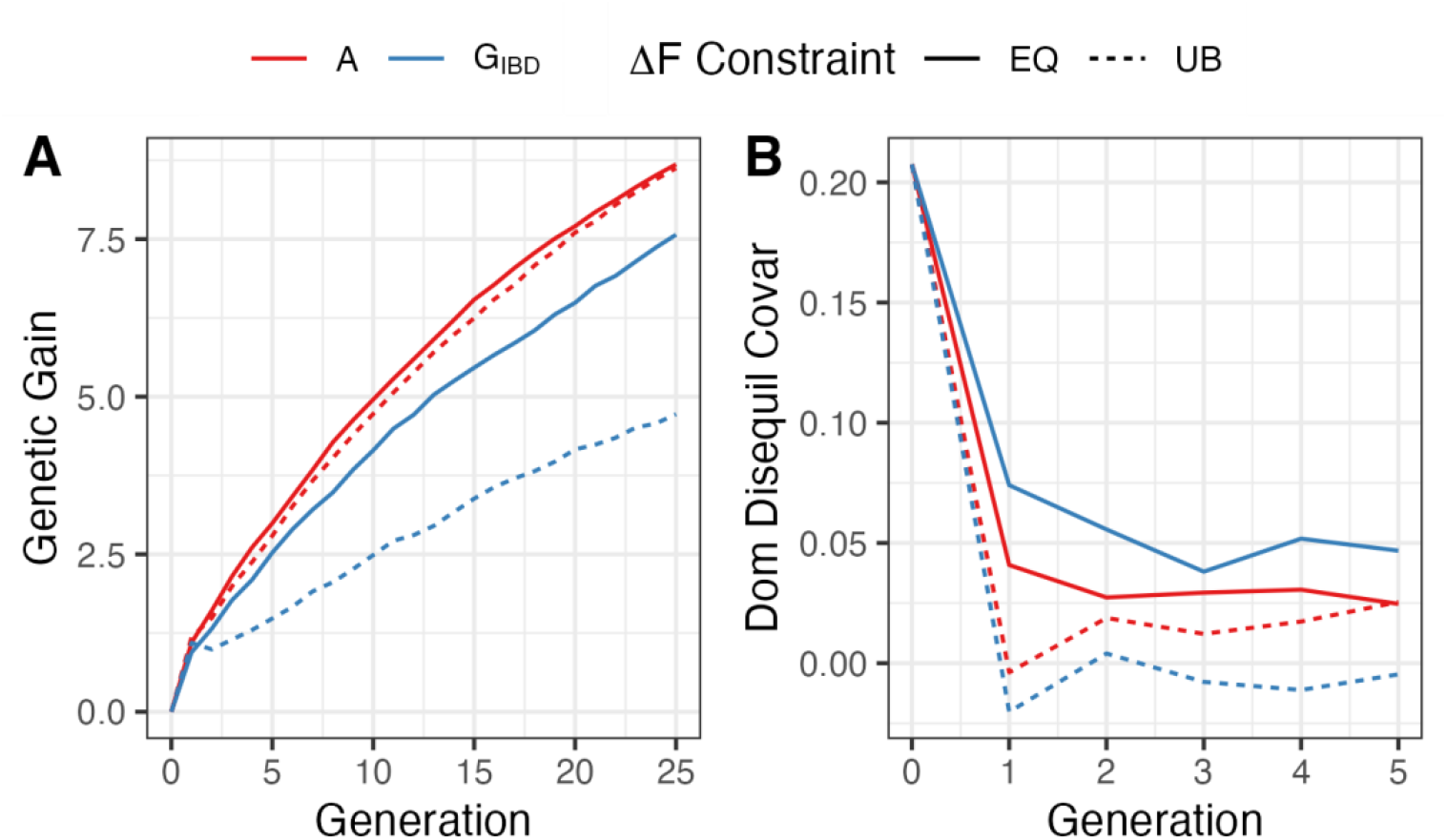
Diploid simulation results for genetic gain and dominance disequilibrium covariance, using COMA with either equality (EQ) or upper bound (UB) constraints for the target inbreeding rate of 0.5%.

The trends for inbreeding coefficient and genetic variance help explain the results for genetic gain. Under the EQ constraint, the target inbreeding rate of 0.5% was achieved for both pedigree and genomic kinship (Fig. 4). But with only an UB constraint, the inbreeding coefficient decreased in the first generation. With pedigree kinship, the inbreeding rate returned to the target value within a few generations, but with genomic kinship, the rate only rebounded to ∼0.1%. When partitioning dominance genetic variance into genic variance (which depends only on allele frequencies) and disequilibrium covariance (DCV), the former was unchanged in the first generation, but the latter decreased substantially (Fig. 3B, Table 2). This was observed for both EQ and UB constraints, but whereas some DCV remained under EQ, it was exhausted under UB.

**Table 2.** ANOVA p-values for the effect of kinship (A vs. G_IBD_) and inbreeding constraint (EQ vs. UB) on the components of genetic variance after one generation of COMA selection.

| Effect | Df | Additive<br>Genic<br>Variance | Additive<br>Disequilibrium<br>Covariance | Dominance<br>Genic<br>Variance | Dominance<br>Disequilibrium<br>Covariance |
| --- | --- | --- | --- | --- | --- |
| Replicate | 9 | $< 2 \times 10^{-16}$ | 0.03 | $< 2 \times 10^{-16}$ | 0.02 |
| Constraint | 1 | 0.45 | 0.85 | 0.79 | $4 \times 10^{-8}$ |
| Kinship | 1 | 0.0001 | 0.09 | 0.55 | 0.37 |
| Constraint:Kinship | 1 | 0.82 | 0.14 | 0.96 | 0.01 |
| Residuals | 27 |  |  |  |  |

**Figure 4.**
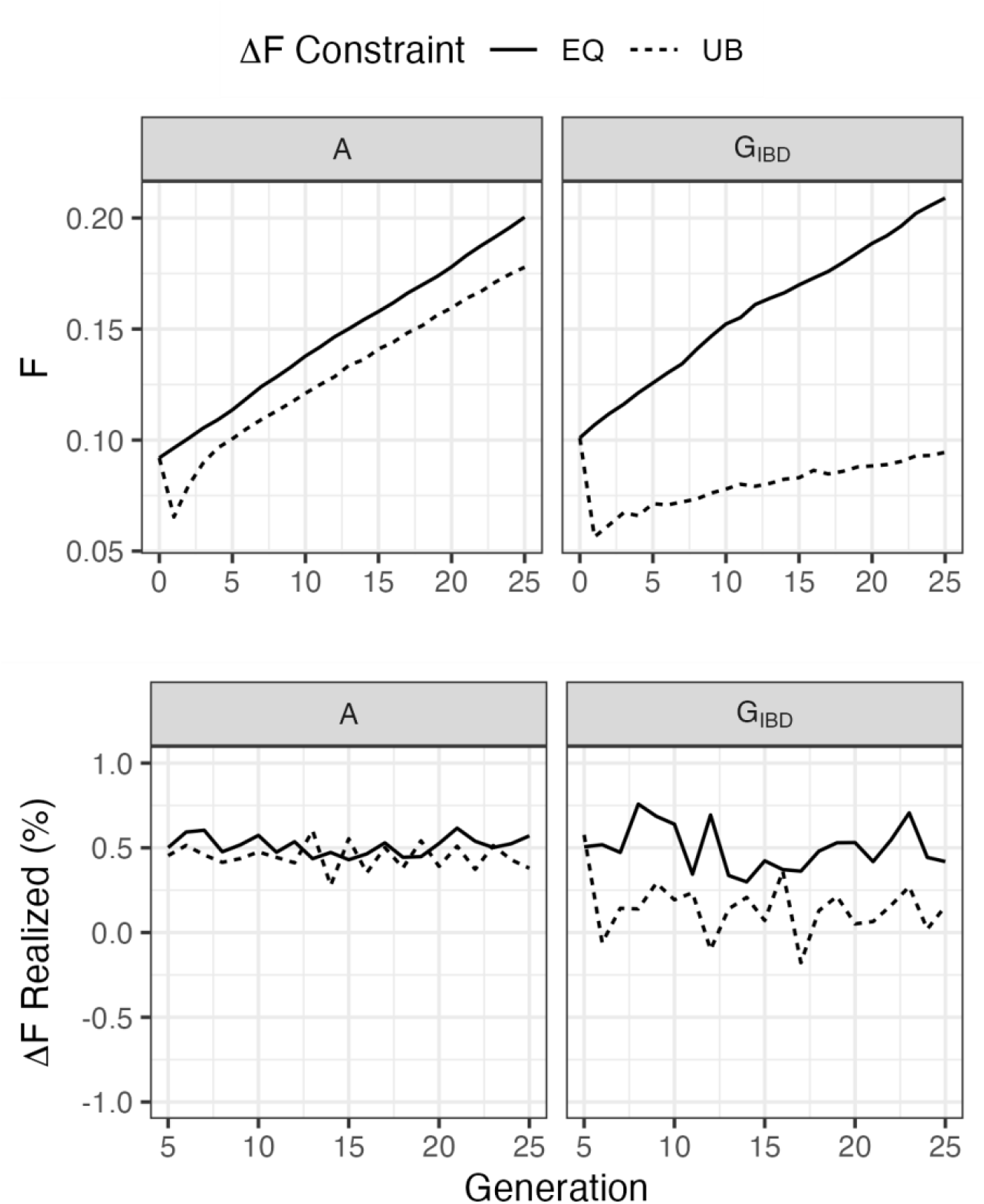
Realized inbreeding coefficients (top) and rates (bottom), using COMA with either equality (EQ) or upper bound (UB) constraints. The target inbreeding rate was 0.5%, based either on pedigree (A) or genomic (G_IBD_) kinship.

When the two kinship methods were compared under EQ constraints, pedigree kinship resulted in significantly higher gain for both the diploid (Fig. 3A, Fig. S2 in File S2) and tetraploid (Fig. S3 in File S2) simulations.

### Comparison with OCS

The COMA software also contains an implementation of OCS, and the two methods were compared at 0.5% and 1.0% inbreeding using pedigree kinship. Both selection methods achieved the target inbreeding rates at both ploidy levels (Fig. S4 in File S2). The number of selected parents was not significantly different between OMA and OCS (Fig. 5) and was higher than the effective population size, 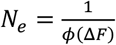, for ploidy *ϕ*. At 0.5% inbreeding, 111 diploid and 74 tetraploid parents were selected on average. At 1.0% inbreeding, those numbers decreased to 72 and 36 parents, respectively. The number of matings with OMA was much lower than with OCS, which involved hundreds of random matings based on the optimum contributions. At 1.0% inbreeding, the average number of matings with OMA was only 60 for diploids and 11 for tetraploids.

**Figure 5.**
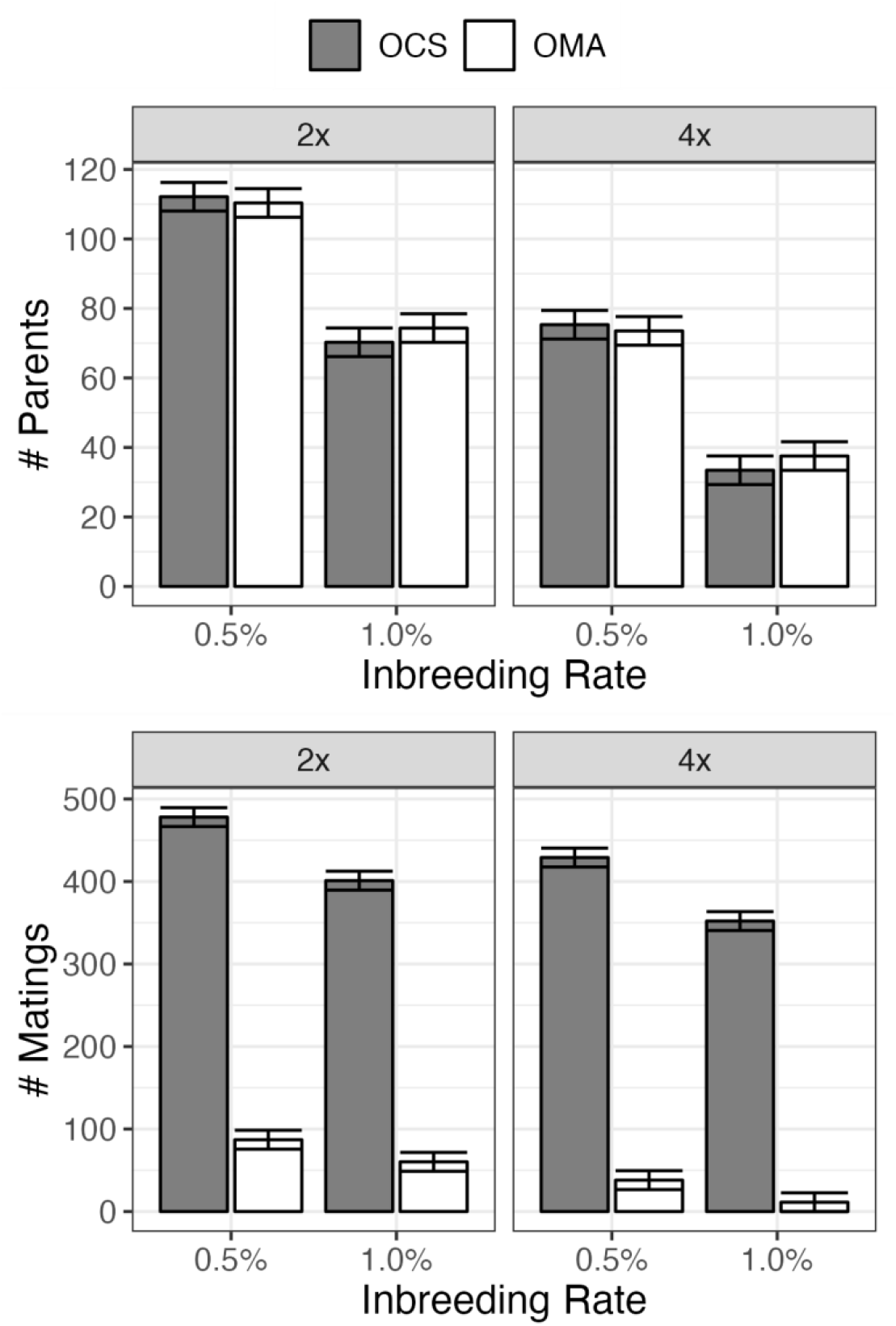
Number of parents and matings for generations 5–25, using either OCS or OMA and pedigree kinship at 0.5% or 1.0% inbreeding. The error bars are the 95% CI based on 10 independent simulations.

OMA achieved the same or higher gain as OCS and was significantly more efficient at converting genic variance into gain, but the difference was small (Fig. 6). At 0.5% inbreeding, the gain advantage of OMA persisted for all 25 generations in tetraploids; for diploids, the gain advantage disappeared by 25 generations, but the genic variance with OMA was significantly higher. Similar results were observed at 1.0% inbreeding (Fig. S5 in File S2).

**Figure 6.**
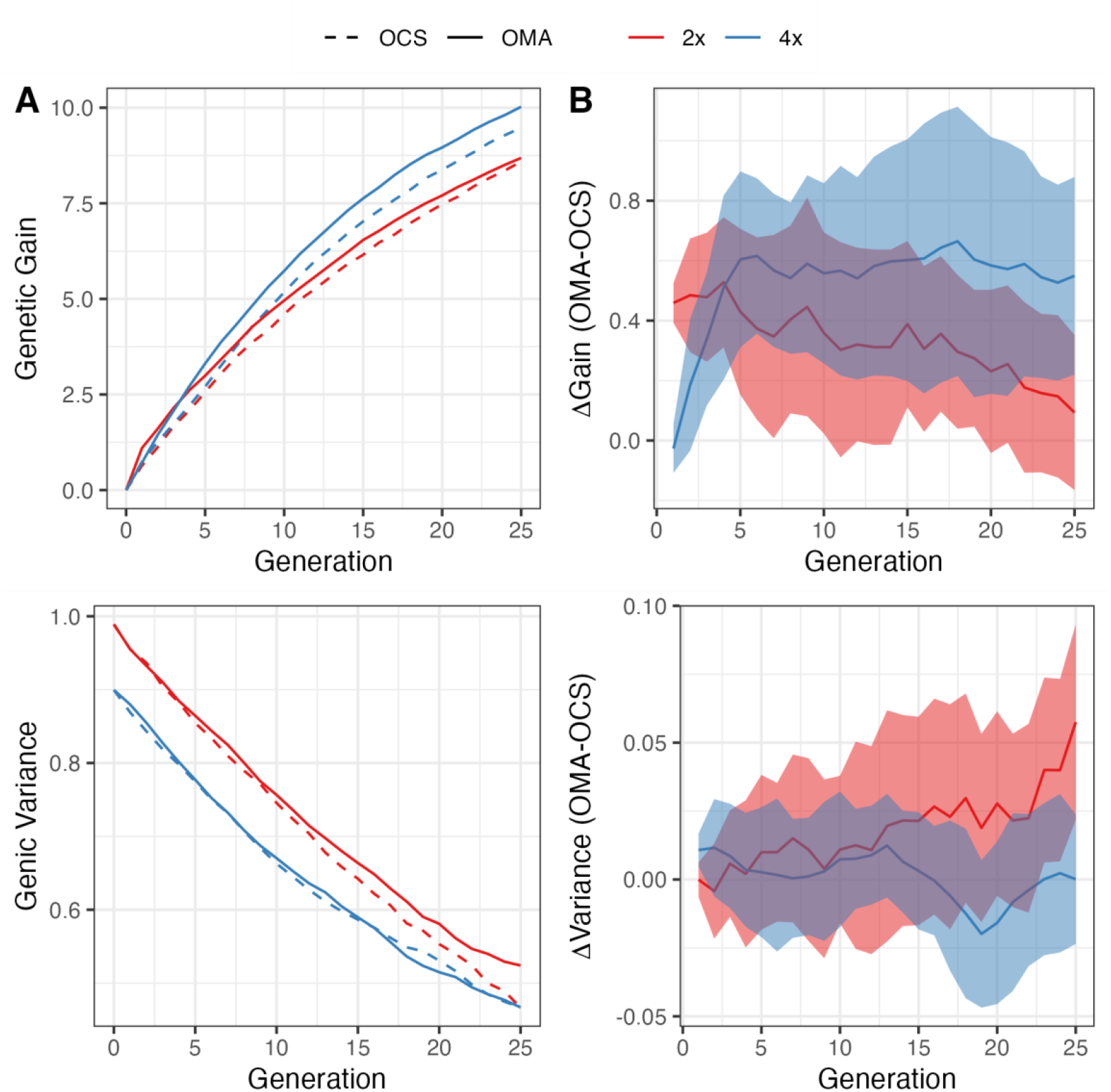
(A) Simulation results for genetic gain and genic variance using OCS vs. OMA at 0.5% inbreeding with pedigree kinship. (B) The solid line is the mean difference across the 10 replicate simulations, and the ribbon is the 95% CI from a paired *t*-test each generation.

Because OMA includes specific combining ability (SCA), the accuracy for genomic prediction of mate performance (GPMP) was consistently 0.2–0.3 higher than with OCS (Fig. 7). GPMP accuracy was not significantly different between the 2x and 4x simulations with OMA, but with OCS the 4x accuracy was higher (Fig. 7). The difference can be attributed to a smaller SCA variance in the 4x simulation, which occurred because (1) the SCA variance at HWE is *V*_*D*_/4 for 2x but *V*_*D*_/6 for 4x (Gallais 2003); and (2) there was only half as much as *V*_*D*_ (relative to *V*_*A*_) in the 4x founders (Table S1 in File S2).

**Figure 7.**
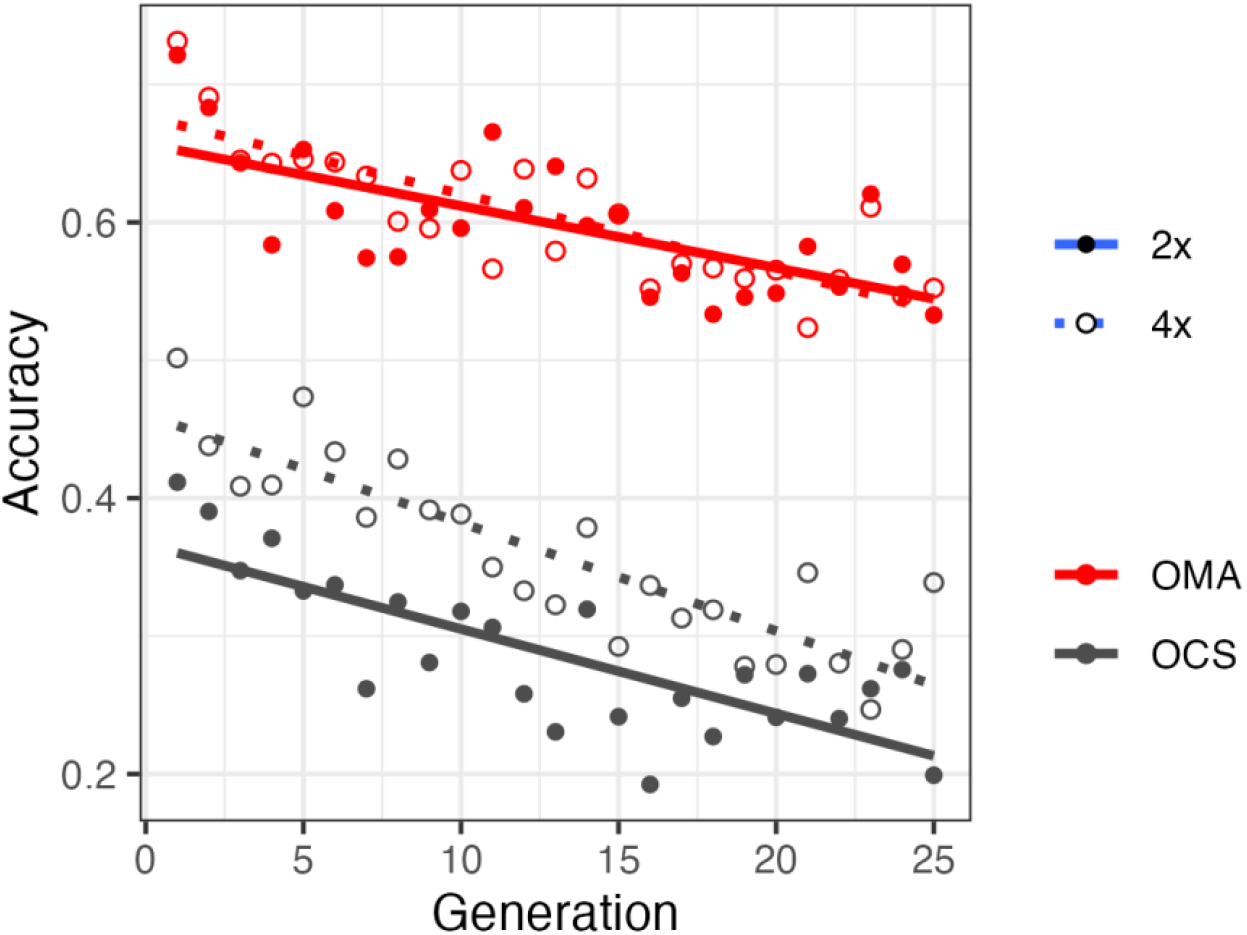
Accuracy of OMA and OCS for predicting mate performance, based on 100 random matings and 50 progeny per mating. Each point is the average of 10 replicate simulations.

### Potato example

The COMA tutorial uses data from the University of Wisconsin potato breeding program (File S5). The parental candidates are 170 autotetraploid clones, with 12K SNP array markers and a pedigree kinship matrix. The inbreeding coefficients ranged from 0.00 to 0.09 (median 0.02), and kinship ranged from 0.01 to 0.19 (median 0.09). Additive and dominance marker effects were predicted by BLUP using a multi-trait selection index and training population of 700 clones.

As mentioned above, COMA can impose lower (LB) and upper bounds (UB) on the inbreeding rate for the next generation, Δ*F*_1_. The UB is also applied to the average inbreeding rate over two generations, Δ*F*_2_, assuming random mating of the progeny. There was no feasible solution with an UB of 0.5%, which reflects the relatedness of the candidates. The smallest feasible UB was 0.8% when no LB was imposed, resulting in Δ*F*_1_ = −0.8% and a predicted progeny mean of 1.22*σ*_*A*_. The tendency toward outbreeding is reminiscent of the simulation results, which raises the concern that the population could get trapped in a local optimum. With a LB of 0.5% for Δ*F*_1_, the smallest feasible value for Δ*F*_2_ was 1.0%, and the predicted merit was 1.23*σ*_*A*_. As seen in the simulation, COMA solutions are sparse: of the 903 possible matings (excluding selfs and reciprocals) for the 43 selected parents, only 43 matings were needed to achieve optimal gain.

The lowest feasible inbreeding rate with OCS was 1.5%. Although the software reports the predicted merit of the optimum solution as 1.22*σ*_*A*_, this neglects the contribution of SCA. When SCA was included, the predicted merit reduced to 1.04*σ*_*A*_, which is lower than the OMA solution despite the higher inbreeding rate.

The COMA tutorial shows how additional constraints can be incorporated. Besides the three quantitative traits in the selection index (yield, fry color, and maturity), resistance to potato virus Y (PVY) is an important qualitative trait, which was scored with a genetic marker. For the original COMA solution with (Δ*F*_1_, Δ*F*_2_) = (0.5%, 1.0%), 19 of the 43 matings had a resistant parent, with an expected allele frequency of 8.6% in the next generation assuming single-copy dosage in the parents (Table 3). One selection strategy would be to require a resistant parent in every mating, which is achieved by setting the maximum allocation to 0 for matings that do not meet this criterion. This increased the resistant allele frequency to 12.5% in the progeny, with the tradeoff being a reduction in predicted merit from 1.23*σ*_*A*_ to 1.02*σ*_*A*_. Alternatively, the allele frequency can be constrained directly with an inequality of the form **s**^**′**^**y** ≥ *p*, where **s** is the vector of allele frequencies in the parents (i.e., ¼ for single-copy resistant and 0 for susceptible) and *p* is the target allele frequency. Using this approach to achieve 12.5% R gene frequency, the predicted merit was slightly higher, at 1.07*σ*_*A*_.

**Table 3.** The influence of constraints for potato virus Y resistance (PVYR) on the COMA solution with Δ*F*1 = 0.5% and Δ*F*2 = 1.0%.

| PVYR Constraint | Response ($\sigma_A$) | # Parents | # Matings | PVYR freq |
| --- | --- | --- | --- | --- |
| None | 1.23 | 43 | 43 | 8.6% |
| At least one R parent<br>per mating | 1.02 | 42 | 40 | 12.5% |
| PVYR freq $\geq 12.5\%$ | 1.07 | 43 | 42 | 12.5% |

## DISCUSSION

Two types of inbreeding coefficient were used in this study. The first, denoted *f*_D_, was introduced as a covariate in the genomic prediction model to account for inbreeding depression. From the theory of directional dominance (Endelman 2023), *f*_*D*_ is proportional to the dominance regressor *Q*, averaged over loci (Eq. 1 & 21). Endelman (2023) observed in the potato dataset that the population mean 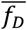 was numerically identical to 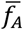, calculated from the diagonal elements of the additive genomic relationship matrix (**G**), although the individual values were not identical. The derivation of this mathematical equality is given in File S1:

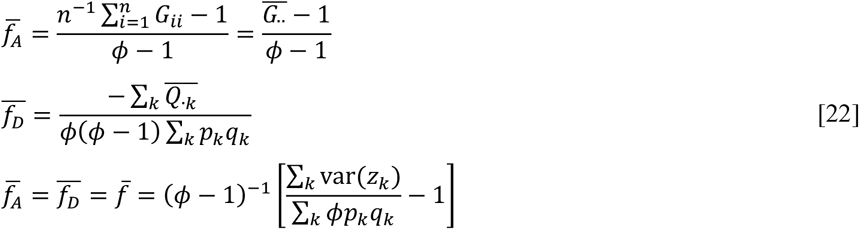

Eq. 22, which uses allele frequency *p*_*k*_ at locus *k* for the current population, indicates 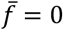 when the population variance of the allele dosages, var(*z*), equals the variance (*ϕpq*) of a binomial random variable with the same allele frequency. The maximum value of 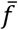 occurs when there are no heterozygotes, in which case var(*z*) = *ϕ*^2^*pq*, so 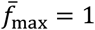. The minimum value of 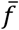 occurs when var(*z*) = 0, so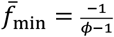. The limits of 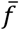 for diploids are [-1,1], consistent with its interpretation as a gametic correlation (Wright 1921; Powell et al. 2010; Endelman and Jannink 2012).

The second type of inbreeding coefficient, denoted *F*, was introduced to limit inbreeding during mate selection. For this application, *F* was defined as the probability of IBD, and both pedigree (A) and genomic (G_IBD_) kinship were effective at meeting the target inbreeding rate in the simulation. But genetic gain was higher using pedigree kinship (Fig. 3), and similar results with OCS have been reported (Meuwissen et al. 2020). This finding is consistent with theoretical and computational work showing the selective advantage of an individual under OCS is dominated by its Mendelian sampling term (Grundy et al. 1998; Avendaño et al. 2004). Because individuals with selective advantage from the same family typically have higher genomic kinship than the expected value from pedigree, there is less flexibility to increase gain by selecting multiple full-sibs (or half-sibs) under genomic control of inbreeding (Henryon et al. 2019).

What about using **G**, rather than **G**_IBD_, for genomic control of inbreeding? This is practically appealing because the latter is computationally intensive for diploids (Meuwissen and Goddard 2010; Whalen et al. 2018) and has not even been reported in autotetraploids; only one generation from multiple founders has been published (Zheng et al. 2021). (In the present study, tetraploid founder haplotypes were tracked during the meiosis simulation, not inferred.) But using **G** for inbreeding control is not recommended within the current framework. The constraint equations are based on the law of total probability (Eq. 10), while the inbreeding coefficient from **G** is a correlation (Eq. 22), not a probability. De Beukelaer et al. (2017) also highlighted this theoretical discrepancy and verified it by simulation, using the current population as the reference population. It is more common to use reference allele frequencies from the founder population, but this also failed under simulated mass selection with COMA: at a target inbreeding rate of 0.5%, the realized rate exceeded 5%, and genic variance was depleted too rapidly (Fig. S6 in File S2).

My initial hypothesis was that OMA would achieve higher gain than OCS because of its ability to predict SCA. The simulation confirmed that GPMP accuracy was 0.2–0.3 higher (Fig. 7), but the gain advantage was small and did not increase over time (Fig. 6). By contrast, when Werner et al. (2023) compared truncation selection on GEBV vs. GPMP, the latter achieved more than 3 times as much genetic gain after 20 cycles. Inbreeding was not controlled in that study, and the realized inbreeding rate under GEBV selection was ∼5% compared to ∼1% for GPMP, but this does not appear to explain the dichotomy. Differences in the structure and parameters of the simulated breeding program may be responsible, which indicates more testing is needed.

Even without a gain advantage for COMA, the sparsity of the mating design and flexibility to incorporate mating constraints offer practical incentives over OCS. Earlier versions of COMA have been used for several years in the University of Wisconsin potato breeding program, but the optimal plan was not realized due to frequent and unpredicted problems with reproductive fertility. COMA is used a second time after the crossing is done, converting the realized seed quantities into upper limits for mate allocation based on our greenhouse seedling capacity; the optimal solution guides how many seeds to sow for the next generation.

## Acknowledgments

This research was supported by USDA NIFA Award 2020-51181-32156.

## File S1: Supplemental Methods

Jeff Endelman 2024-10-03

### Eccentricity of MPH level curves

Because the lengths of the ellipse semi-axes are the inverse square-roots of the eigenvalues *λ* of the matrix of the quadratic form (Eq. 4), the eccentricity 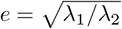. The eigenvalues are the roots of the characteristic equation:

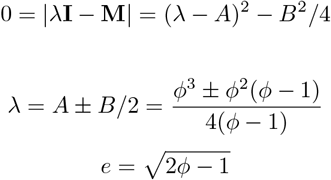

### Equation 9

The OMA objective when SCA is neglected is

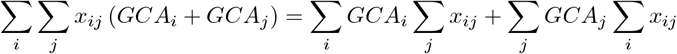

Summing over one index of the mate allocation variable generates sex-specific contributions (Eq. 8), so the above expression simplifies to

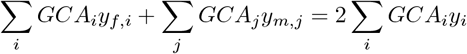

The final expression follows because the index variables are arbitrary, and the total contribution *y*_*i*_ is the average of the sex-specific contributions (Eq. 8). Eq. 9 follows because breeding value is two times GCA at HWE.

### Equation 19

Eq. 18 is the inbreeding coefficient *F*_*ij,ab*_ of an individual in generation *t* + 2 that is the offspring of individuals *ij* and *ab* in generation *t*, with grandparents *i, j, a, b* in generation *t*. The frequency of *ij* is determined by the mate allocation in generation *t*: *x*_*ij*_. Under random mating in generation *t* + 1, the frequency of *ij, ab* offspring is the product of the parental frequencies, *x*_*ij*_*x*_*ab*_, and therefore the average inbreeding coefficient is

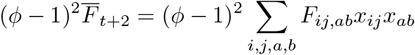

Each summation index goes from 1 to *n*, the number of individuals in generation *t*. Substituting Eq. 18 into above,

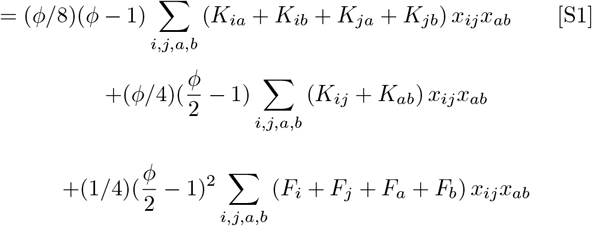

Using the following intermediate results based on Eq. 8,

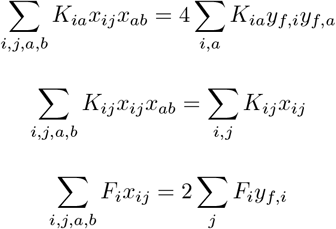

the first line of Eq. S1 becomes

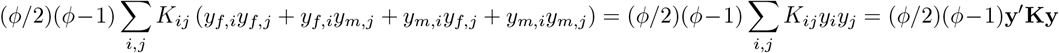

the second line of Eq. S1 becomes

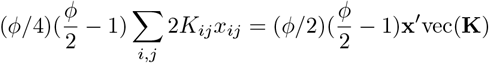

and the third line of Eq. S1 becomes

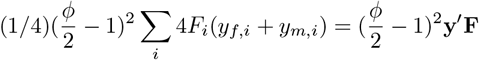

The sum of these three pieces equals Eq. 19.

### Equation 22

The diagonal elements of **G** (VanRaden Method 1) depend on the individual allele dosages *z*_*ik*_, which have population mean *ϕp*_*k*_.

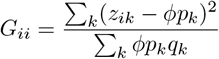

The population mean of *G*_*ii*_ and inbreeding coefficient 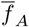 are therefore

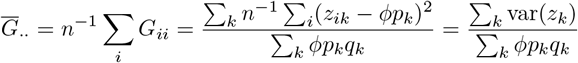

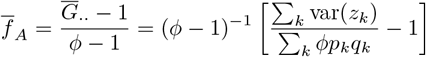

The average inbreeding coefficient from the directional dominance model, 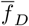, depends on the average value of the dominance regressor (Eq. 1):

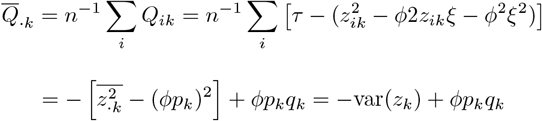

Substituting the above result into following expression (Eq. 21) demonstrates equality with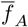.

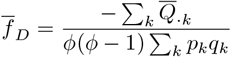

## File S2: Supplemental Tables and Figures

Endelman JB. Genomic prediction of heterosis, inbreeding control, and mate allocation in outbred diploid and tetraploid populations.

**Table S1.**
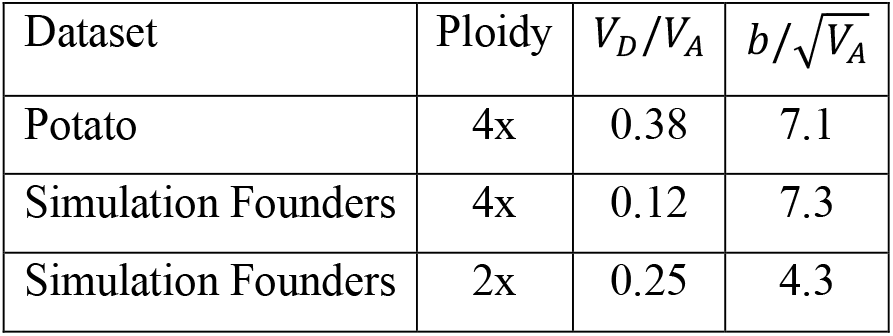
Dominance variance (*VD*) and inbreeding depression (*b*) relative to additive variance (*VA*) and standard deviation, respectively.

| Dataset | Ploidy | $V_D/V_A$ | $b/\sqrt{V_A}$ |
| --- | --- | --- | --- |
| Potato | 4x | 0.38 | 7.1 |
| Simulation Founders | 4x | 0.12 | 7.3 |
| Simulation Founders | 2x | 0.25 | 4.3 |

**Figure S1.**
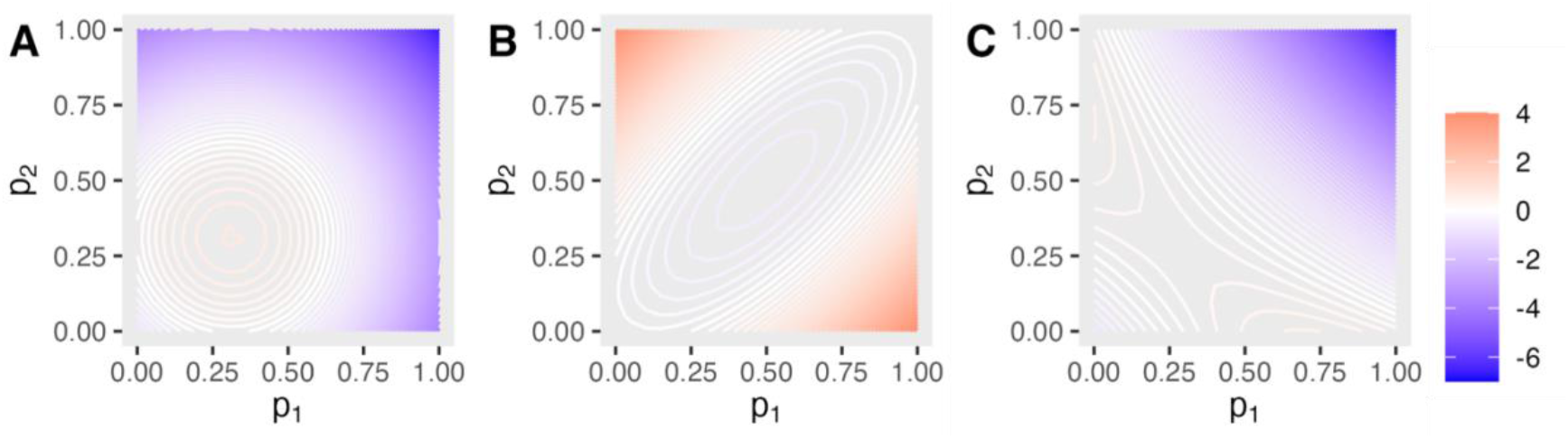
Level curves for mid-parent dominance value (A), mid-parent heterosis (B), and their sum (C), in tetraploids. The population frequency is ¼.

**Figure S2.**
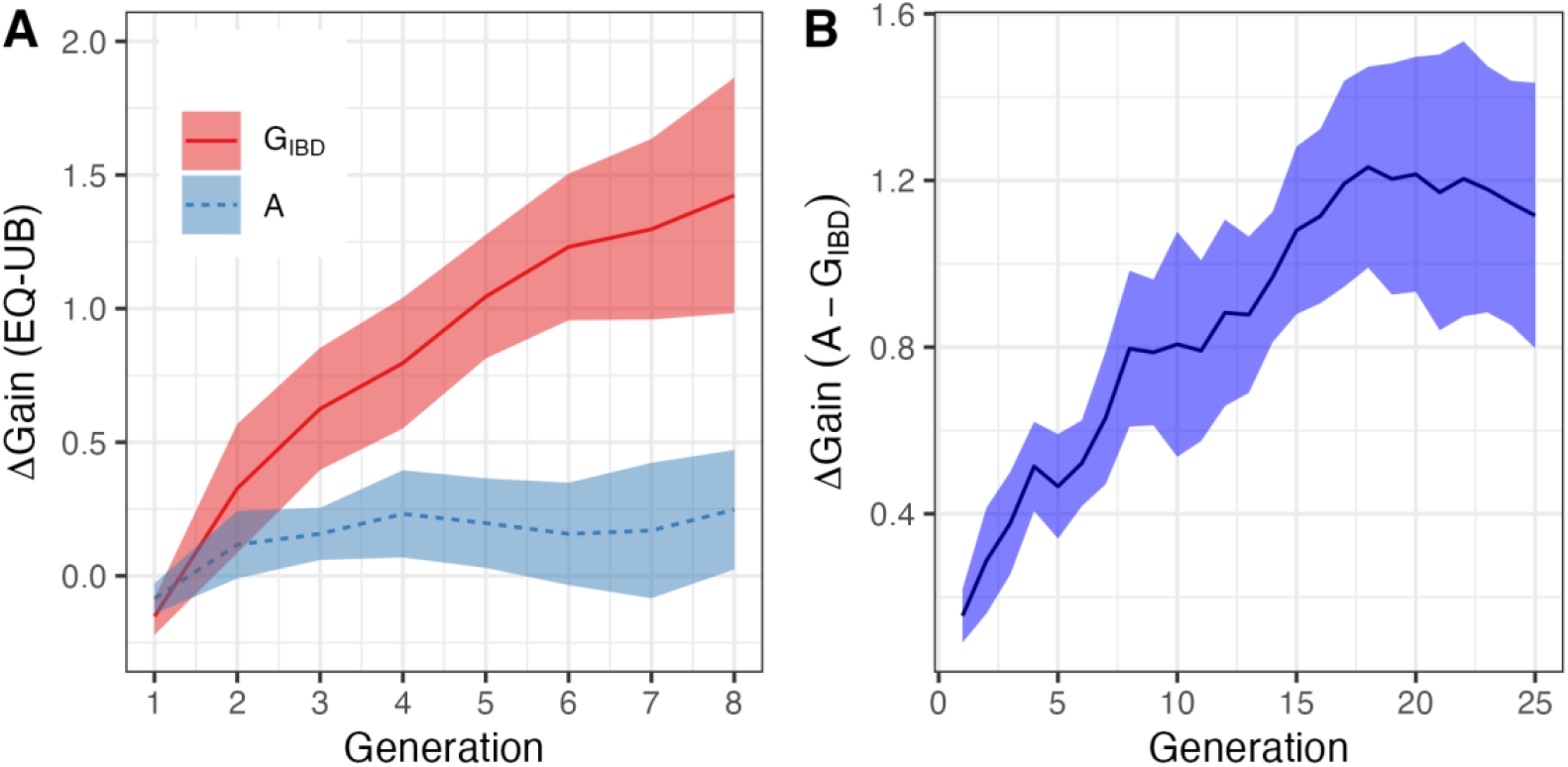
(A) Genetic gain difference at 0.5% inbreeding for equality (EQ) vs. upper bound (UB) constraints with COMA in diploids, using either pedigree or genomic kinship. The solid line is the mean difference across the 10 replicate simulations, and the ribbon is the 95% CI from a paired *t*-test at each generation. (B) Genetic gain difference between pedigree and genomic kinship at 0.5% inbreeding with EQ constraints in diploids.

**Figure S3.**
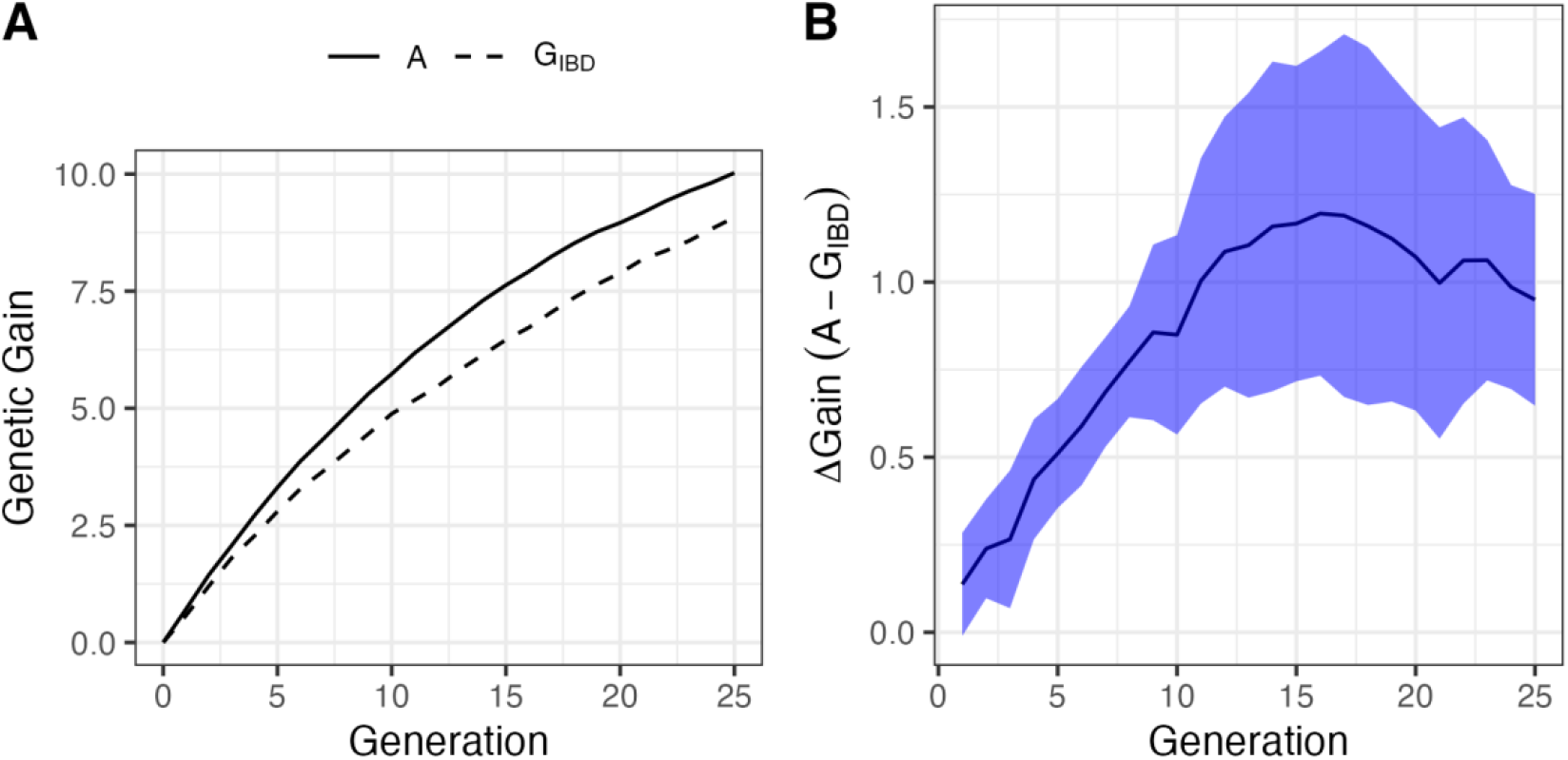
(A) Genetic gain for the 4x simulation at 0.5% inbreeding, using COMA with EQ constraints. (B) Mean difference between the pedigree and genomic kinship results, with the ribbon showing the 95% CI.

**Figure S4.**
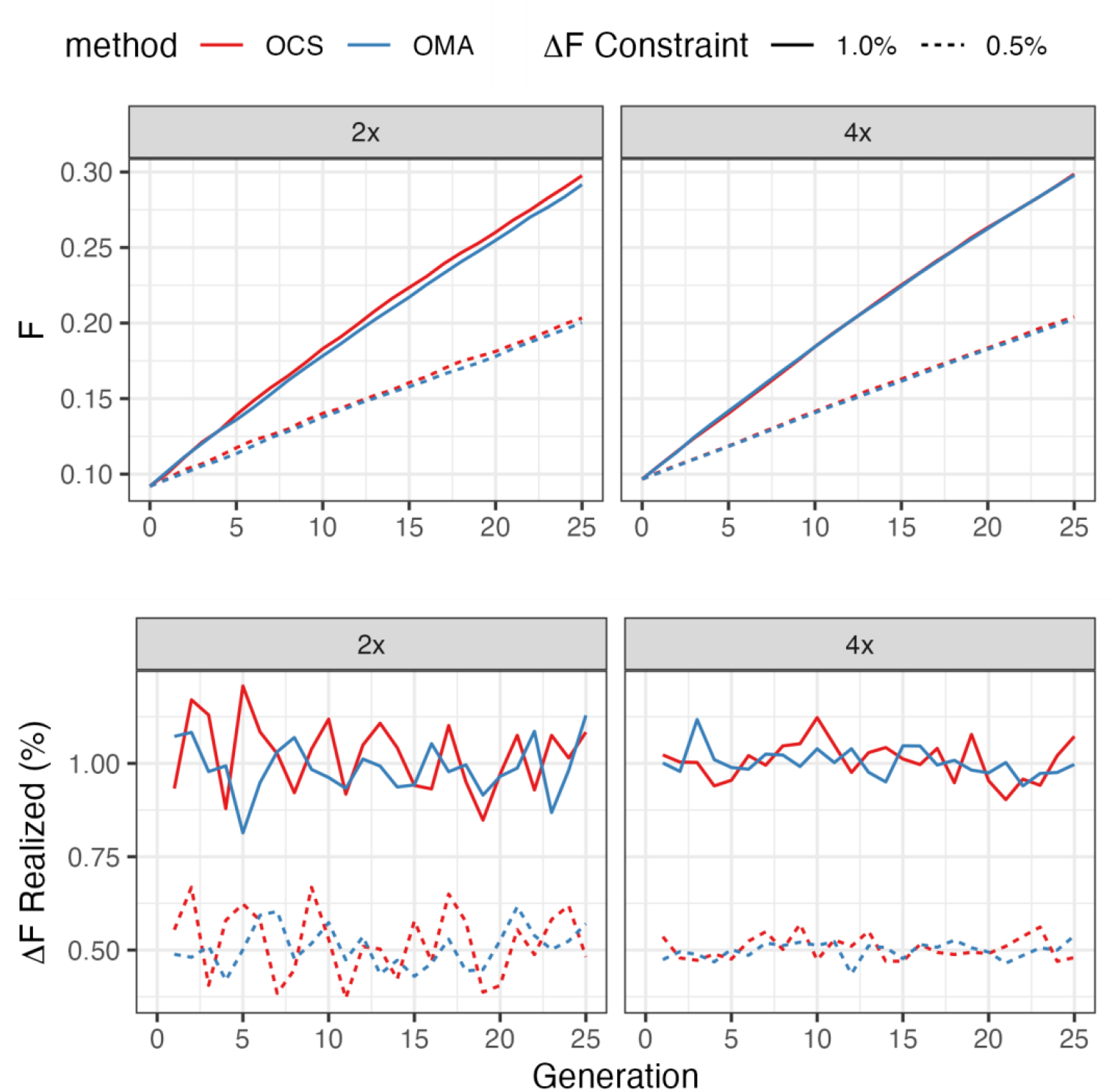
Realized inbreeding coefficients (top) and rates (bottom) at target inbreeding rates of 0.5% and 1.0%. On the left are diploid (2x) results; on the right are tetraploid (4x) results.

**Figure S5.**
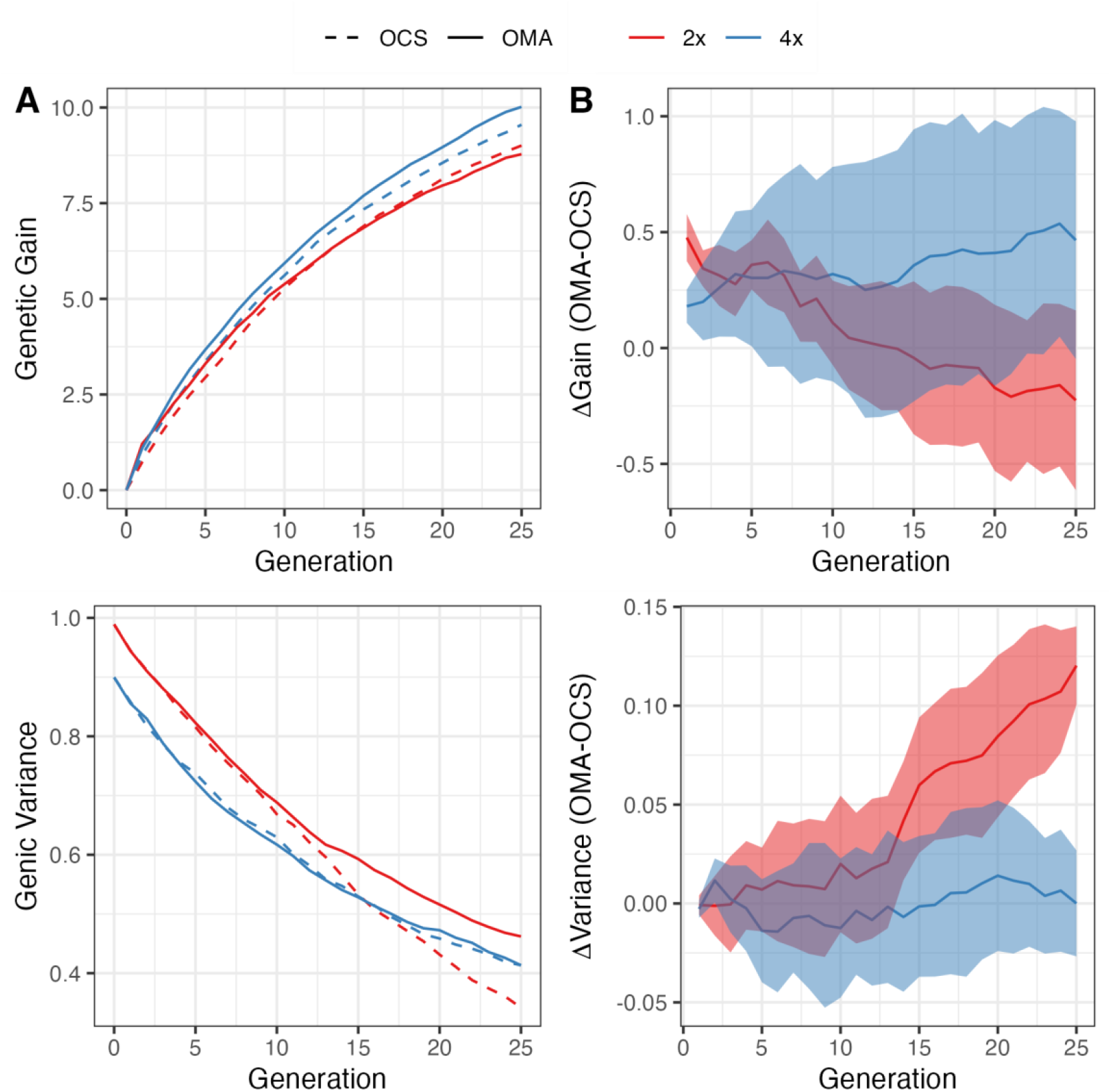
(A) Simulation results for genetic gain and genic variance using OCS vs. OMA at 1.0% inbreeding with pedigree kinship. (B) The solid line is the mean difference across the 10 replicate simulations, and the ribbon is the 95% CI from a paired *t*-test each generation.

**Figure S6.**
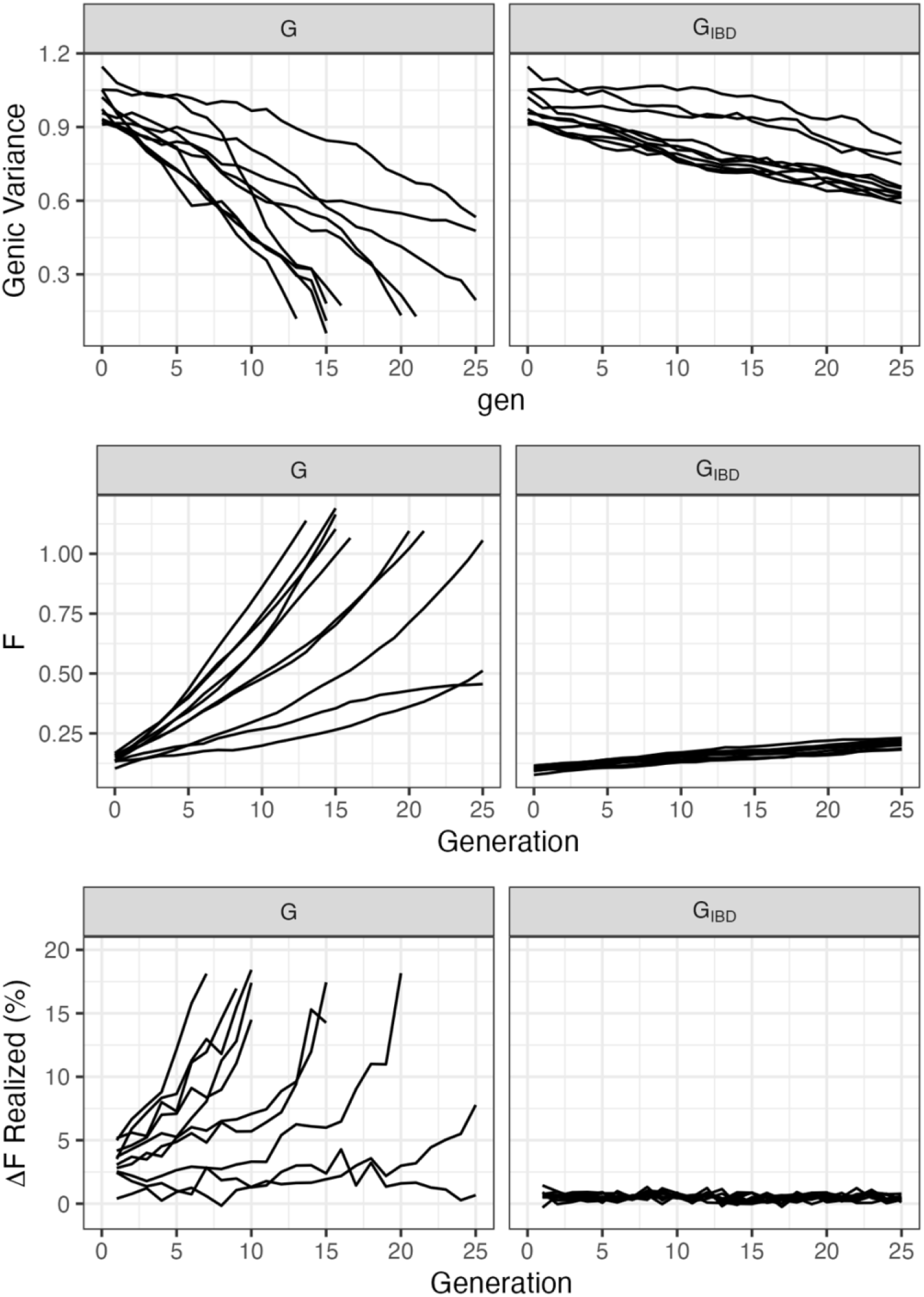
Comparison of **G** (VanRaden Method 1) vs. **G**IBD for genomic control of inbreeding with COMA at target rate 0.5%. Each line is an independent simulation. Inbreeding coefficients for **G** are defined in Eq. 22.

## File S3: COMA Vignette

Jeff Endelman

The COMA package implements a new algorithm for optimum mate allocation (OMA), as well as the more traditional method, optimum contribution selection (OCS).

COMA requires two input files. The first file contains the marker effects and allele dosage data for the parental candidates. Dominance marker effects are optional, but when included should correspond to a breeding value parameterization, including the effect of heterosis/inbreeding depression (Endelman 2023). The `blup` command in R/StageWise is one option to compute marker effects.

The vignette dataset comes from the University of Wisconsin potato breeding program. There are 170 tetraploid clones genotyped at 12K markers, and the marker effects are derived from a multi-trait index.

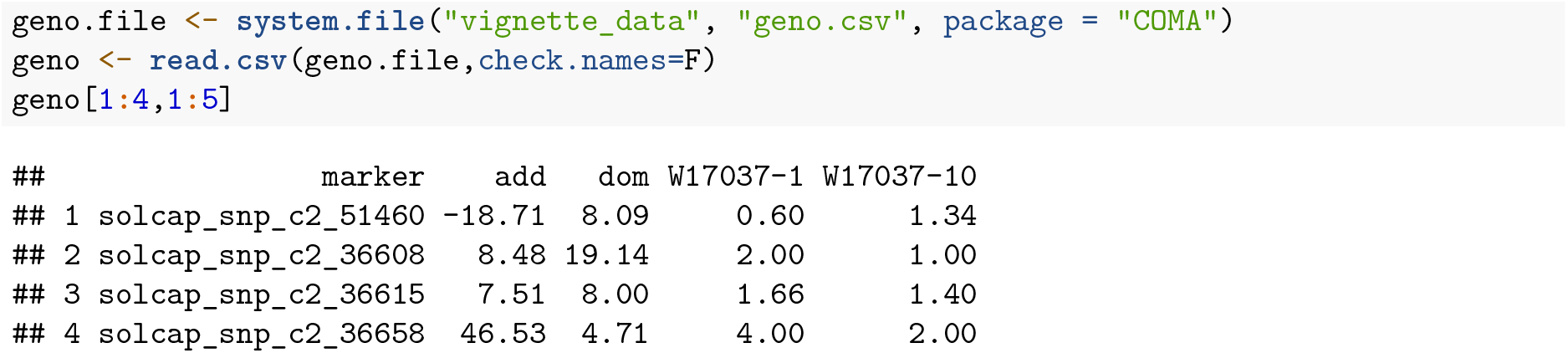

The second input file is the kinship matrix to control inbreeding. Based on simulation results (Endelman 2024), the pedigree IBD kinship matrix is recommended for either OCS or OMA. The kinship matrix for the potato clones was generated using R package AGHmatrix.

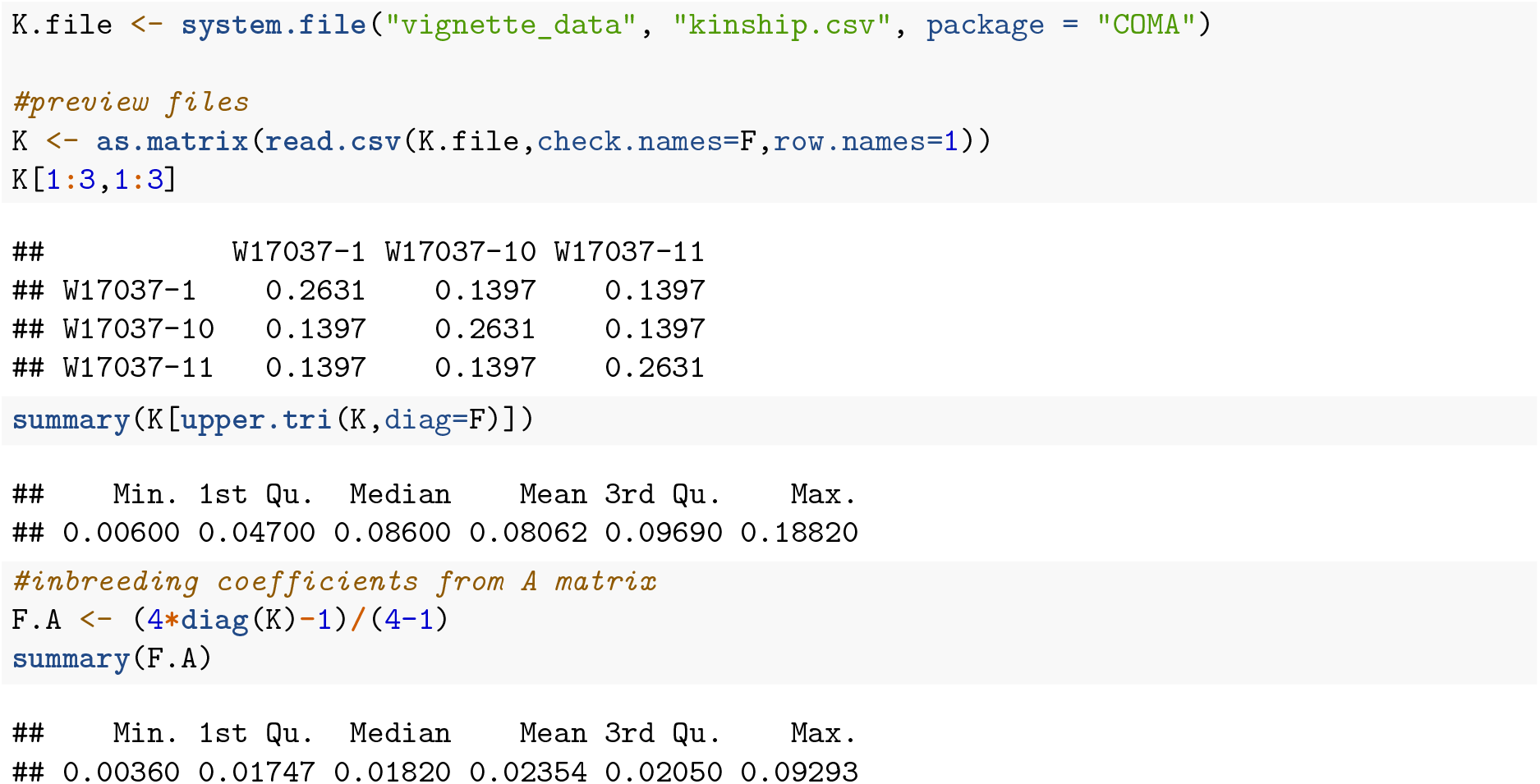

Now we are ready for `read_data`:

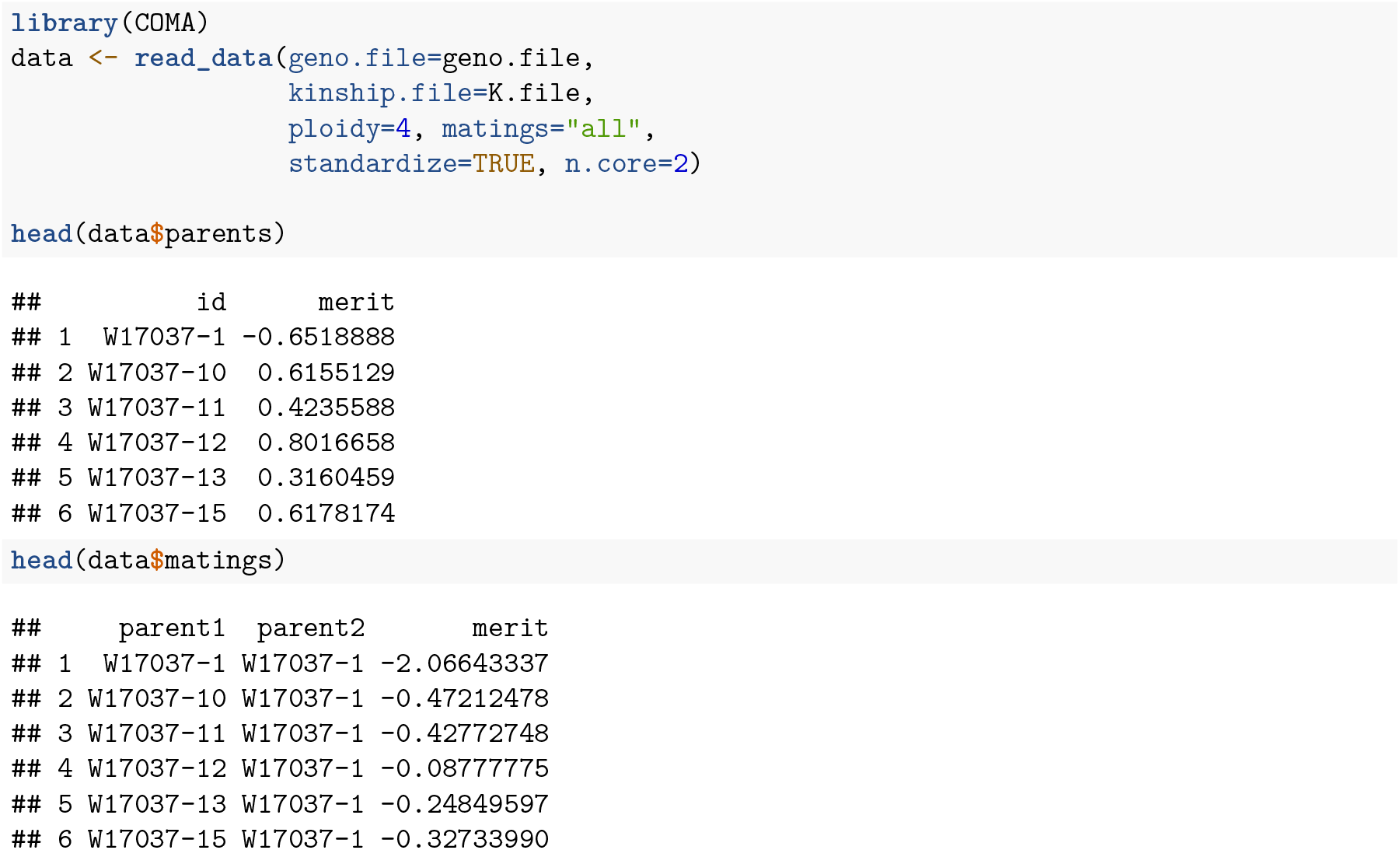

The `matings` argument for `read_data` indicates how to create the table of possible matings. The above example used “all”, which creates all unique parental combinations. Consult the function documentation for other options. When `standardize`=TRUE, genetic values are scaled by the additive standard deviation (*σ*_*A*_). This is for convenience when interpreting the response values.

As shown above, the `parents` output contains the predicted merit (GEBV) for each individual. The `matings` output contains the predicted merit (F1 progeny mean) for each possible mating. For dioecious species (with separate sexes), a data frame with sex information is needed as argument `sex in read_data`, and the column headers for the matings are “female”,”male” instead of “parent1”,”parent2”.

### OMA and OCS

The first argument of the `oma` function is `dF`=c(min,max), which represents lower and upper bounds on the inbreeding rate for the next generation, dF1. The upper bound is also applied to the average inbreeding rate over two generations, dF2, assuming random mating of the progeny. The second argument, `parents`=data.frame(id,min,max), contains lower and upper bounds on the contribution of each parent (do not include a column for parental merit). The third argument, `matings`, is a data frame of the merit for each possible mating, as well as lower and upper bounds on the allocation.

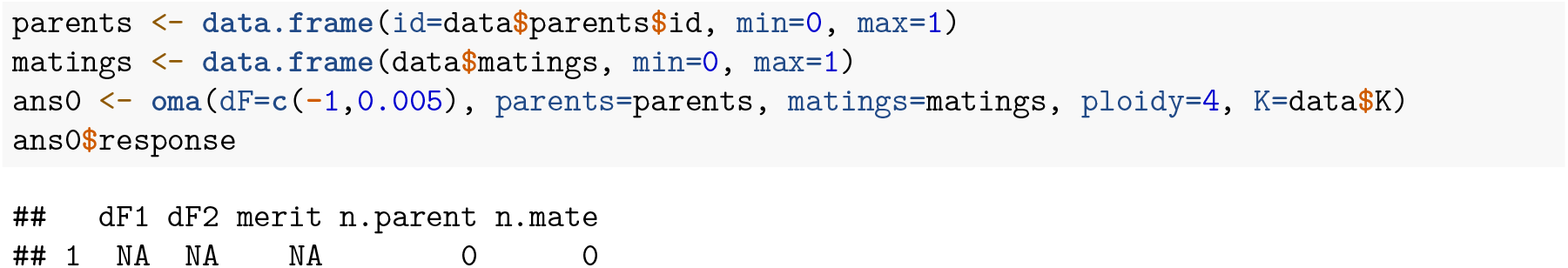

The above result shows there is no feasible solution at 0.5% inbreeding for both generations, due to the relatedness of the clones. The argument `dF.adapt` can be used to automatically increasing the upper bound until a solution is found:

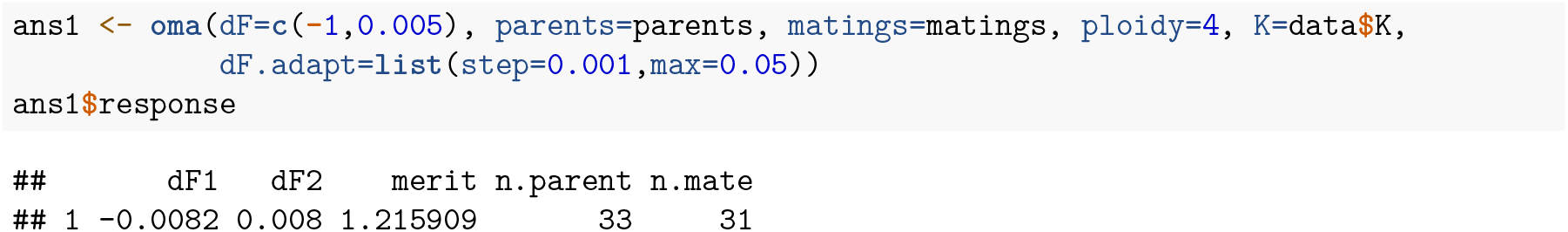

With no lower bound, a solution was found at dF2 = 0.8%, while the negative value for dF1 indicates the progeny would be 0.8% more outbred than the parents. When a similar approach was used in a simulation study (Endelman 2024), the population became trapped in a local optimum with limited long-term gain. Imposing a lower bound of 0.5% for dF1 led to more long-term gain in the simulation. As the following code shows, for the potato dataset, this required increasing the dF2 rate to 1.0%.

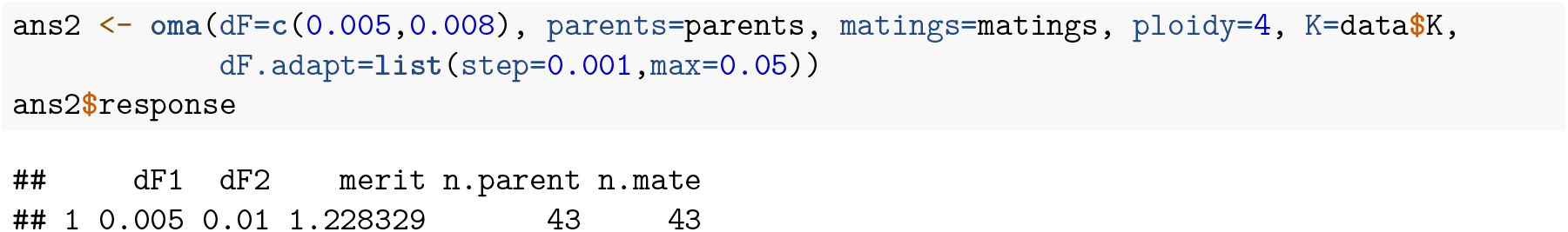

Interestingly, the second solution uses more parents even though the inbreeding rate is higher. The optimal parental contributions are returned as `oc`, and the optimal mate allocations are returned as `om`.

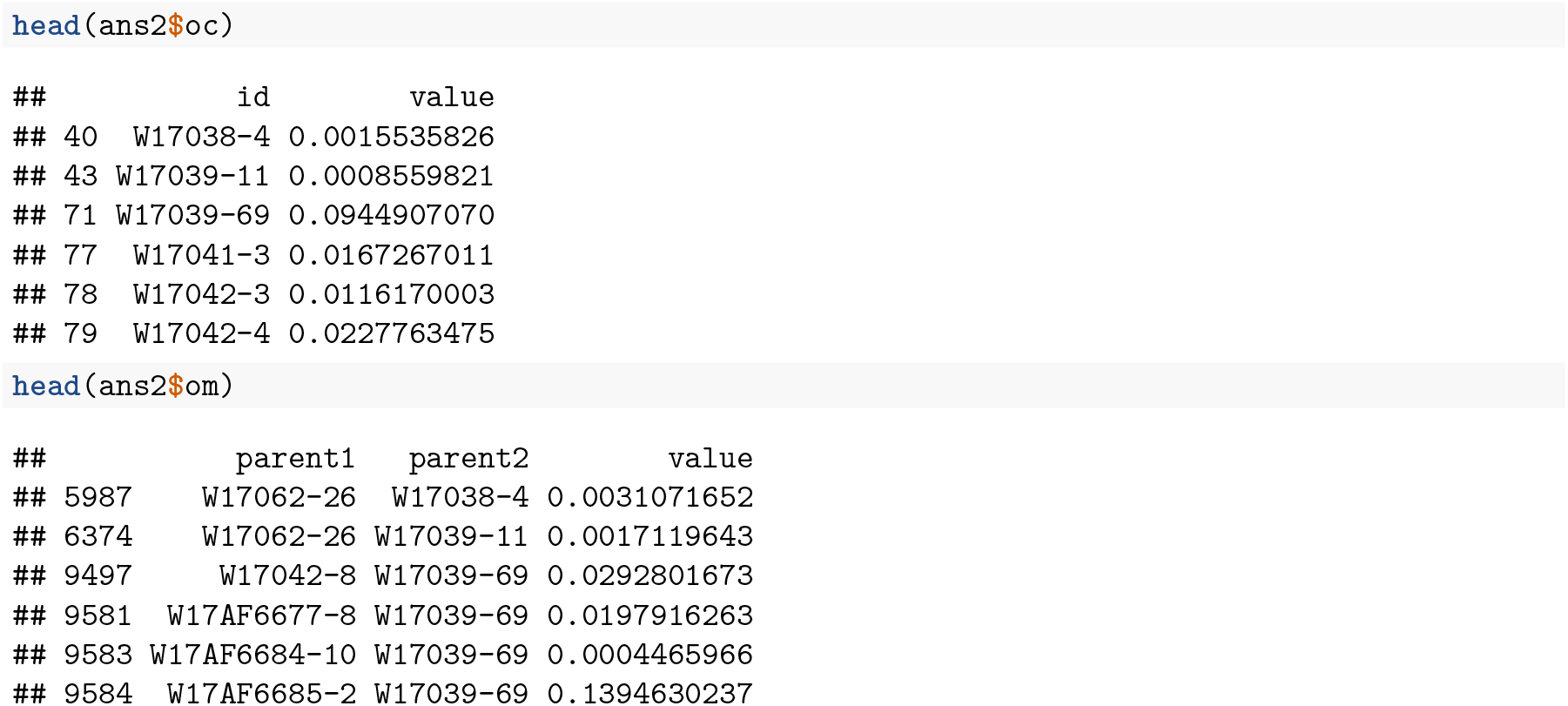

The parental contributions of the two solutions can be visually compared with the COMA function `plot_ribbon`:

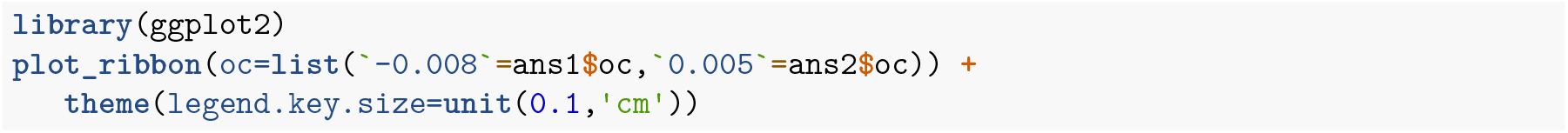
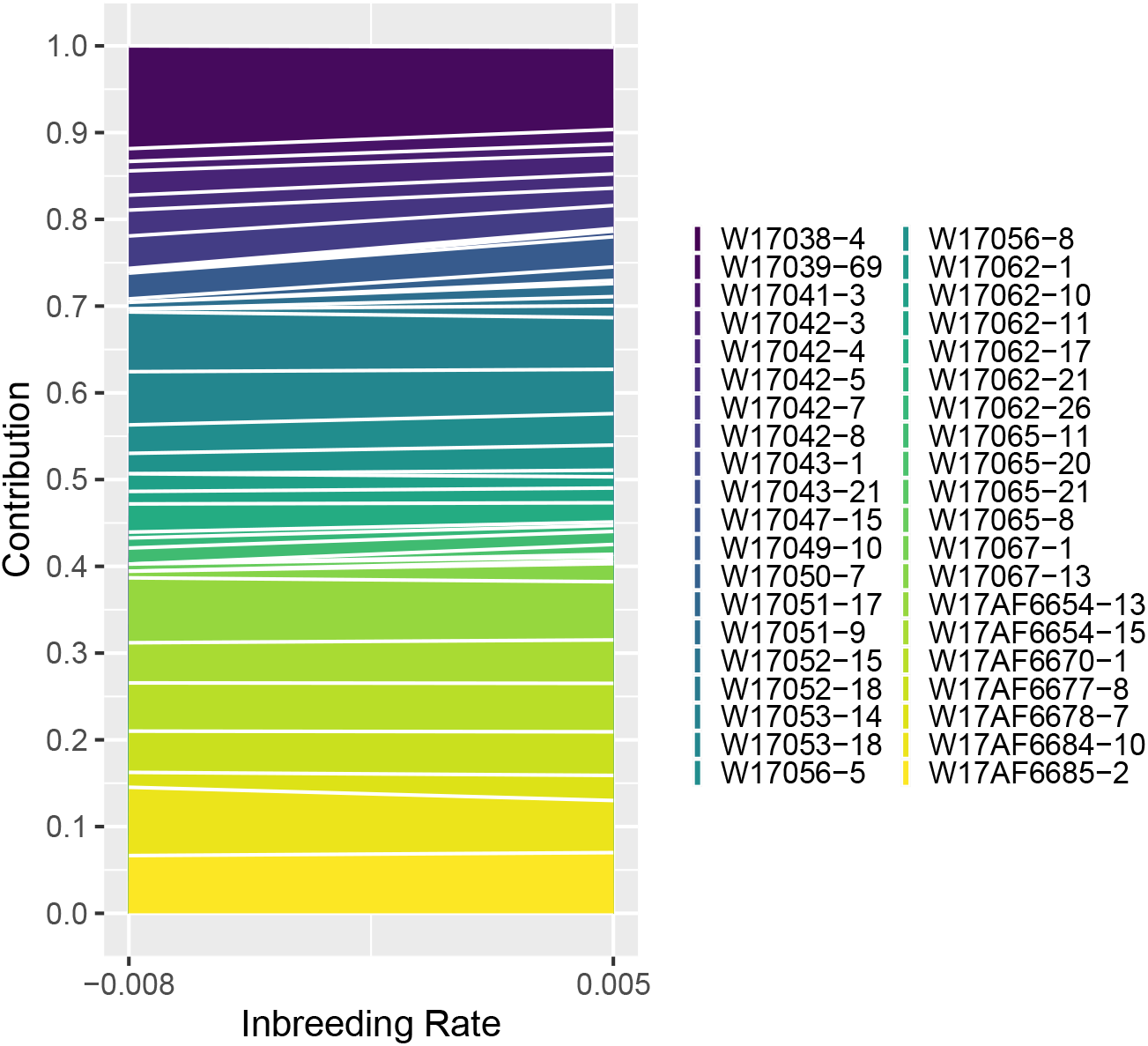

COMA also has a function for OCS, which optimizes parental contributions but not specific matings. The arguments are similar, except dF is only a single number for the upper bound on group kinship, and the `parents` data frame includes a column for merit:

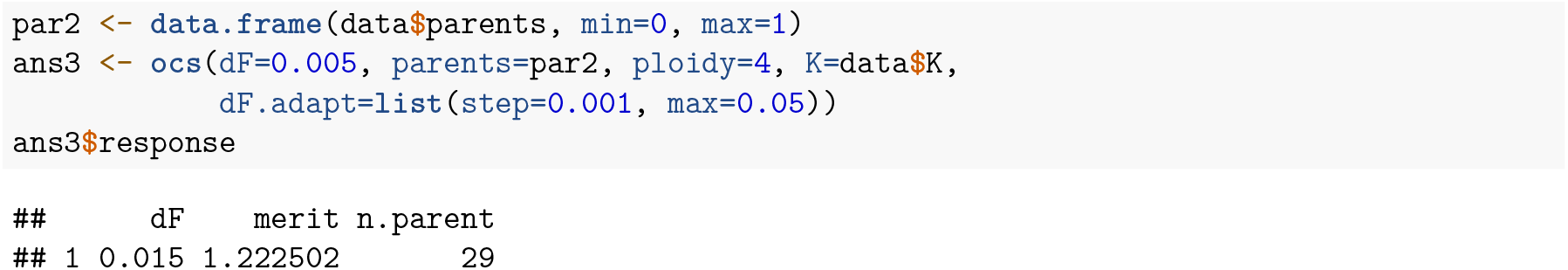

The smallest feasible inbreeding rate under OCS was 1.5%. The predicted merit is based on parental breeding values and therefore neglects specific combining ability (SCA). When SCA was accounted for, the OCS solution (using random mating) decreased from 1.2 to 1.0 *σ*_*A*_.

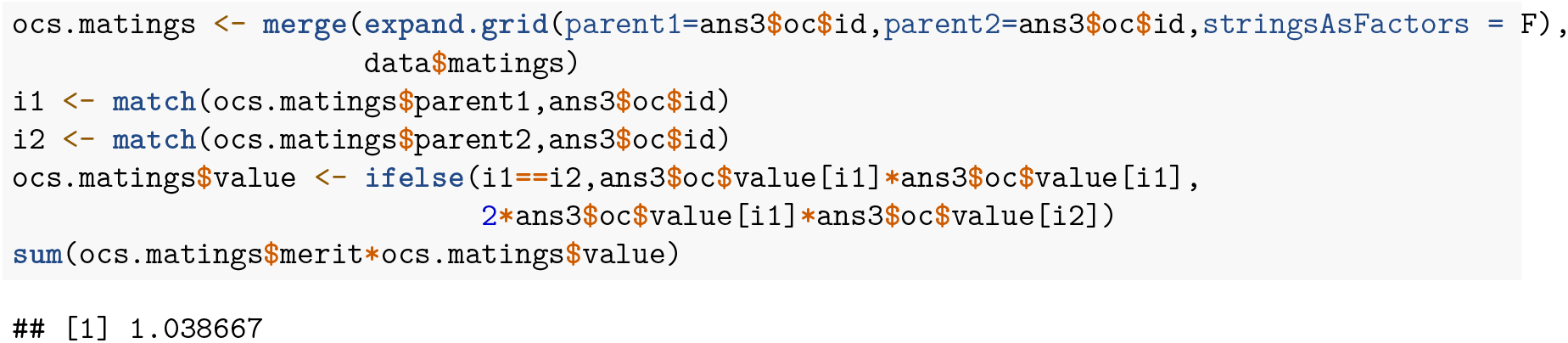

### Other Constraints

Because COMA is based on convex optimization, additional linear constraints can be incorporated. The potato dataset contains presence/absence information for a genetic marker for potato virus Y resistance (PVYR), which is a critical trait. Two different selection approaches will be illustrated: (1) only allowing matings with a resistant parent; (2) imposing a lower bound on the R gene frequency.

The following code implements method 1 with the 0.5% lower bound:

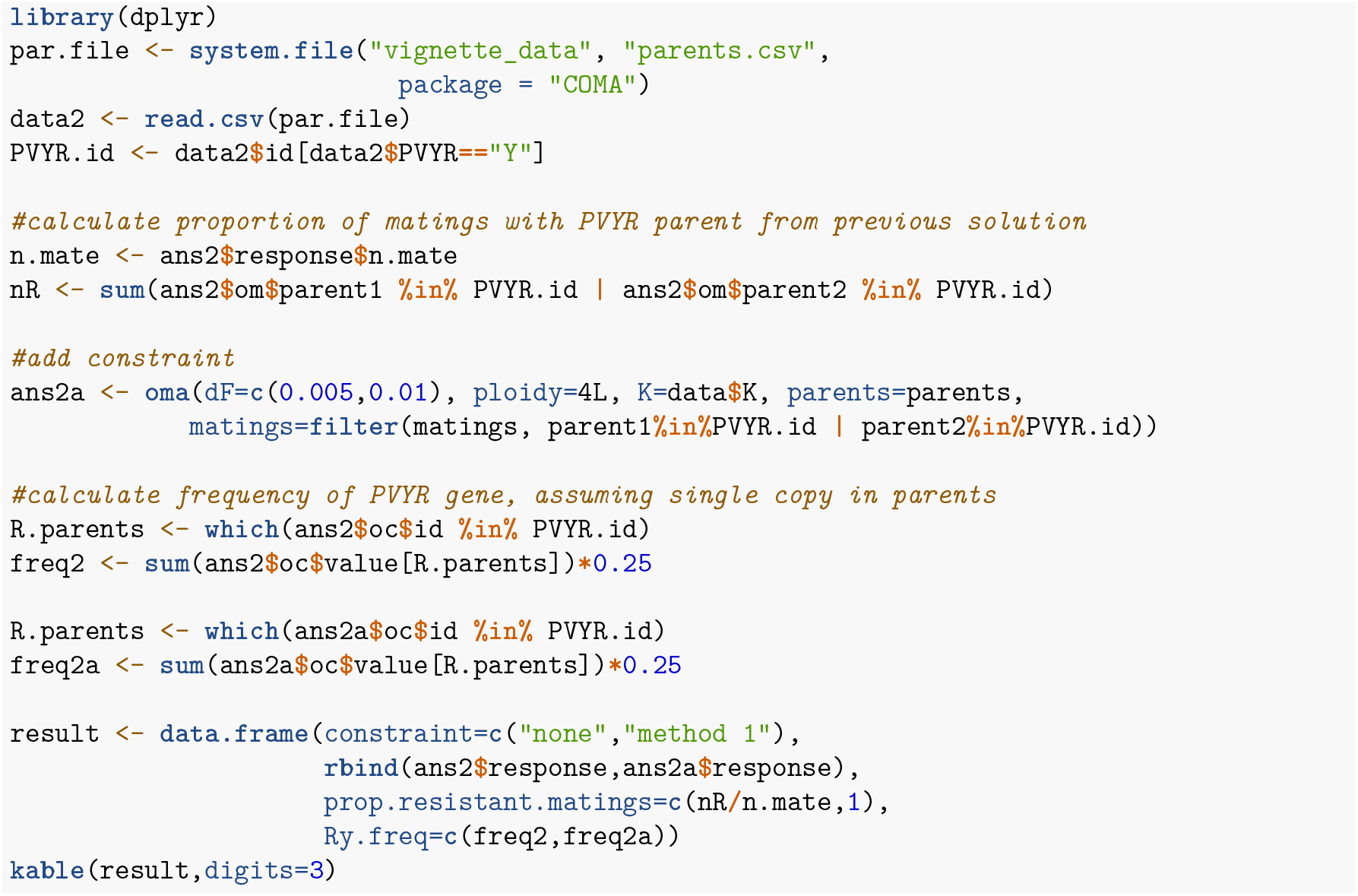

| constraint | dF1 | dF2 | merit | n.parent | n.mate | prop.resistant.matings | Ry.freq |
| --- | --- | --- | --- | --- | --- | --- | --- |
| none | 0.005 | 0.01 | 1.228 | 43 | 43 | 0.442 | 0.086 |
| method 1 | 0.005 | 0.01 | 1.018 | 42 | 40 | 1.000 | 0.125 |

The above table compares the solution with and without the PVYR constraint. As expected, increasing the proportion of PVYR matings to 100% reduced the predicted merit. The frequency of the R gene in the progeny is also shown, assuming a single copy in the resistant parents.

Under method 2, the R gene frequency is constrained directly, which is less restrictive than method 1 because some matings could have two PVYR parents while others have none. Both ocs and oma allow linear inequality or equality constraints on any linear combination of the parental contributions. When the coefficients, **s**, of the contribution variables, **y**, are the R gene frequencies of each parent, this creates the desired constraint. The coefficients are included as an additional column in the `parents` data.frame, and the column header is parsed to complete the equation. In this case, the header “gt0.125” represents greater than or equal to 0.125:

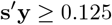

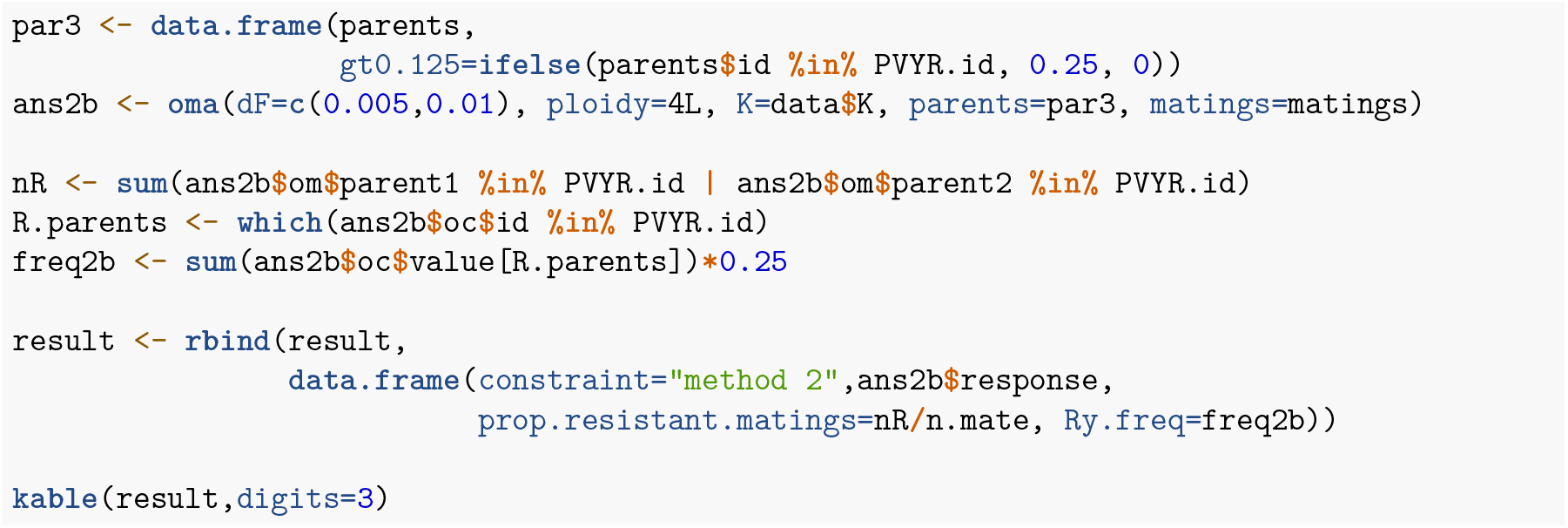

| constraint | dF1 | dF2 | merit | n.parent | n.mate | prop.resistant.matings | Ry.freq |
| --- | --- | --- | --- | --- | --- | --- | --- |
| none | 0.005 | 0.01 | 1.228 | 43 | 43 | 0.442 | 0.086 |
| method 1 | 0.005 | 0.01 | 1.018 | 42 | 40 | 1.000 | 0.125 |
| method 2 | 0.005 | 0.01 | 1.070 | 43 | 42 | 0.628 | 0.125 |

As expected, the predicted merit under method 2 was higher than method 1.

## Package ‘COMA’

October 1, 2024

**Title** Convex Optimization of Mate Allocation

**Version** 0.19

**Author** Jeffrey B. Endelman

**Maintainer** Jeffrey Endelman

**Description** Convex Optimization of Mate Allocation

**Depends** R (>= 4.0)

**License** GPL-3

**RoxygenNote** 7.2.3

**Encoding** UTF-8

**Imports** ggplot2, CVXR, tidyr

**Suggests** knitr, rmarkdown, Rmosek

### Index 6

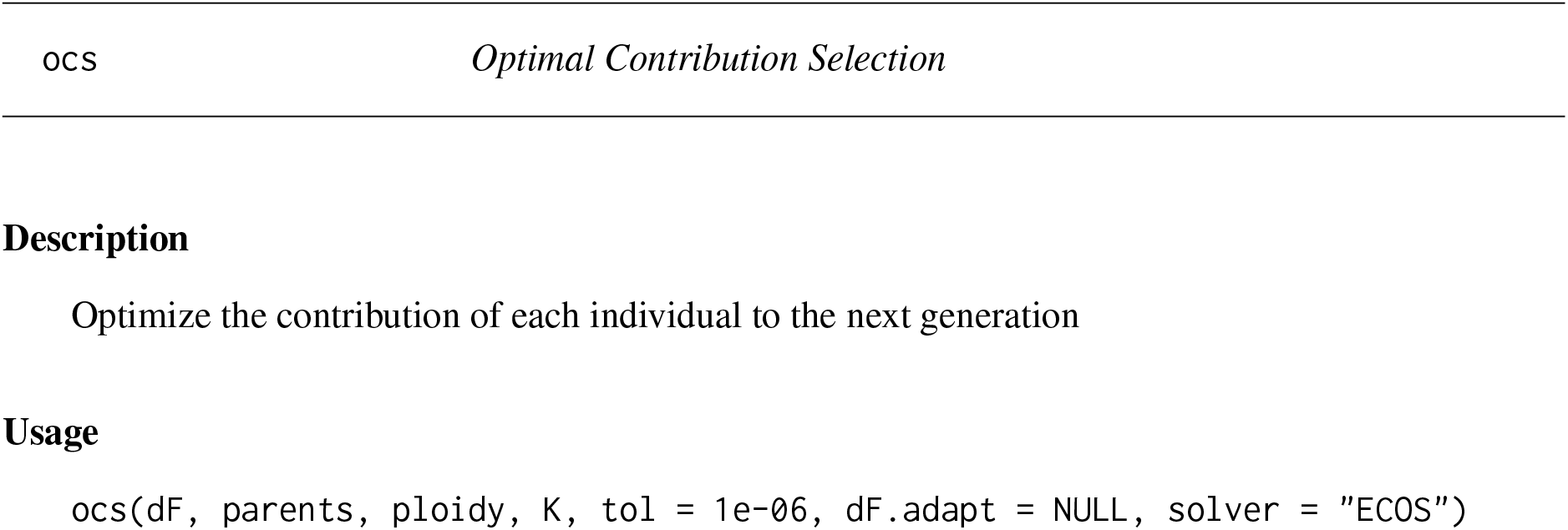
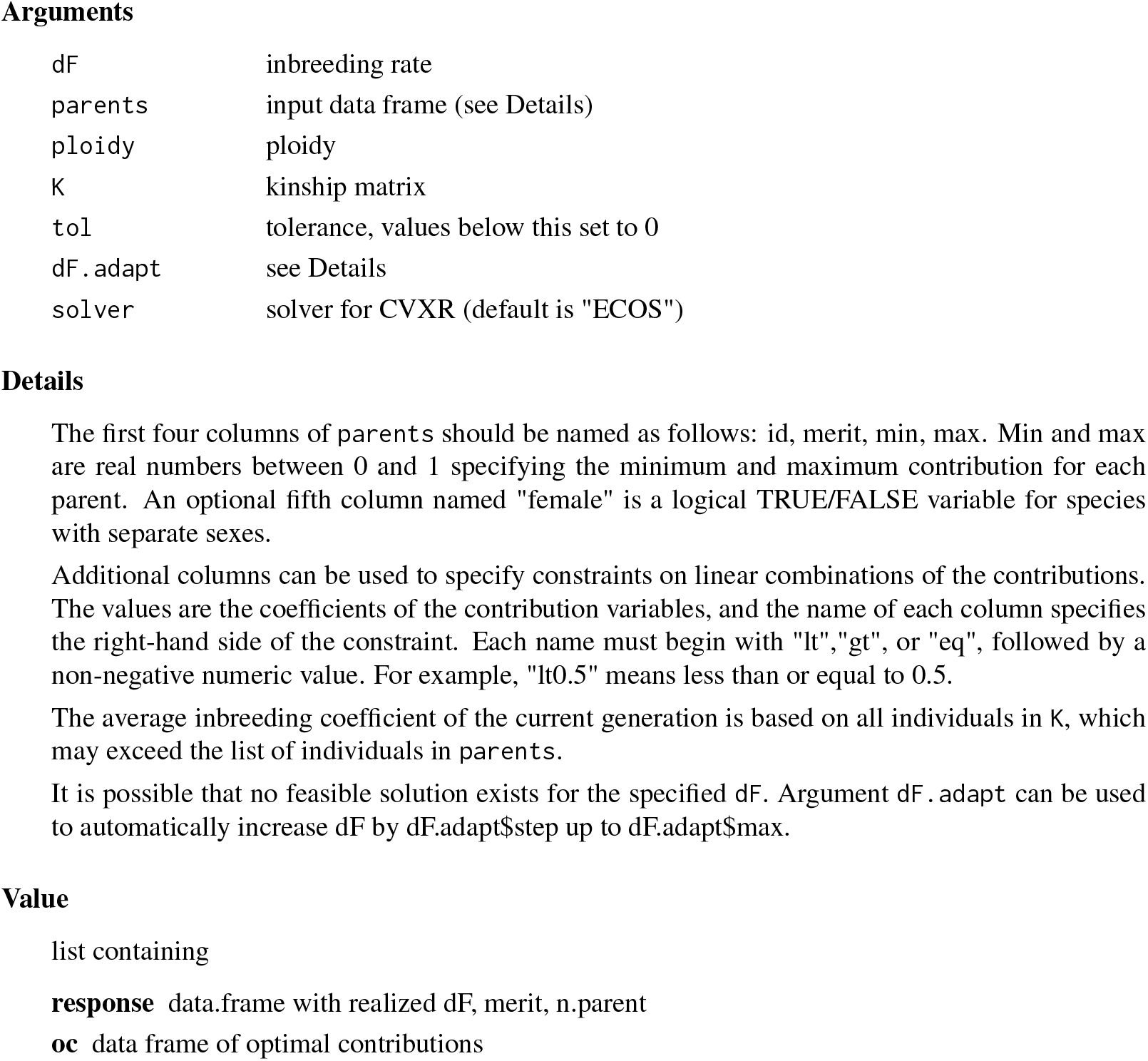

|  |  |
| --- | --- |
| ocs | *Optimal Contribution Selection* |
| Arguments |  |
| dF | inbreeding rate |
| parents | input data frame (see Details) |
| ploidy | ploidy |
| K | kinship matrix |
| tol | tolerance, values below this set to 0 |
| dF.adapt | see Details |
| solver | solver for CVXR (default is "ECOS") |

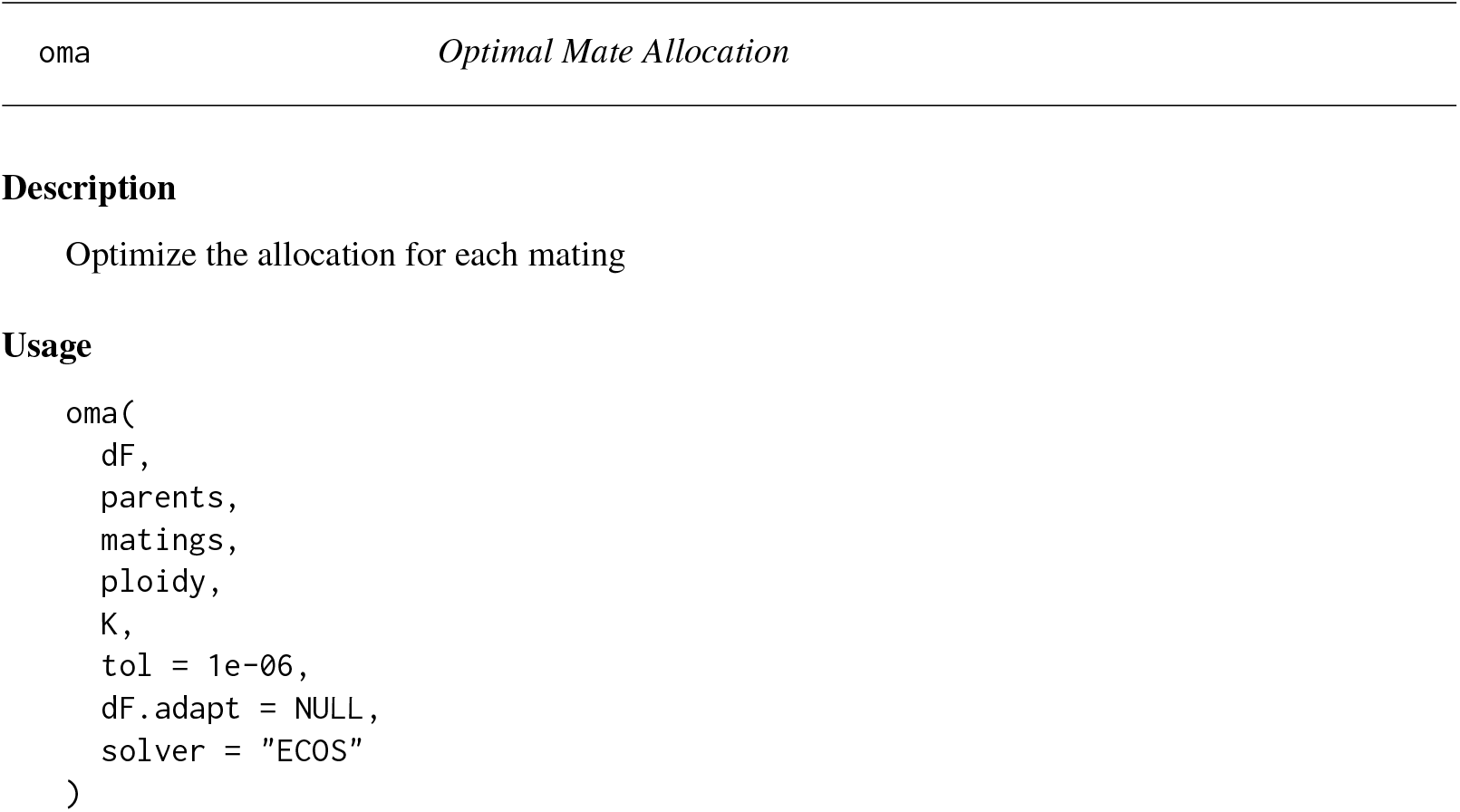
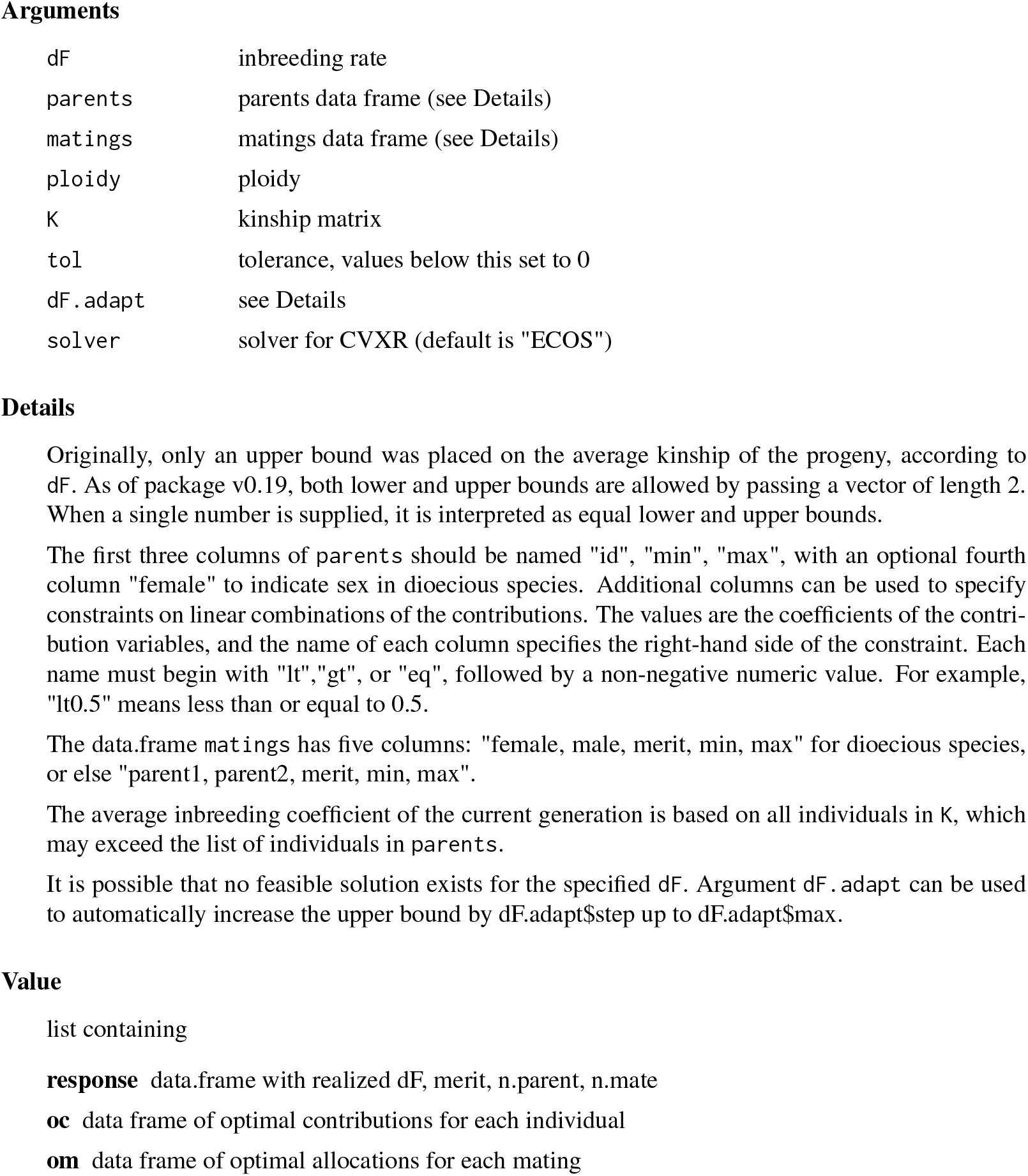

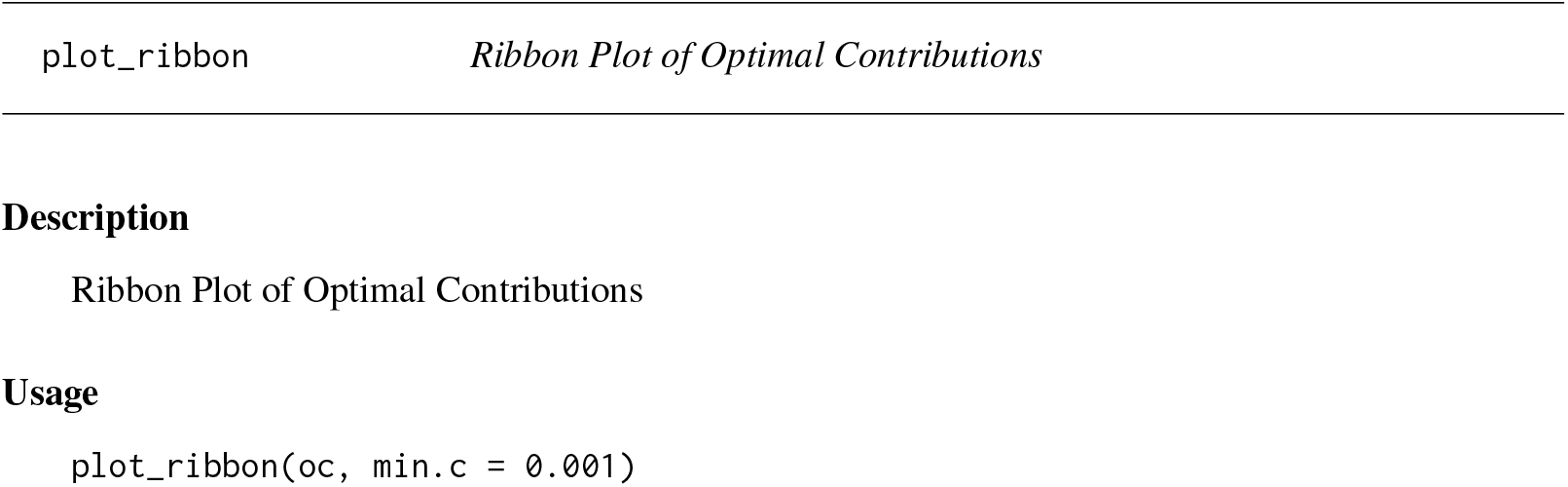
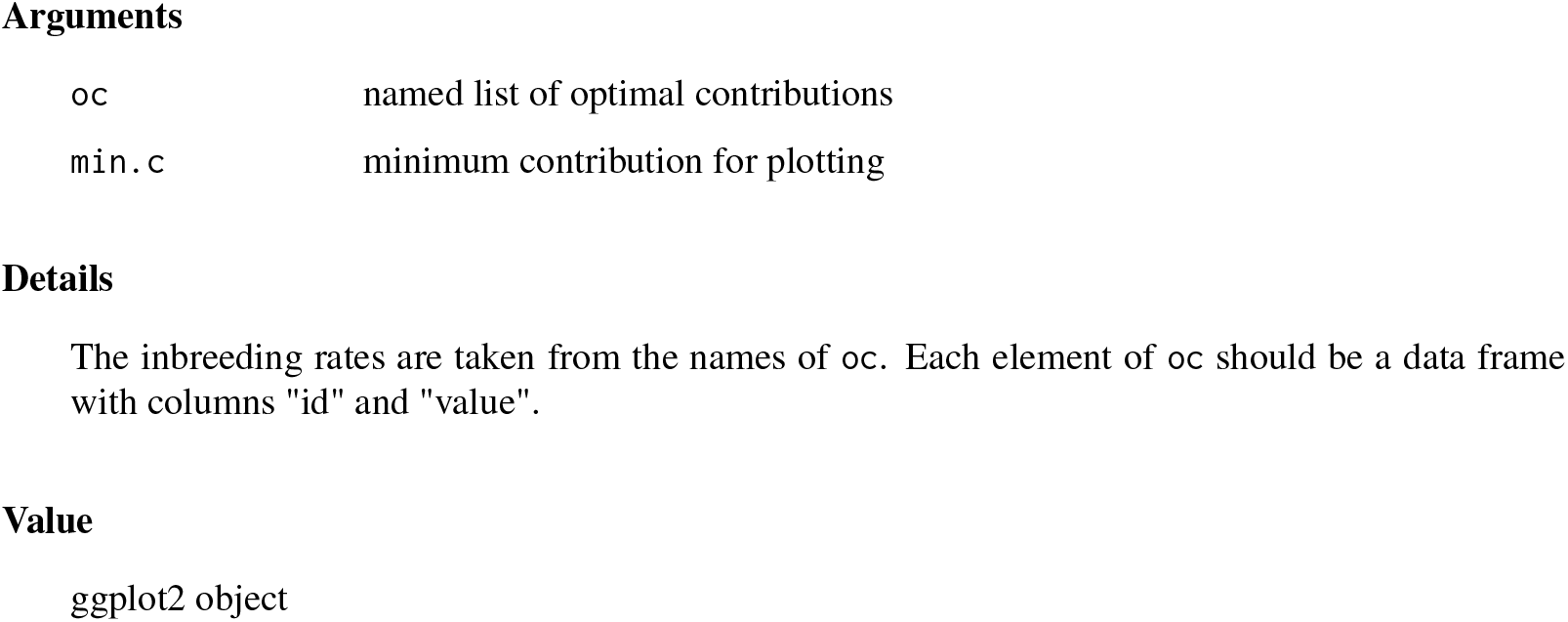

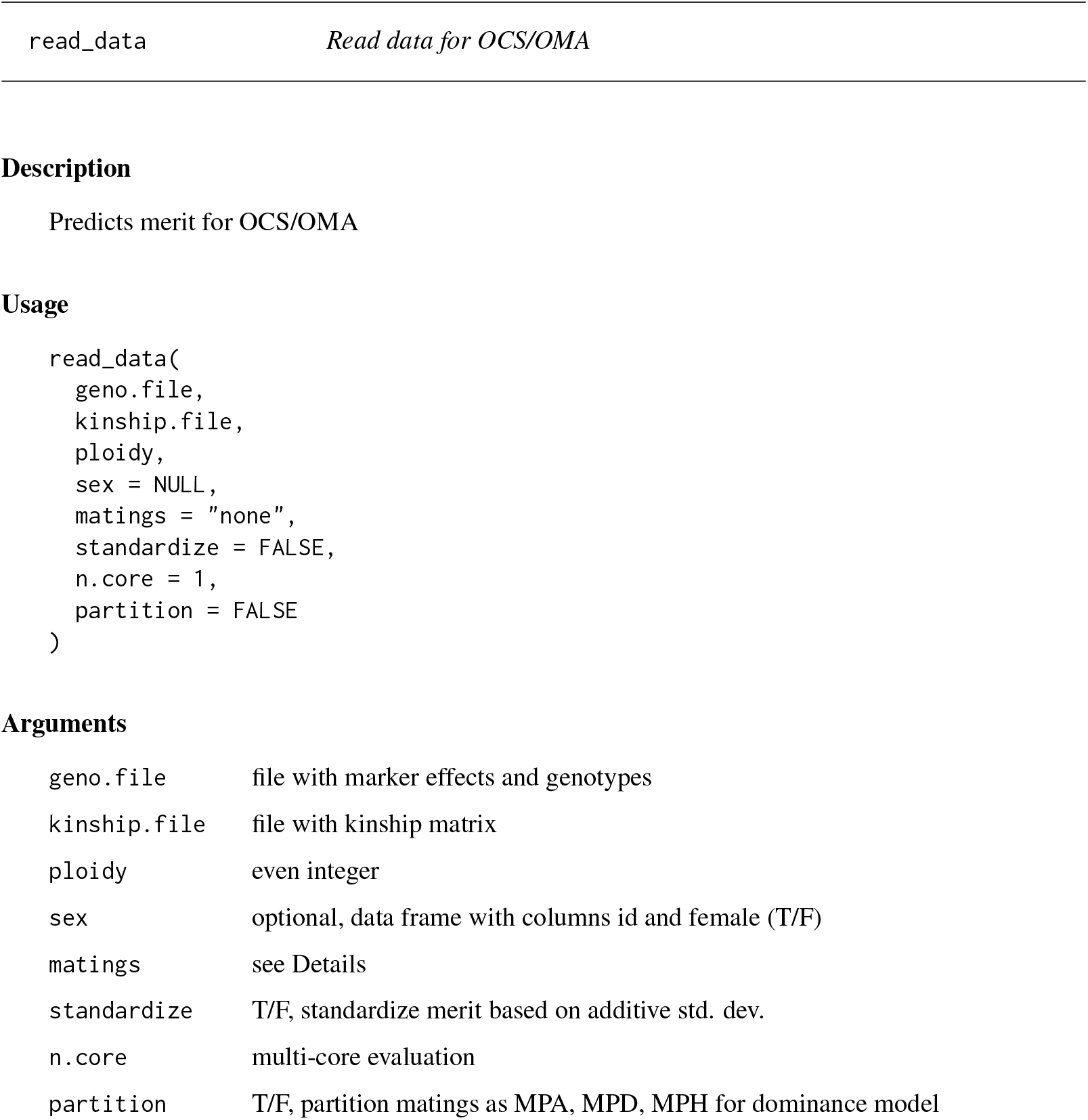
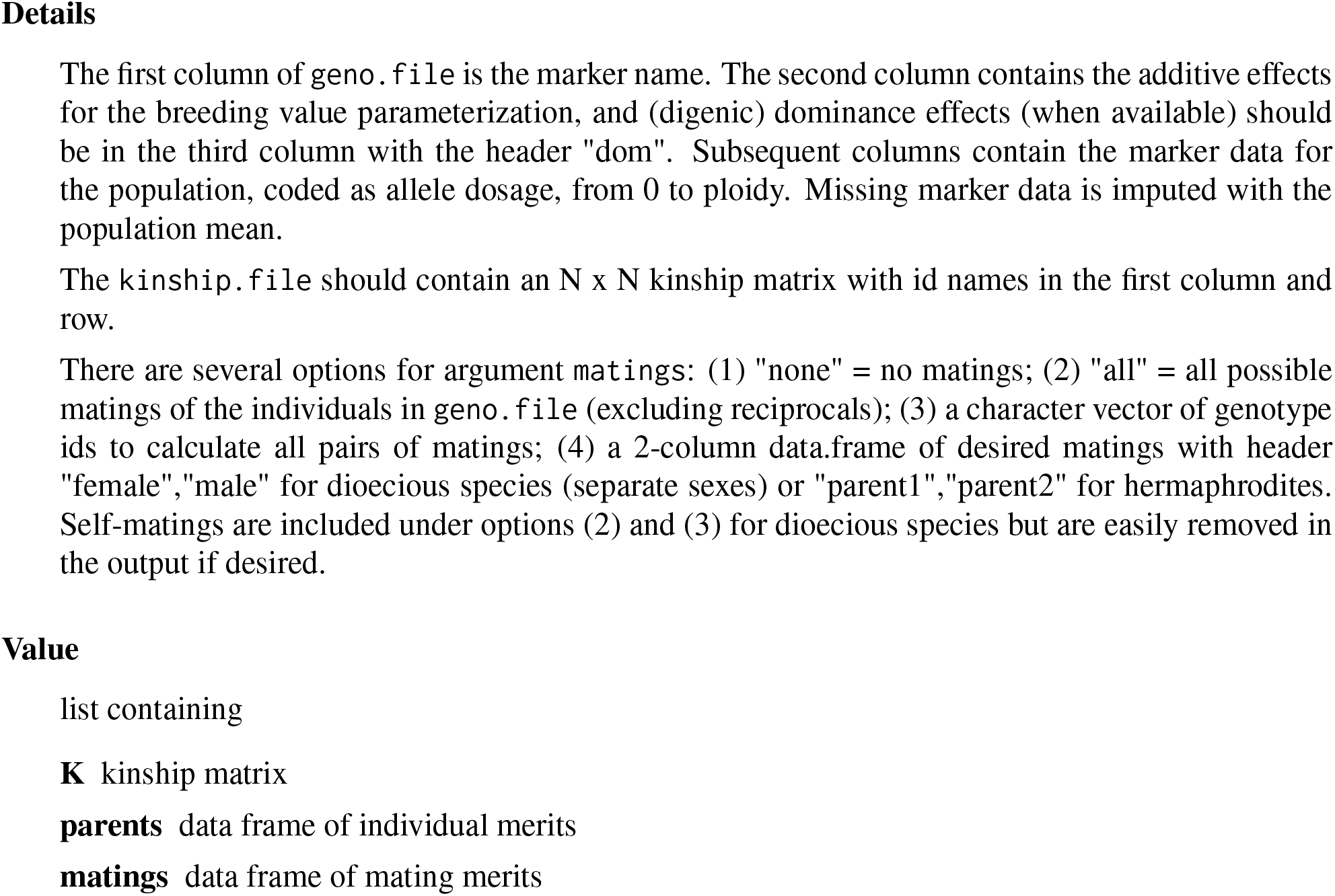

## Index

`ocs`, 1

`oma`, 2

`plot_ribbon`, 3

`read_data`, 4

## File S4: SimPlus Vignette

Jeff Endelman

SimPlus was created for testing COMA with AlphaSimR. COMA provides functions for optimum contribution selection (OCS) and optimum mate allocation (OMA). For OCS, the objective is to maximize the average GEBV of the parents, weighted by their contributions. For OMA, the objective is to maximize the average GPMP (genomic prediction of mate performance), weighted by the mate allocations.

The following code creates a founder population in mutation-drift equilibrium and sets the variance parameters for a trait with additive and dominance effects.

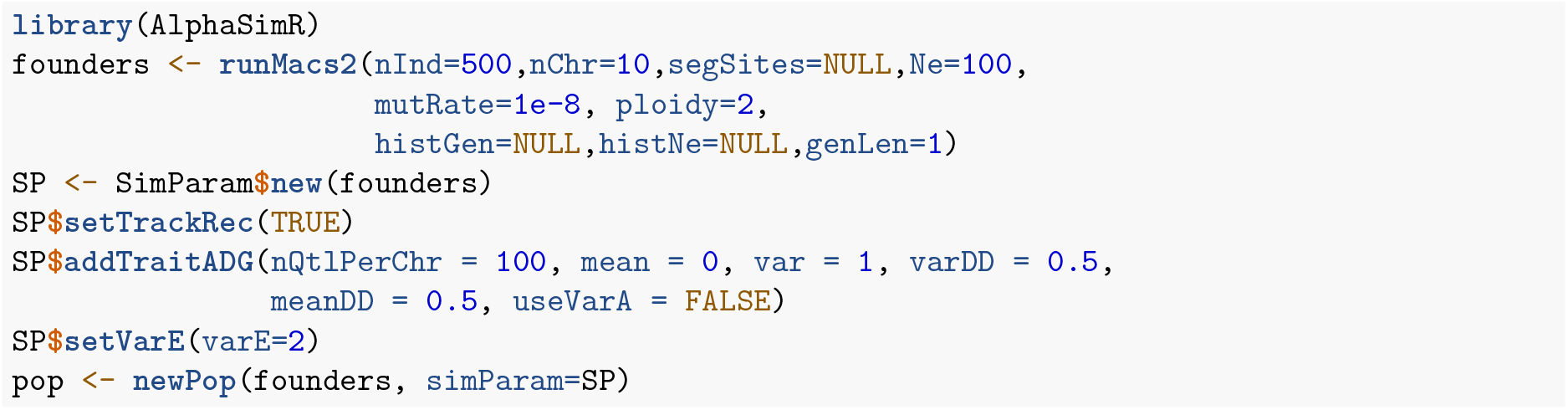

### OCS-A-Pheno

The following code runs 20 generations of mass selection at 1% inbreeding, using phenotypes for genetic merit. The population is saved after 5 generations to compare the response with genomic selection.

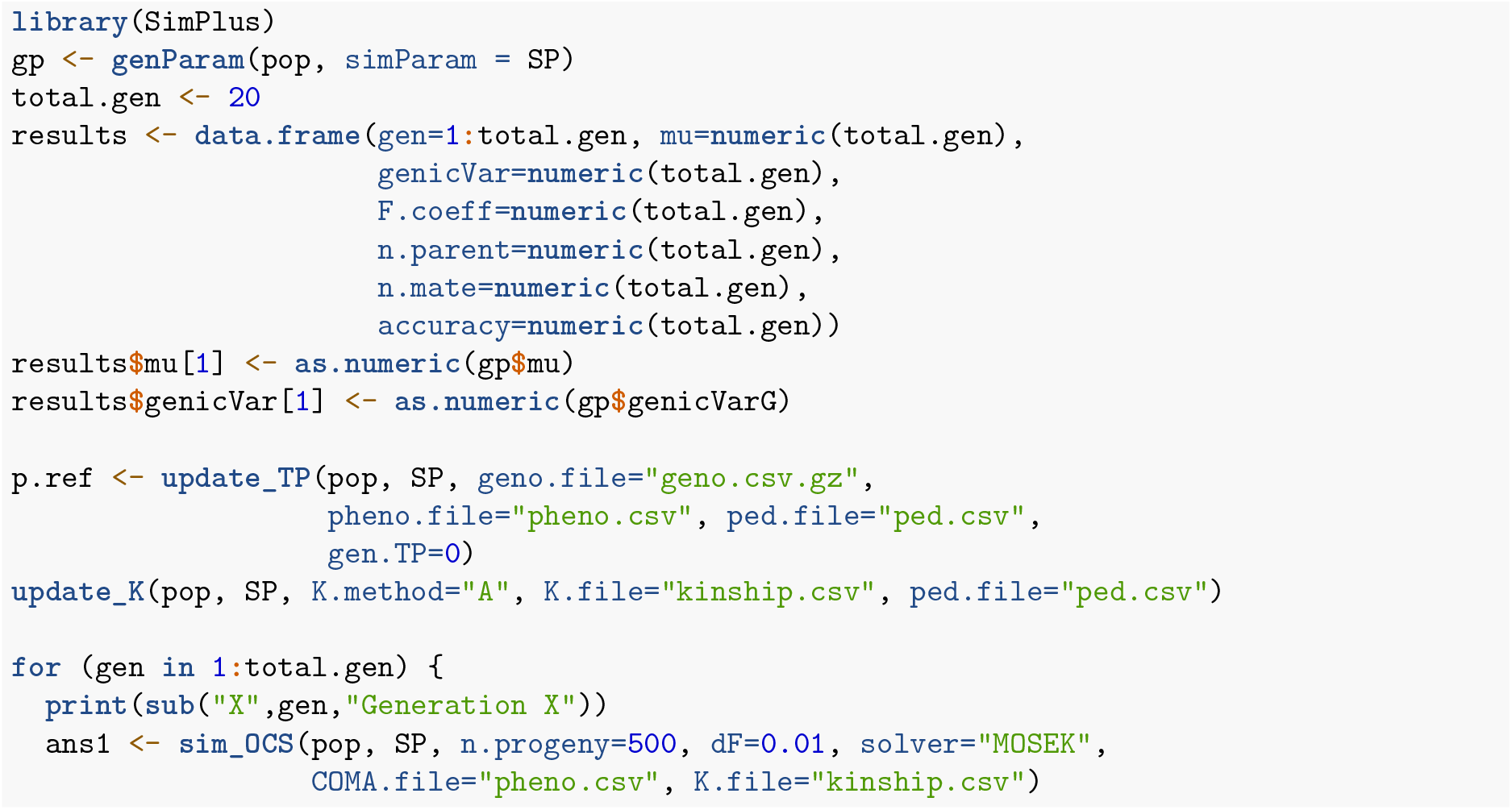
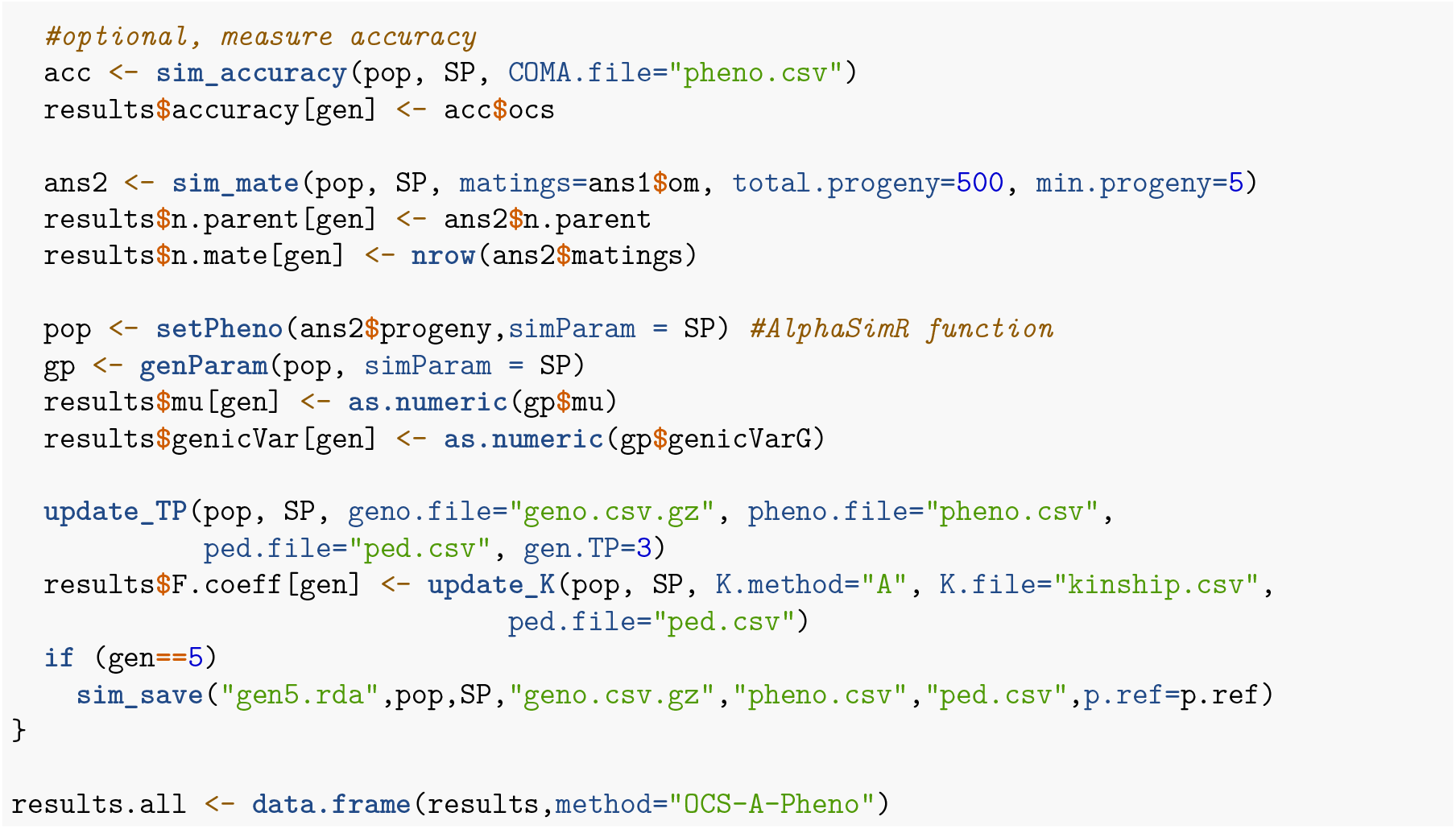

Even though there is no genomic selection in the above simulation, the function `update_TP` is still required to update the phenotype and pedigree files for OCS. The argument `gen.TP` controls the number of generations in the TP (including the candidates, which are selected after phenotyping). It is set at 3 to anticipate the simulation below, ensuring the full TP is available at the onset of GS in generation 5.

### OCS-A-GEBV

To simulate GS requires another function, `sim_StageWise`, to predict the marker effects using the StageWise package. The file with the predicted marker effects (COMA.file) is now used as input for `sim_OCS` instead of the phenotype file.

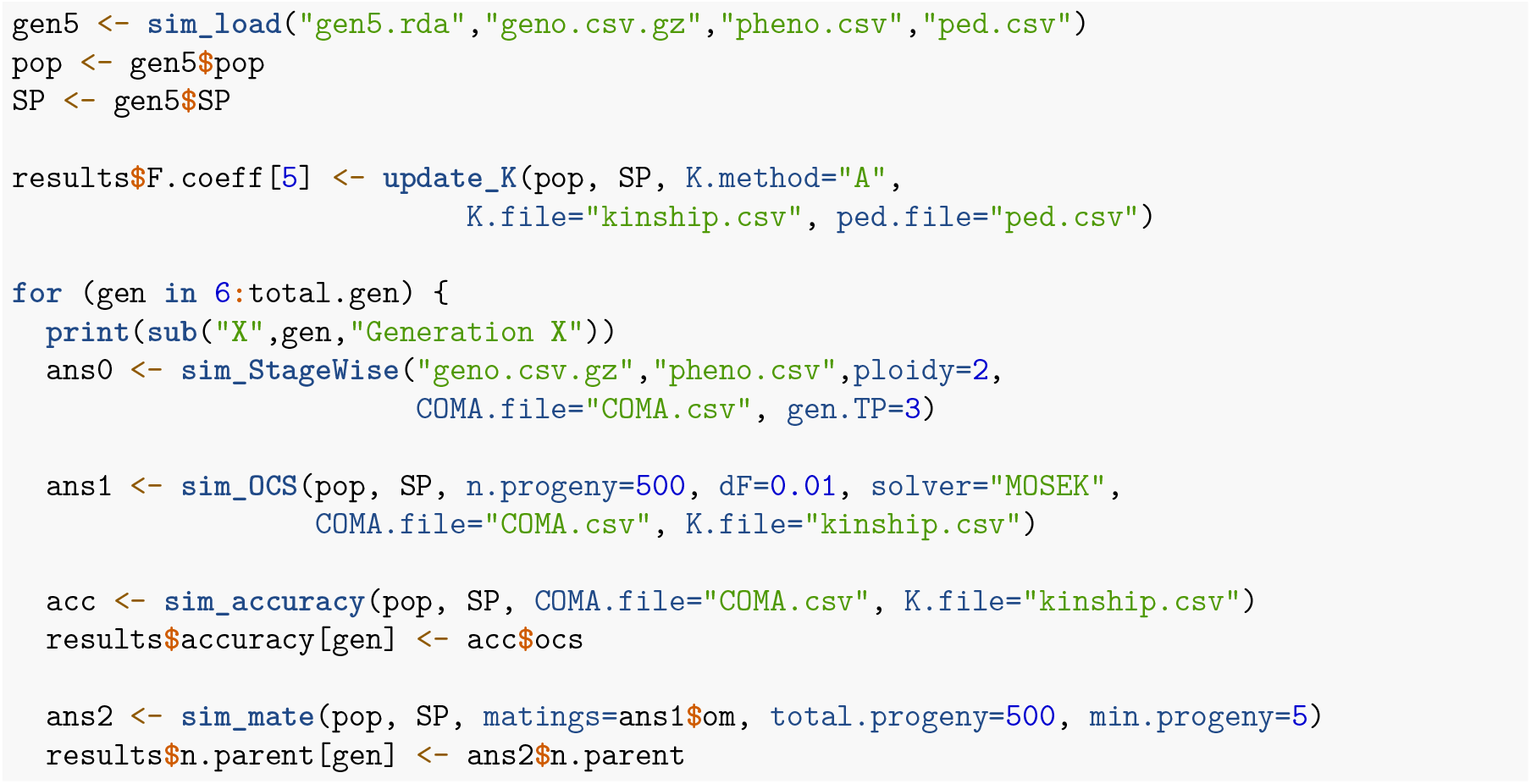
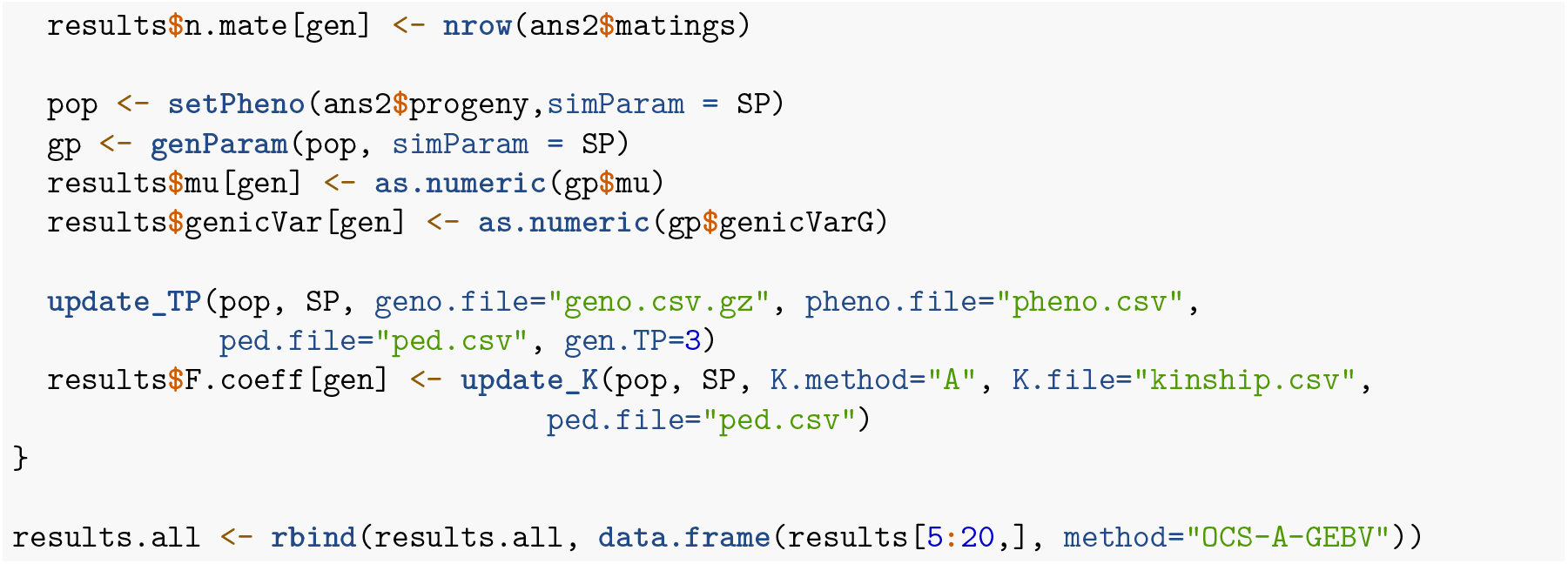

### OCS-G.IBD-GEBV

Instead of pedigree kinship, genomic IBD (G.IBD) kinship can be used to control inbreeding. To reduce computational time, not all segregating sites are needed for this calculation. The parameter “ibd.loci” in the function `update_K` controls how many loci per chromosome to use. The default is 100, and since the chromosomes are 100 cM, this implies 1 marker per cM.

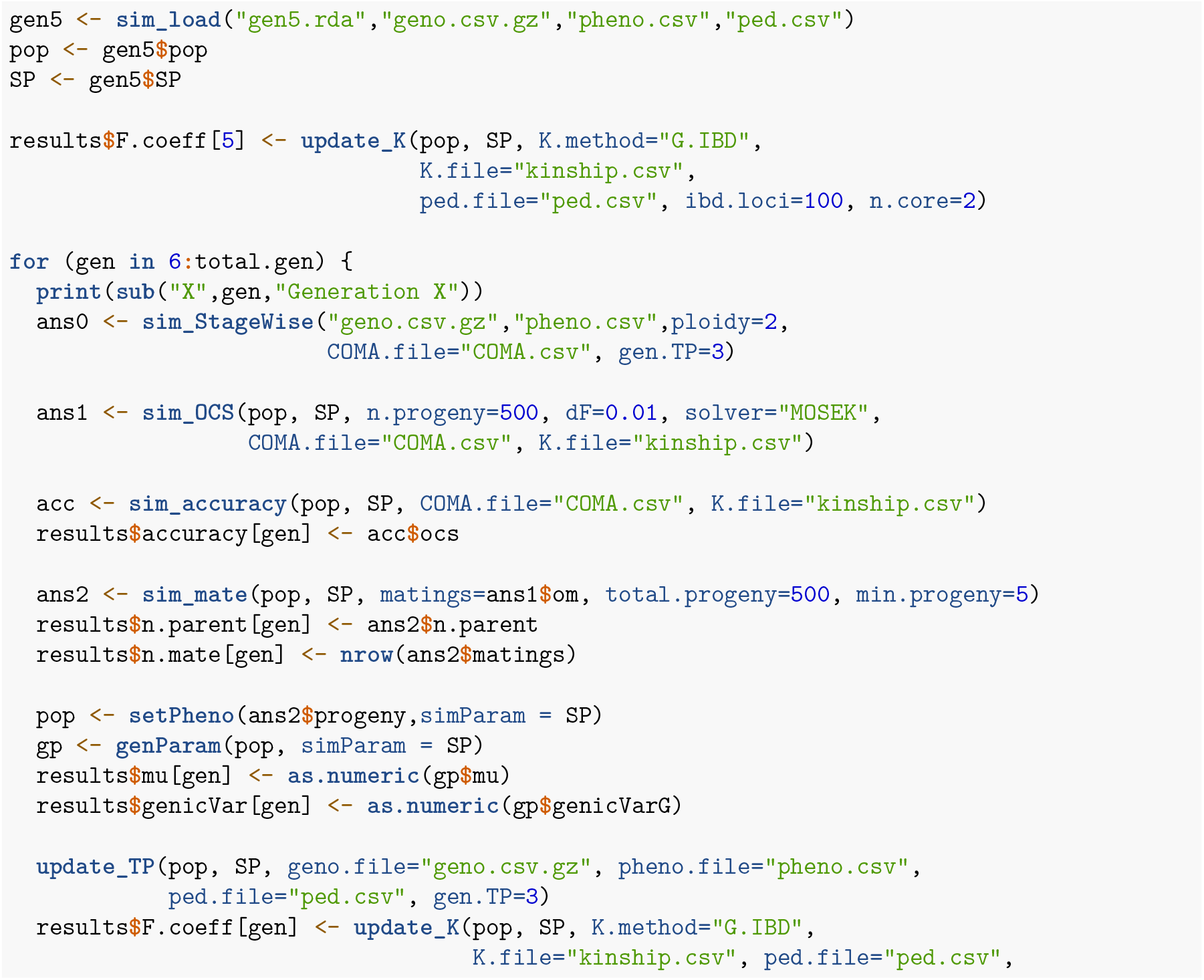
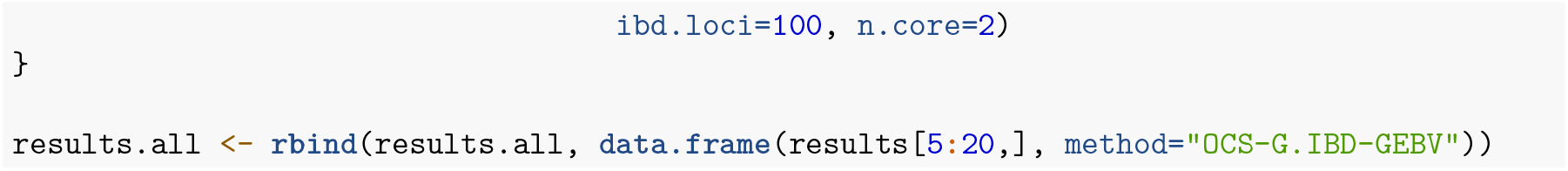

### OMA-A-GPMP

Simulating OMA follows the same code as above but replaces `sim_OCS` with `sim_OMA`, which has an additional argument `max.parent`. To limit the size of the computational problem for OMA, OCS is first used to reduce the number of candidates to `max.parent`. The dF argument for `sim_OMA` is a vector of 2 numbers for the lower and upper bounds. In this case, the numbers are identical to create an equality constraint.

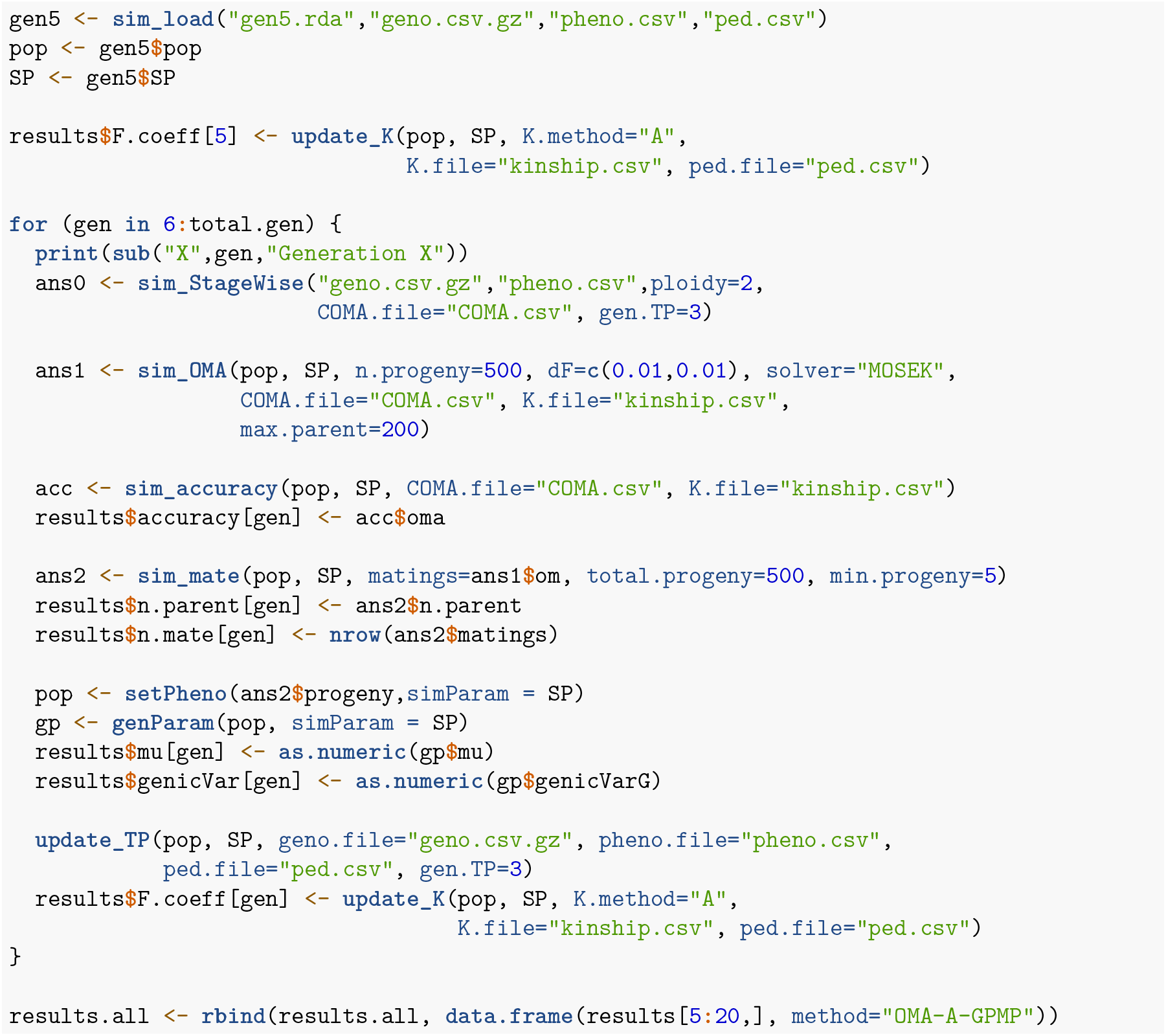

The average inbreeding rate at time *t* follows

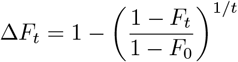

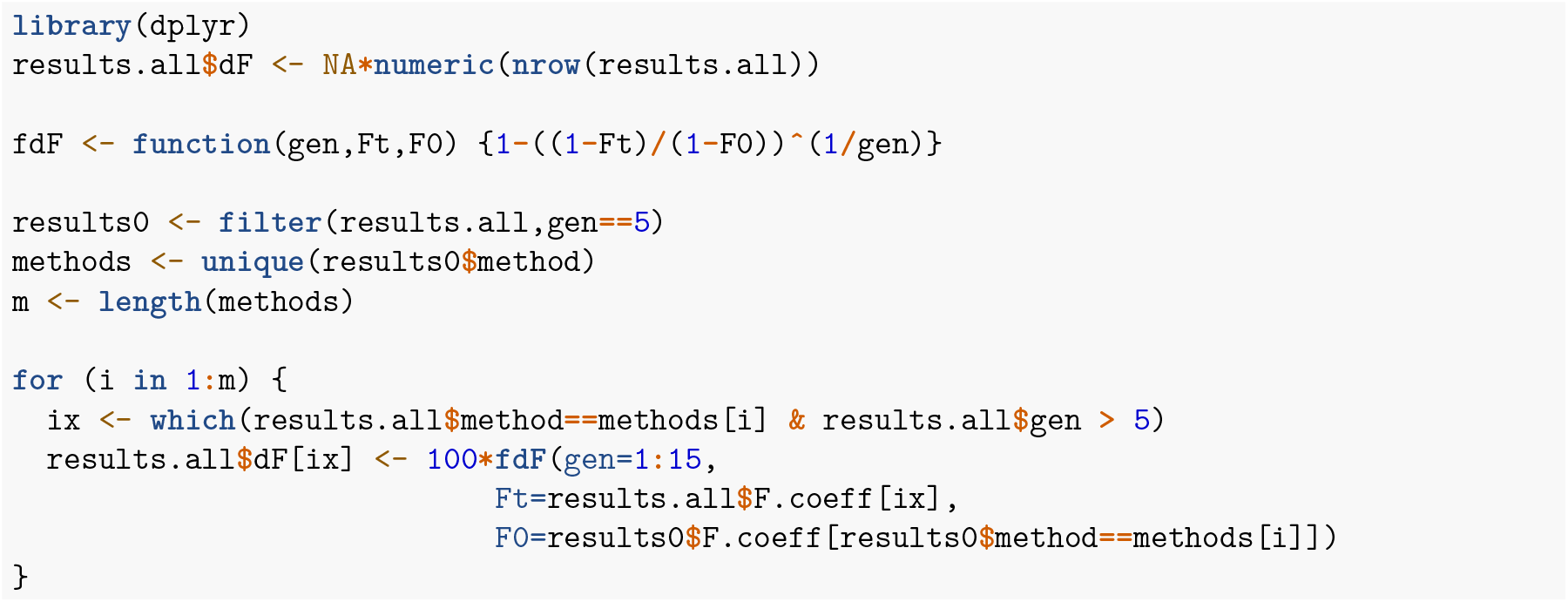

We can now plot the results.

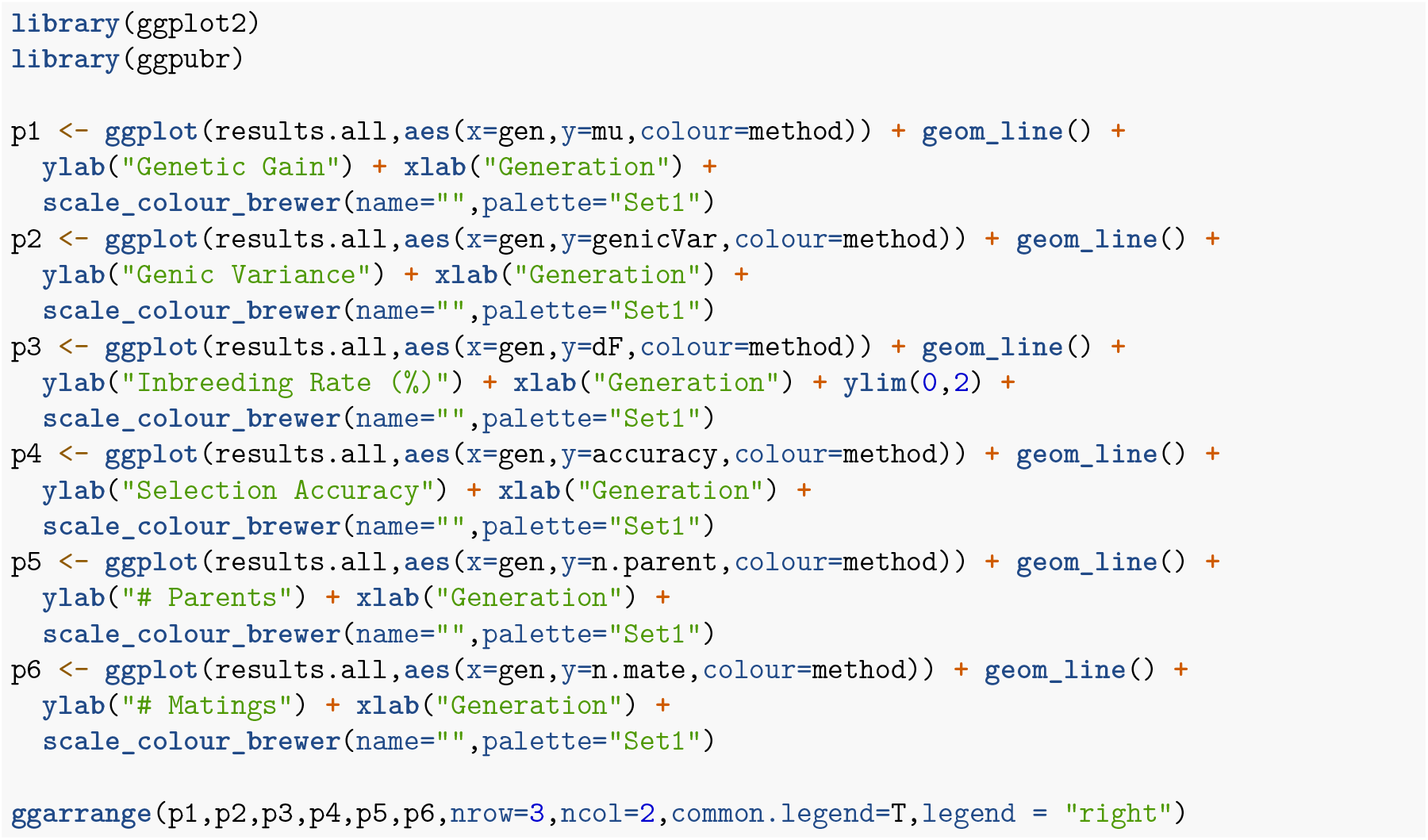
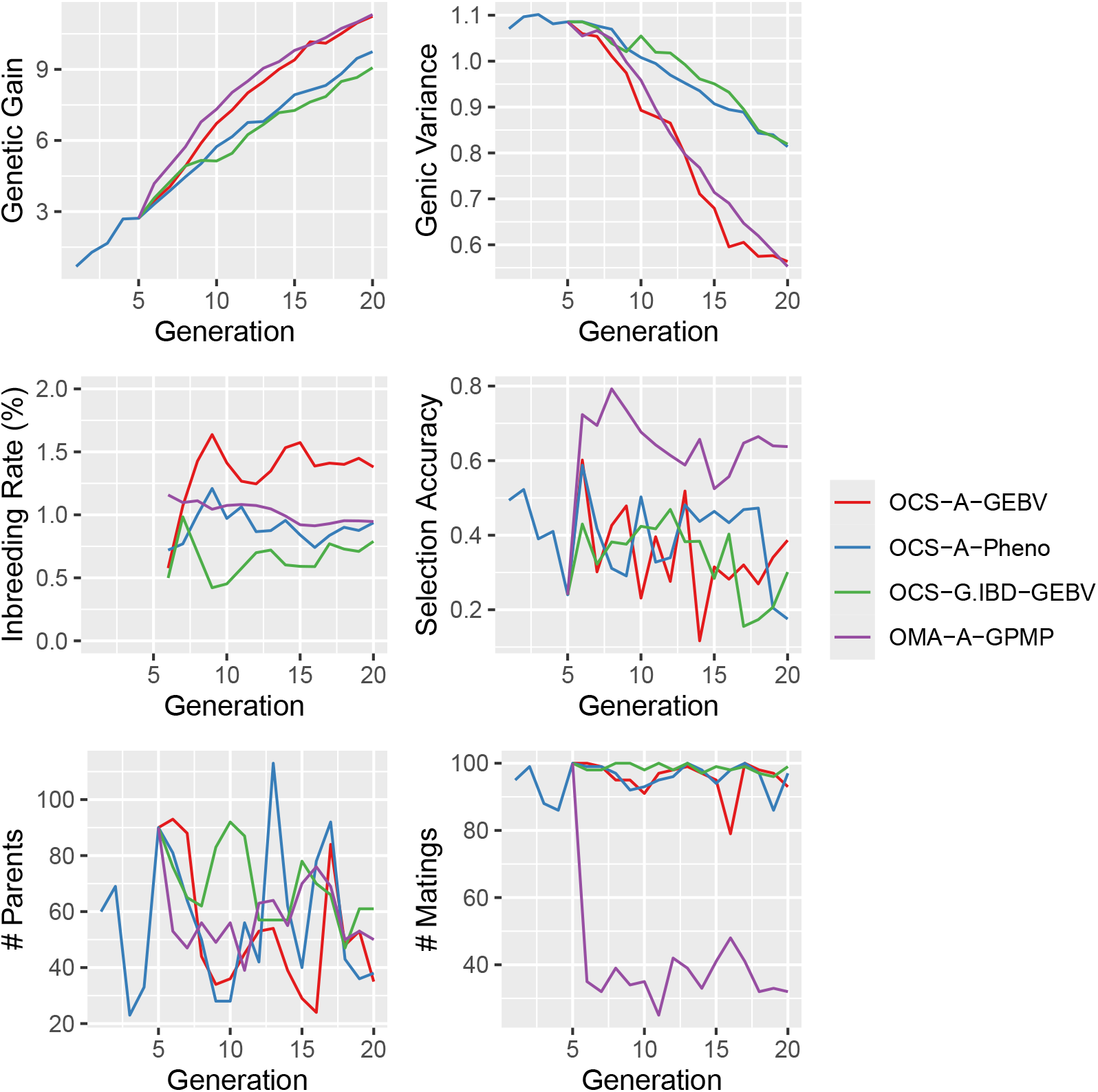

This short simulation illustrates some key points, which are discussed in greater depth in the publication. Compared to OCS, OMA has higher accuracy for predicting mate performance and sparser mating designs. OMA generated more gain in the short term, but long-term gains were similar.

## Package ‘SimPlus’

October 4, 2024

**Title** Functions to enhance AlphaSimR

**Version** 0.06

**Author** Jeffrey B. Endelman

**Maintainer** Jeffrey Endelman

**Description** Functions to enhance AlphaSimR

**Depends** R (>= 4.1)

**License** GPL-3

**LazyData** true

**RoxygenNote** 7.2.3

**Encoding** UTF-8

**Imports** utils, AlphaSimR, dplyr, polyBreedR, pedigree, data.table, COMA

**Suggests** knitr, rmarkdown, StageWise

**VignetteBuilder** knitr

### Index 8

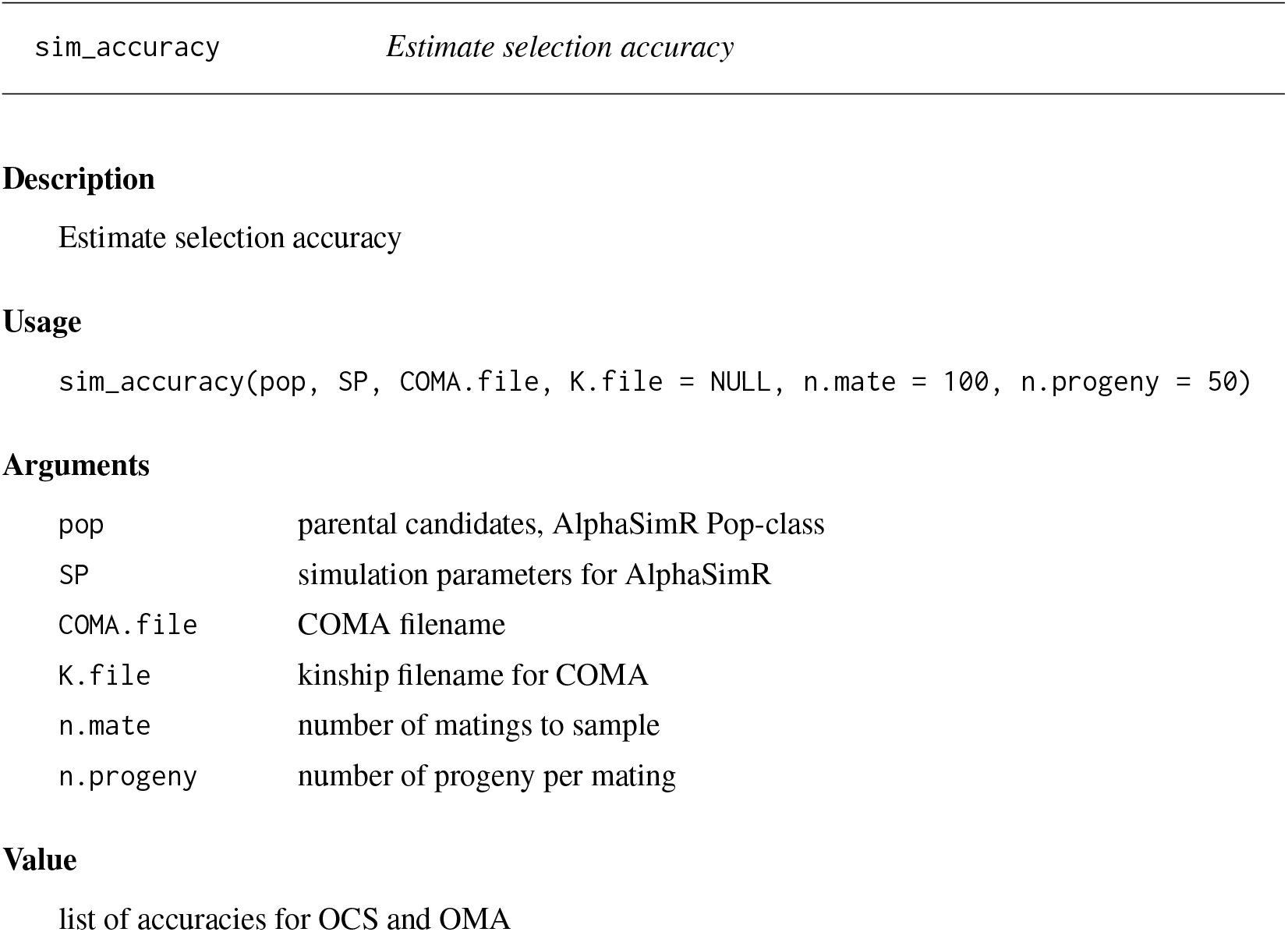

|  |  |
| --- | --- |
| sim_accuracy | *Estimate selection accuracy* |
| **Arguments** |  |
| pop | parental candidates, AlphaSimR Pop-class |
| SP | simulation parameters for AlphaSimR |
| COMA.file | COMA filename |
| K.file | kinship filename for COMA |
| n.mate | number of matings to sample |
| n.progeny | number of progeny per mating |

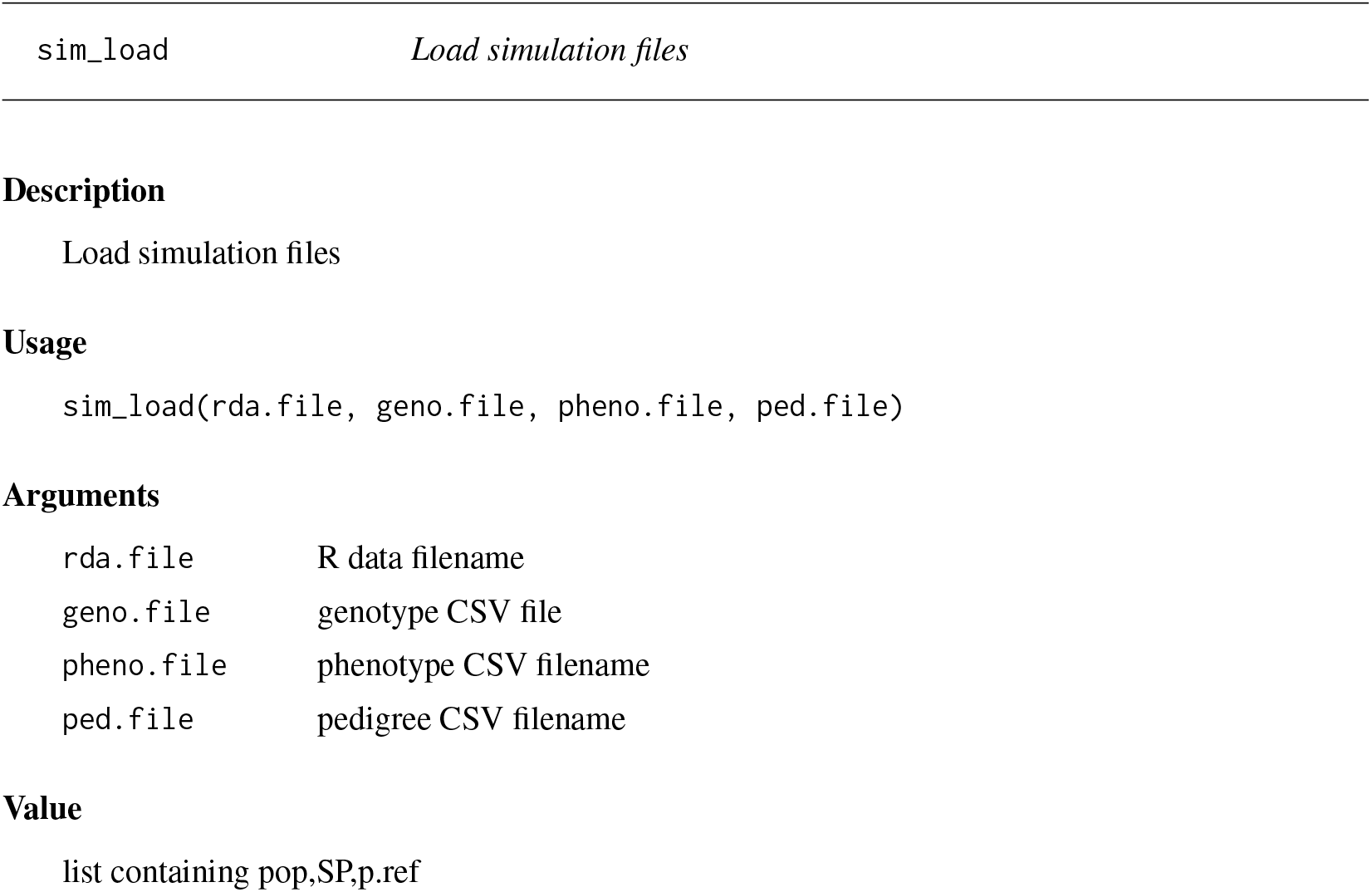

|  |  |
| --- | --- |
| sim_load | *Load simulation files* |
| **Arguments** |  |
| rda.file | R data filename |
| geno.file | genotype CSV file |
| pheno.file | phenotype CSV filename |
| ped.file | pedigree CSV filename |

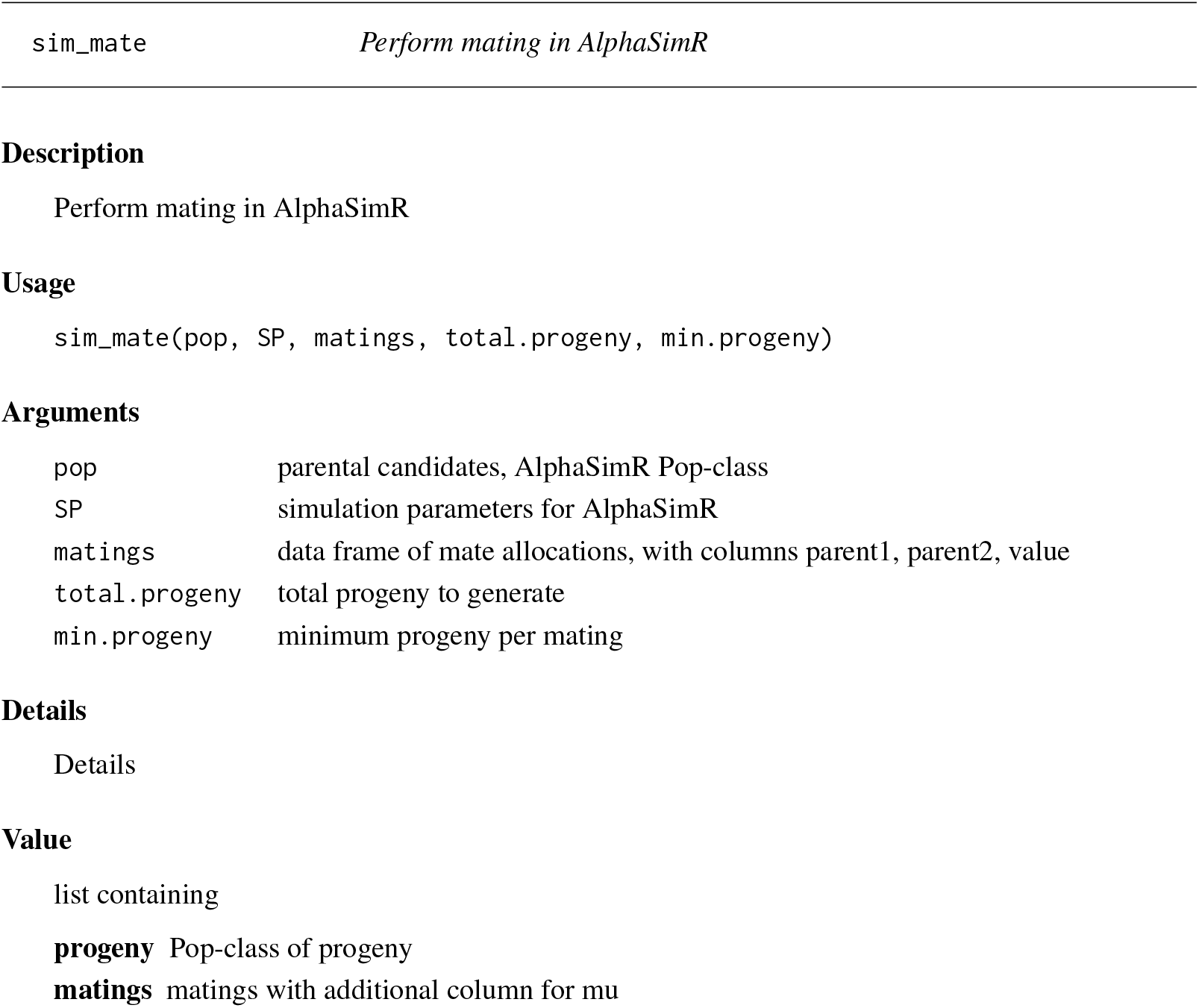

|  |  |
| --- | --- |
| sim_mate | *Perform mating in AlphaSimR* |
| Arguments |  |
| pop | parental candidates, AlphaSimR Pop-class |
| SP | simulation parameters for AlphaSimR |
| matings | data frame of mate allocations, with columns parent1, parent2, value |
| total.progeny | total progeny to generate |
| min.progeny | minimum progeny per mating |

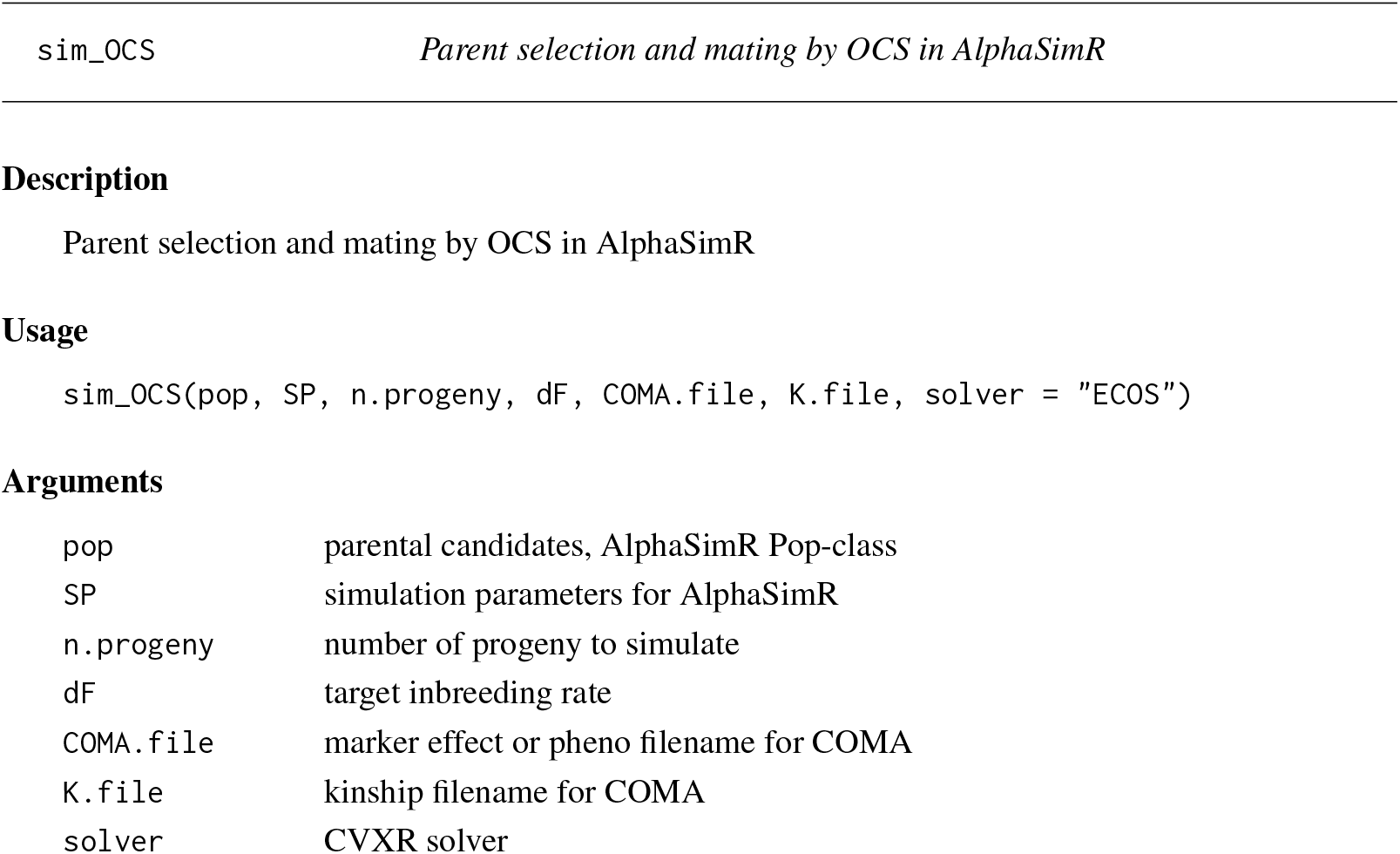
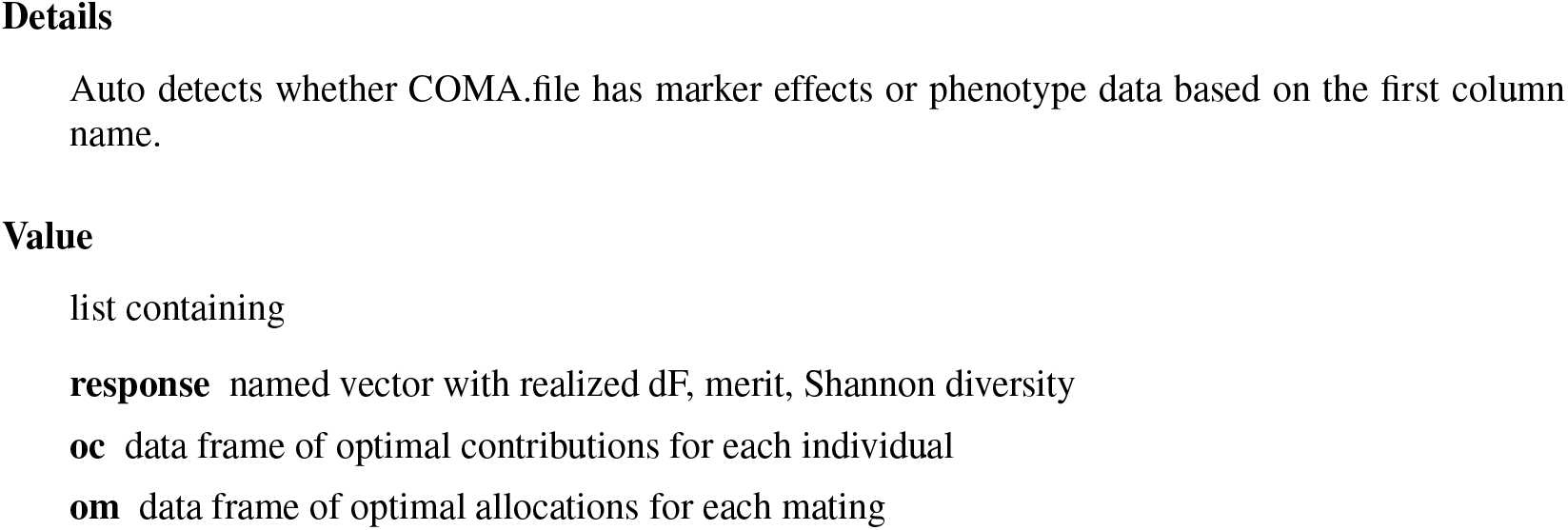

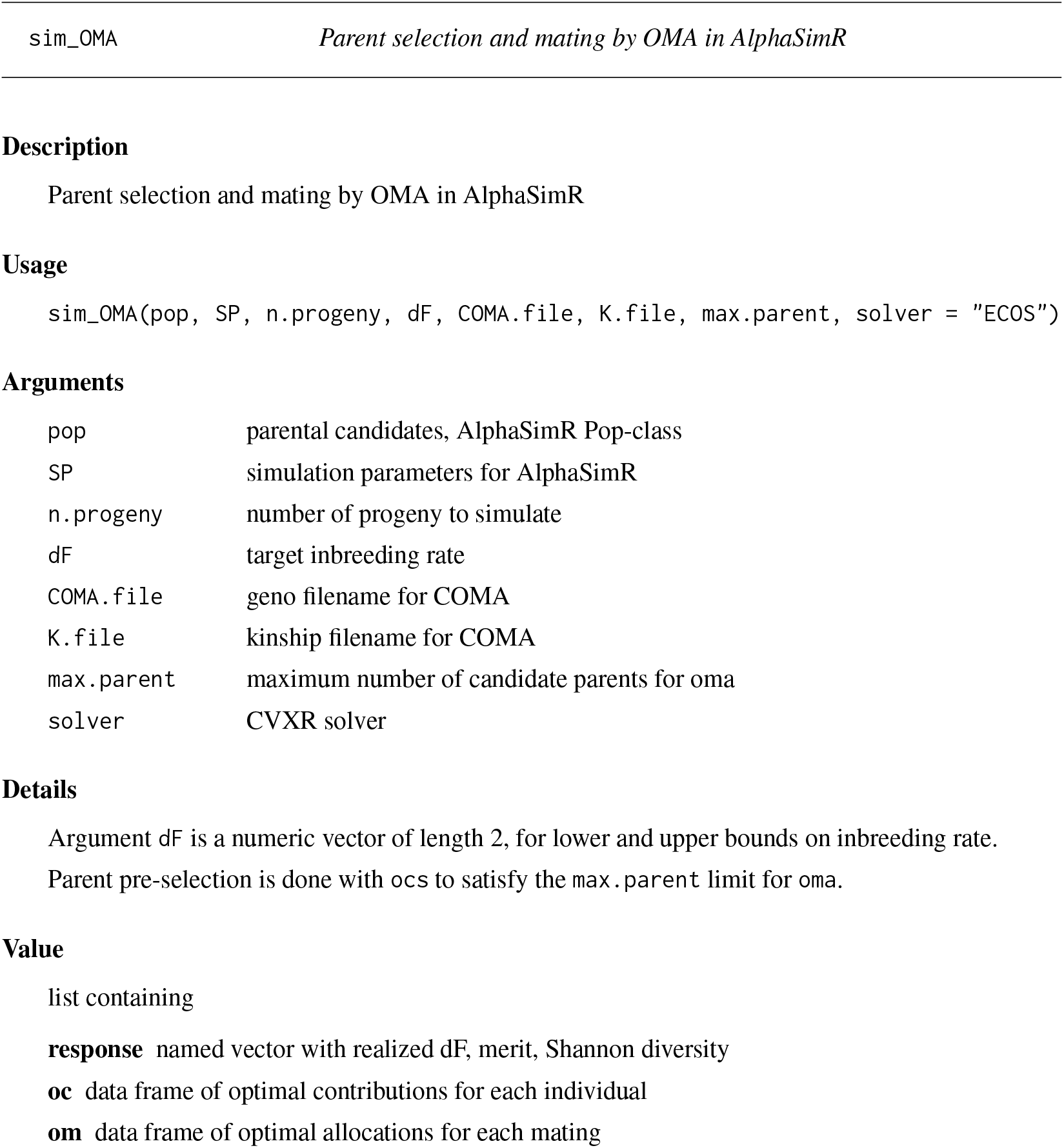

|  |  |
| --- | --- |
| sim_OMA | *Parent selection and mating by OMA in AlphaSimR* |
| **Arguments** |  |
| pop | parental candidates, AlphaSimR Pop-class |
| SP | simulation parameters for AlphaSimR |
| n.progeny | number of progeny to simulate |
| dF | target inbreeding rate |
| COMA.file | geno filename for COMA |
| K.file | kinship filename for COMA |
| max.parent | maximum number of candidate parents for oma |
| solver | CVXR solver |

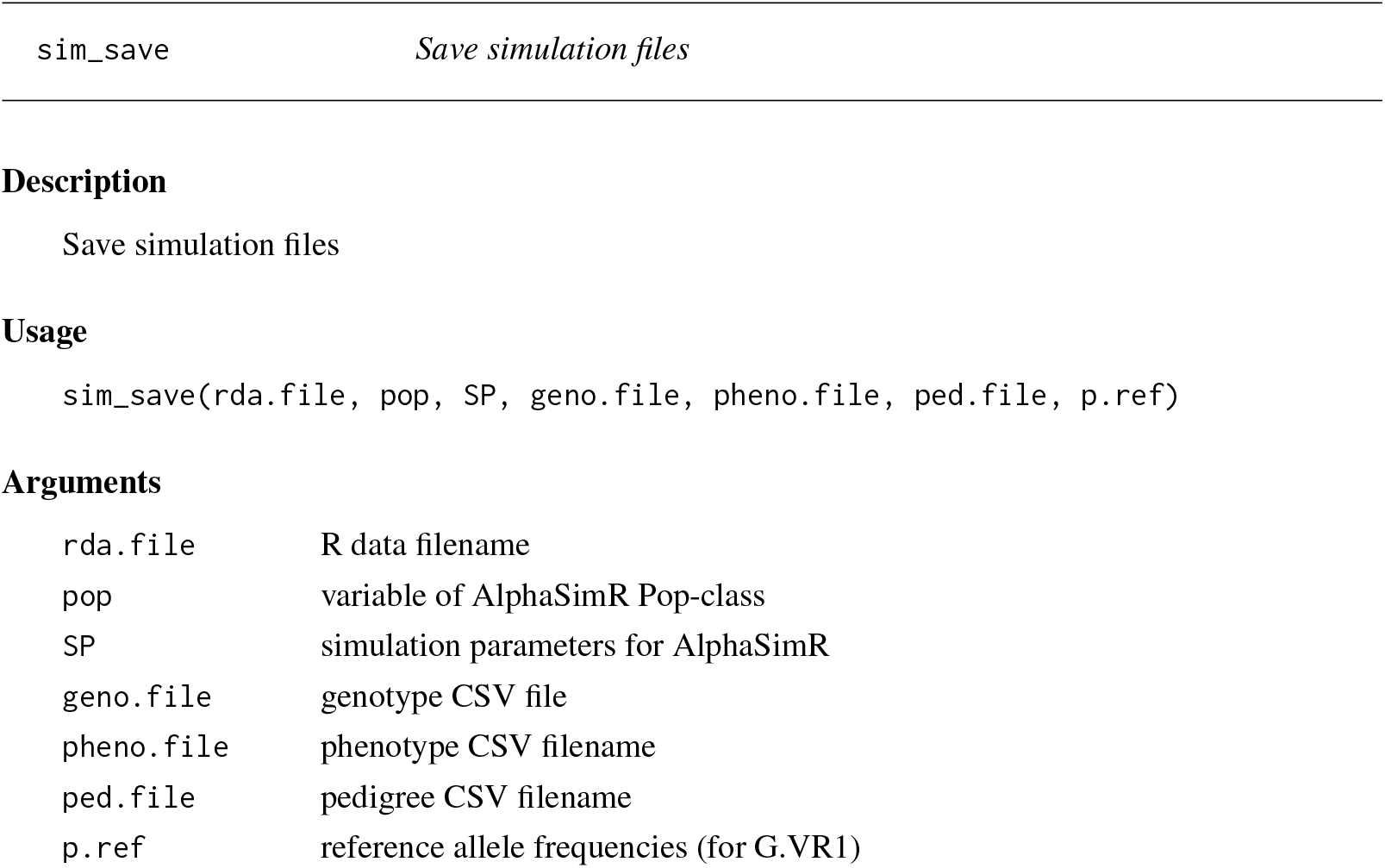

|  |  |
| --- | --- |
| sim_save | *Save simulation files* |
| **Arguments** |  |
| rda.file | R data filename |
| pop | variable of AlphaSimR Pop-class |
| SP | simulation parameters for AlphaSimR |
| geno.file | genotype CSV file |
| pheno.file | phenotype CSV filename |
| ped.file | pedigree CSV filename |
| p.ref | reference allele frequencies (for G.VR1) |

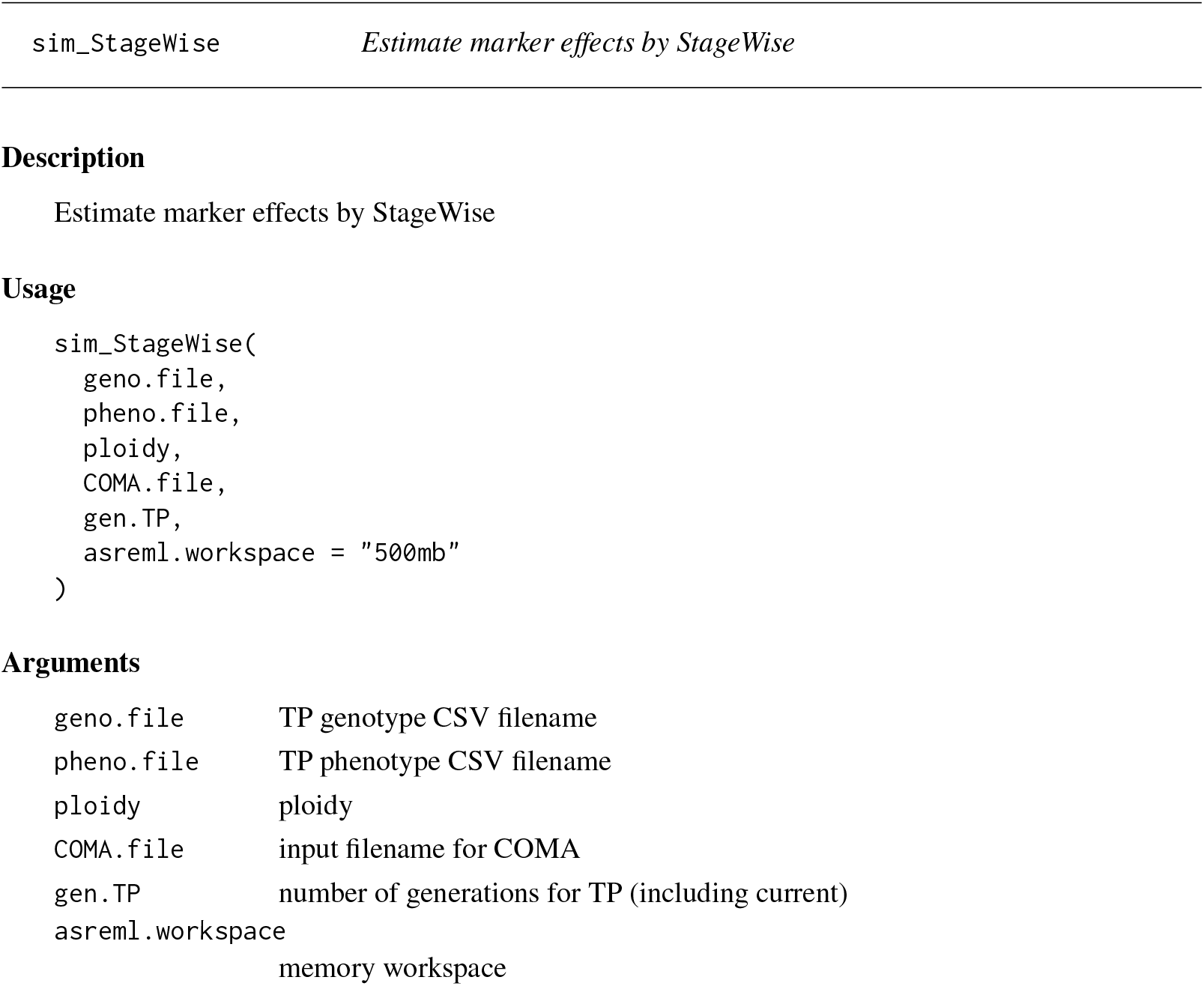
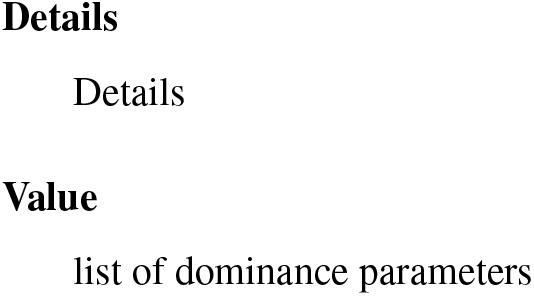

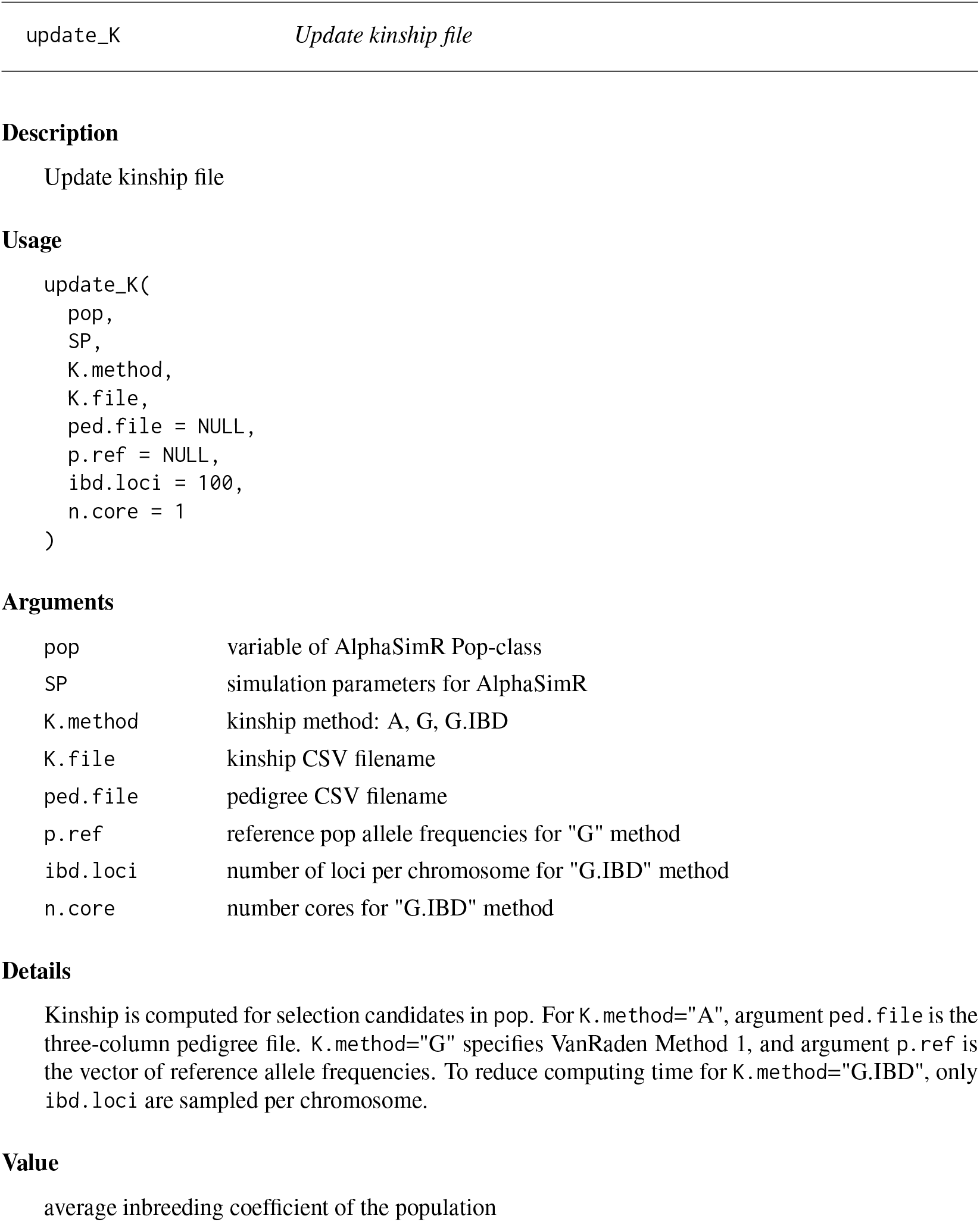

|  |  |
| --- | --- |
| update_K | *Update kinship file* |
| **Arguments** |  |
| pop | variable of AlphaSimR Pop-class |
| SP | simulation parameters for AlphaSimR |
| K.method | kinship method: A, G, G.IBD |
| K.file | kinship CSV filename |
| ped.file | pedigree CSV filename |
| p.ref | reference pop allele frequencies for "G" method |
| ibd.loci | number of loci per chromosome for "G.IBD" method |
| n.core | number cores for "G.IBD" method |

**Arguments**
|  |  |
| --- | --- |
| pop | variable of AlphaSimR Pop-class |
| SP | simulation parameters for AlphaSimR |
| geno.file | genotype CSV file |
| pheno.file | phenotype CSV filename |
| ped.file | pedigree CSV filename |
| gen.TP | # generations recorded for the TP (including current) |

## Index

`sim_accuracy`, 2

`sim_load`, 2

`sim_mate`, 3

`sim_OCS`, 3

`sim_OMA`, 4

`sim_save`, 5

`sim_StageWise`, 5

`update_K`, 6

`update_TP`, 7

## File S5: Potato Marker Effects

Jeff Endelman 2024-07-27

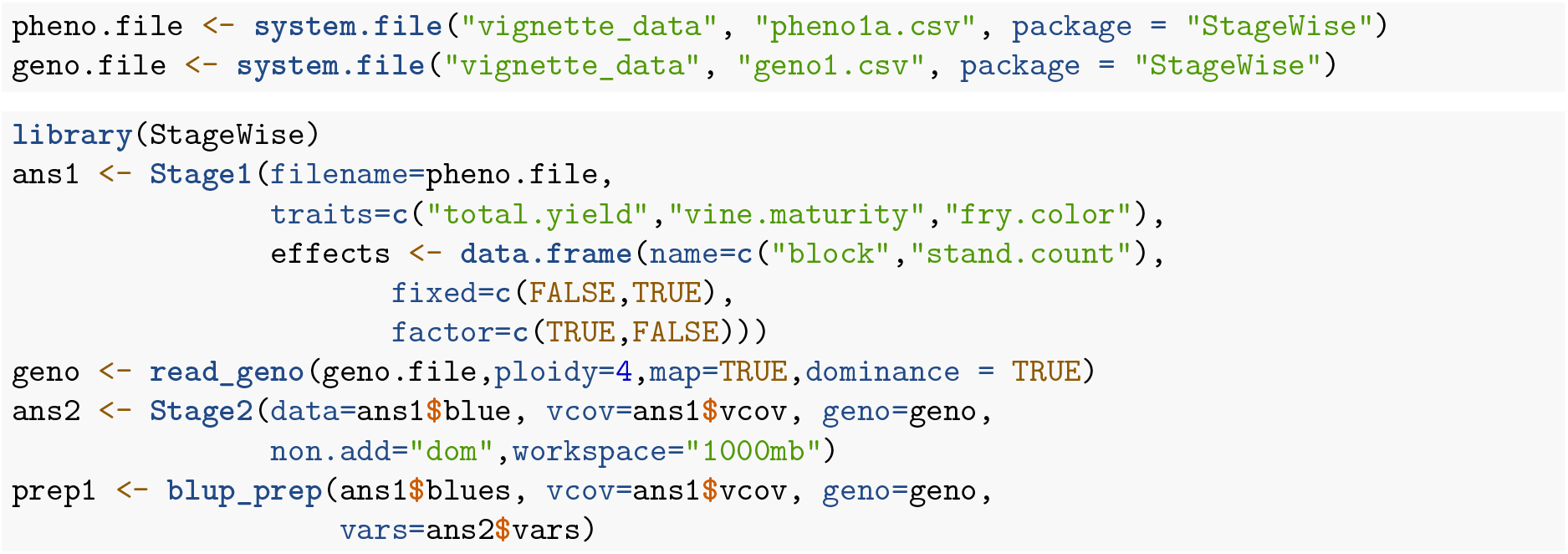

Yield and fry color have similar importance as breeding objectives, while later maturity is undesirable. Initially, a restricted index was calculated, with equal merit for yield and fry color and zero gain for maturity.

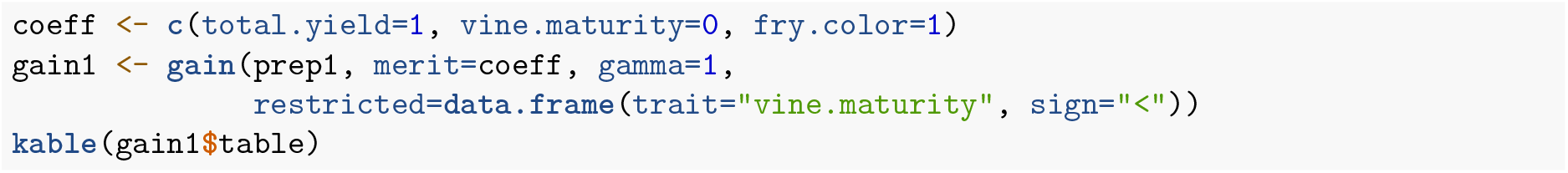

| trait | response | index |
| --- | --- | --- |
| total.yield | 0.256 | 0.636 |
| vine.maturity | 0.000 | -0.436 |
| fry.color | 0.436 | 0.636 |

Due to the unfavorable correlation between yield and late maturity, the yield response was only 65% of the response for fry color. As an alternative, a desired gains index was calculated, based on equal gains for yield and fry color and zero gain for maturity.

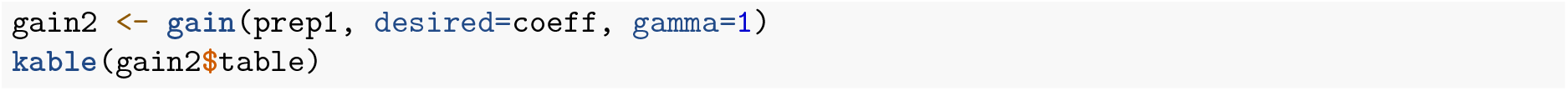

| trait | response | index |
| --- | --- | --- |
| total.yield | 0.34 | 0.702 |
| vine.maturity | 0.00 | -0.459 |
| fry.color | 0.34 | 0.544 |

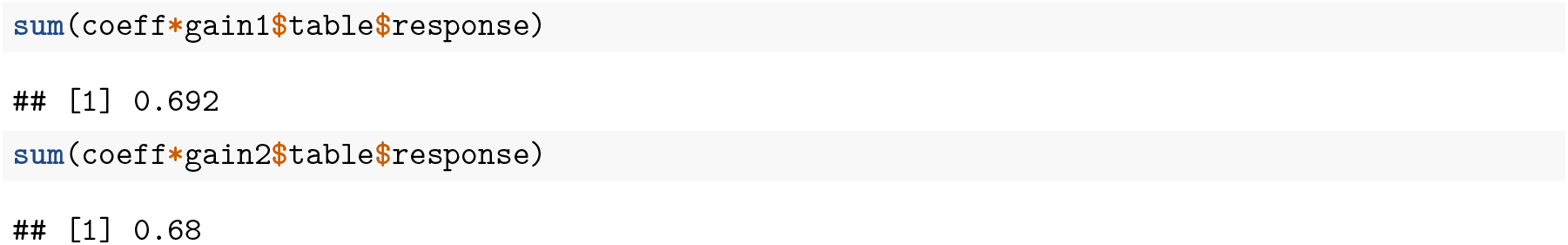

Imposing equal gains reduced total merit by less than 1%, and given the uncertainty of the true merit coefficients, this index was selected.

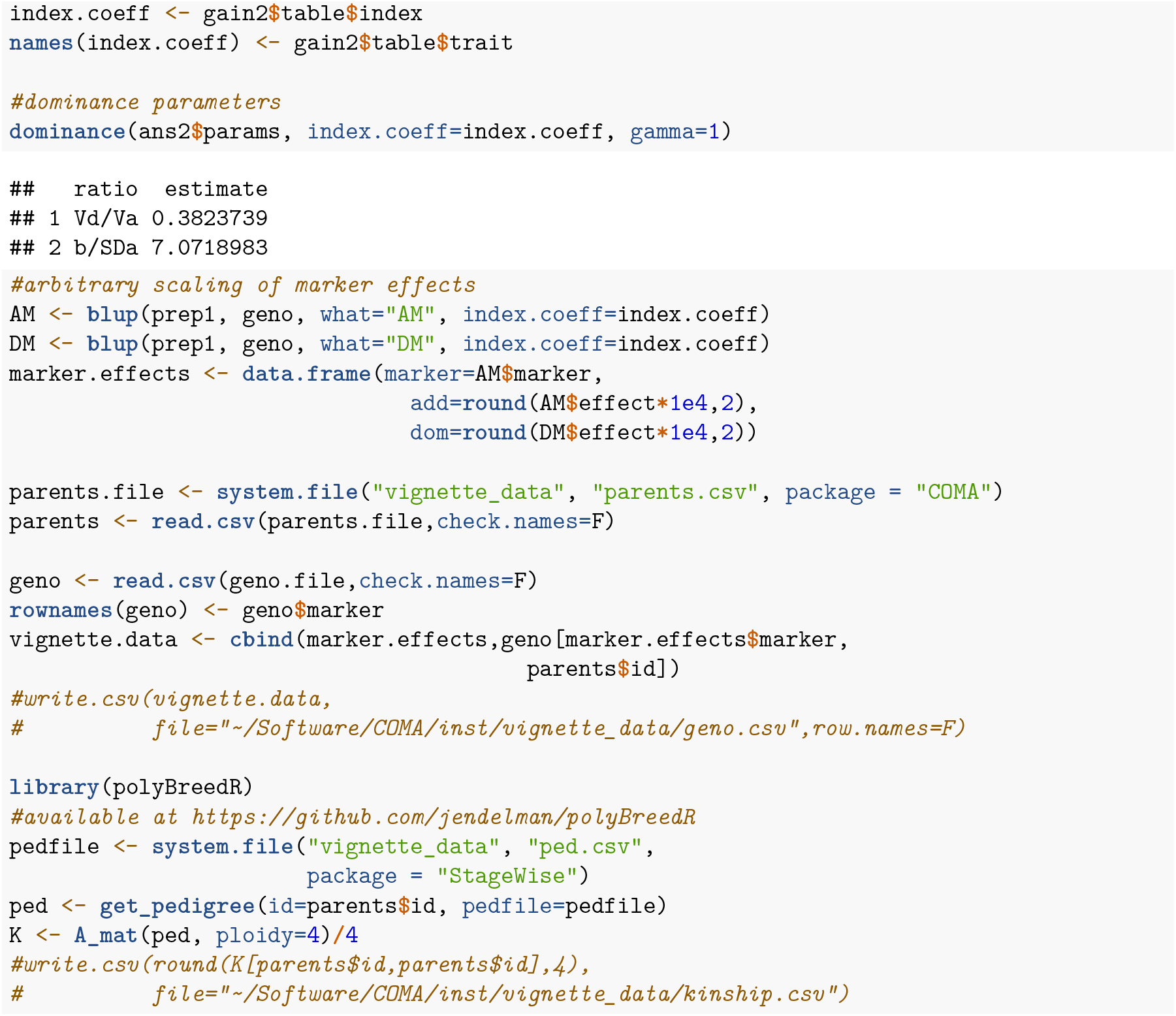

## File S6: R Code for Figures

Jeff Endelman 2024-10-06

**Figure 1**

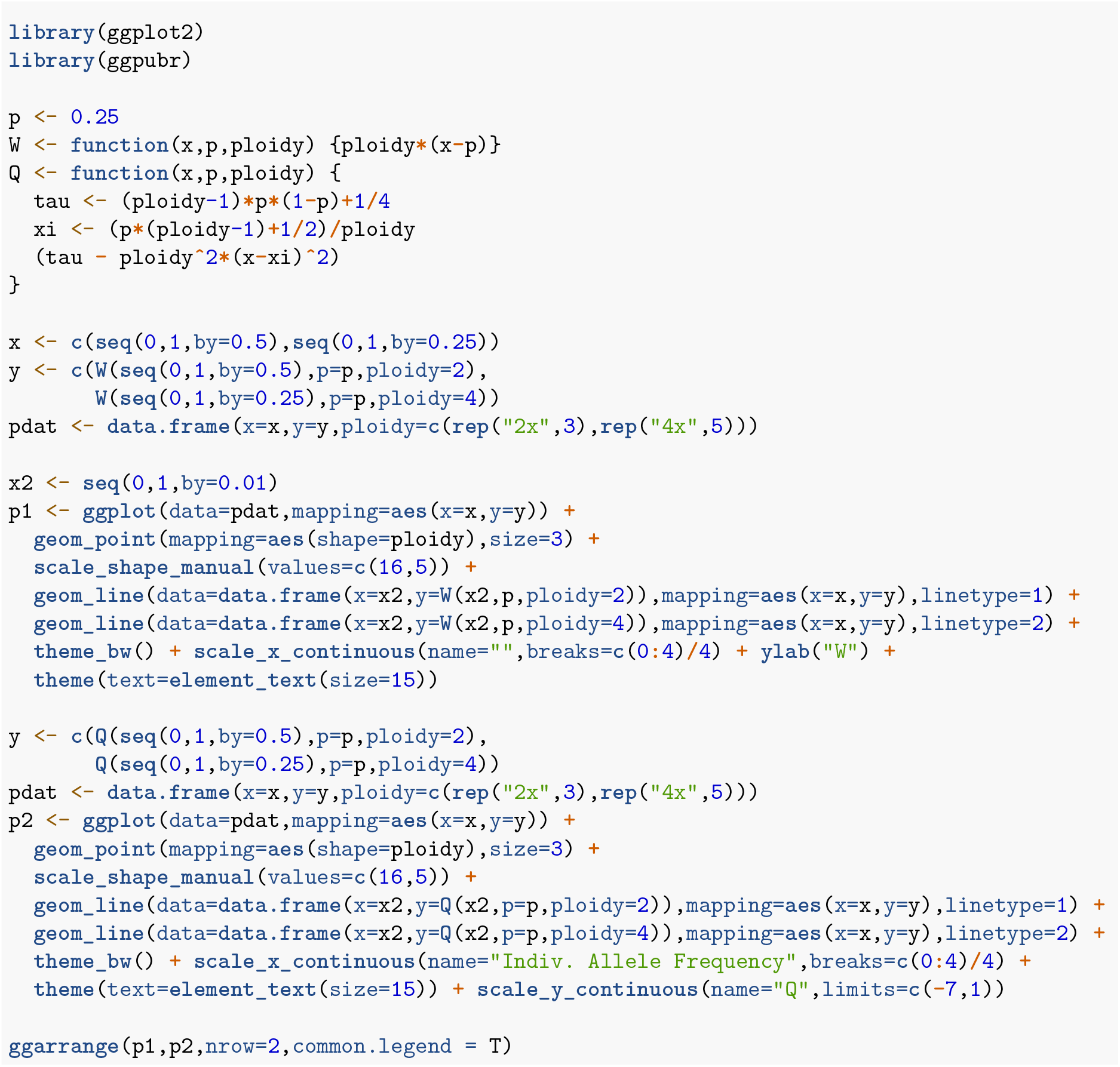
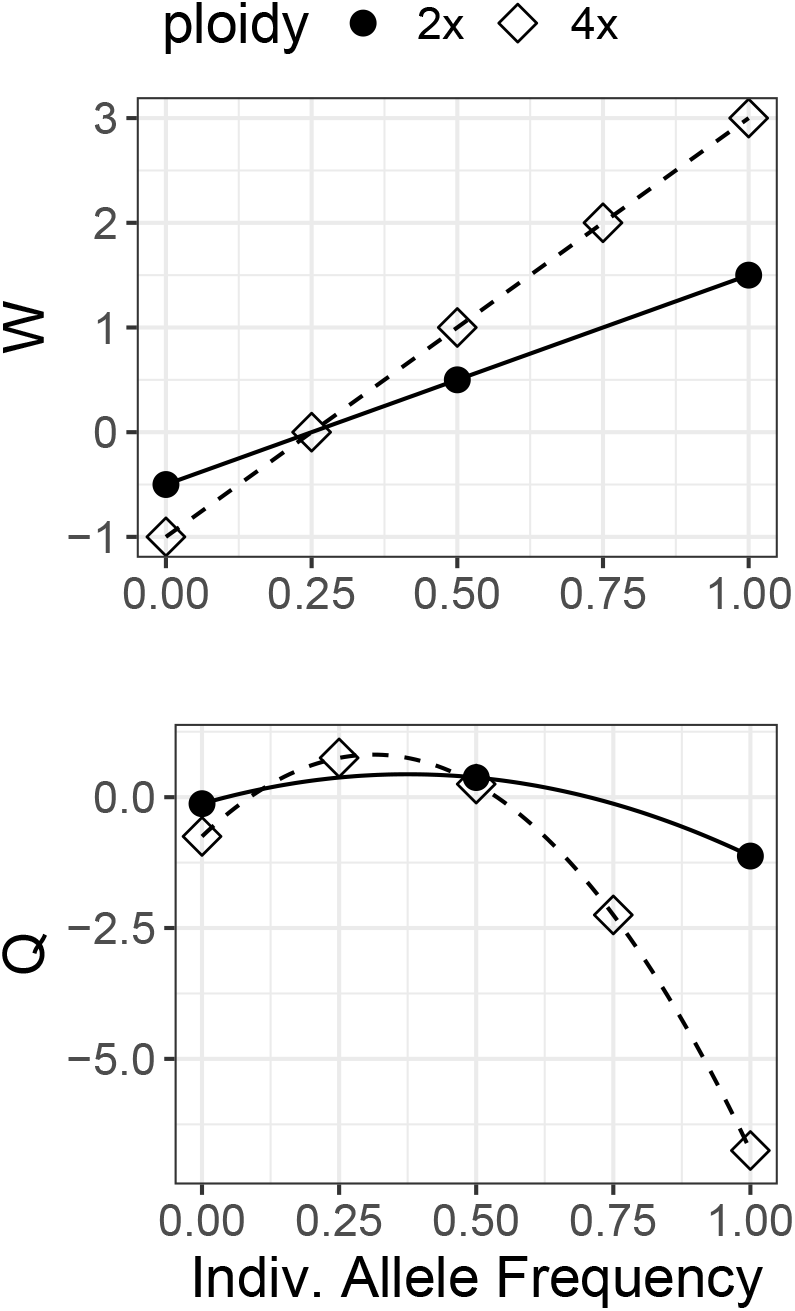

**Figure 2**

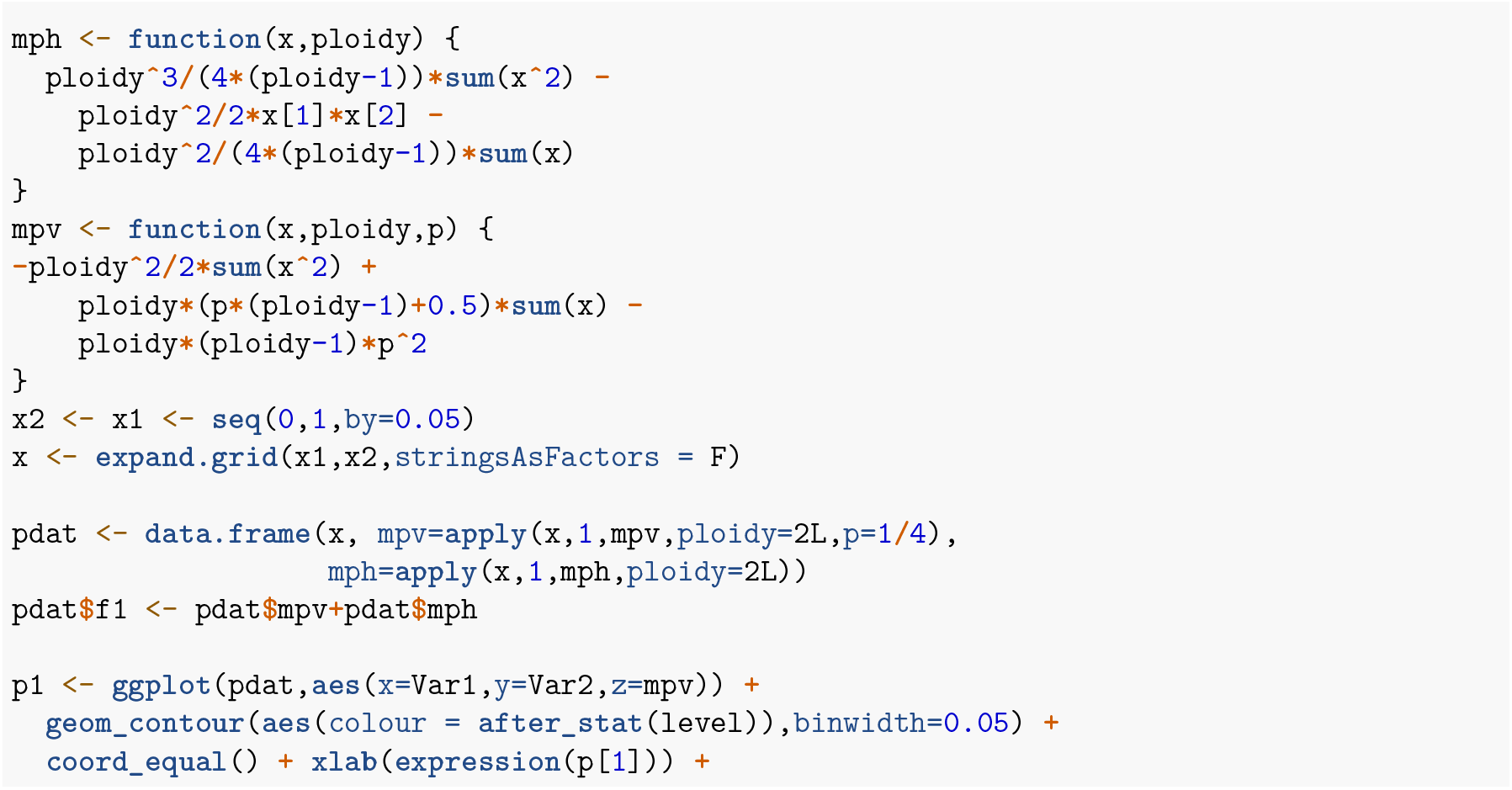
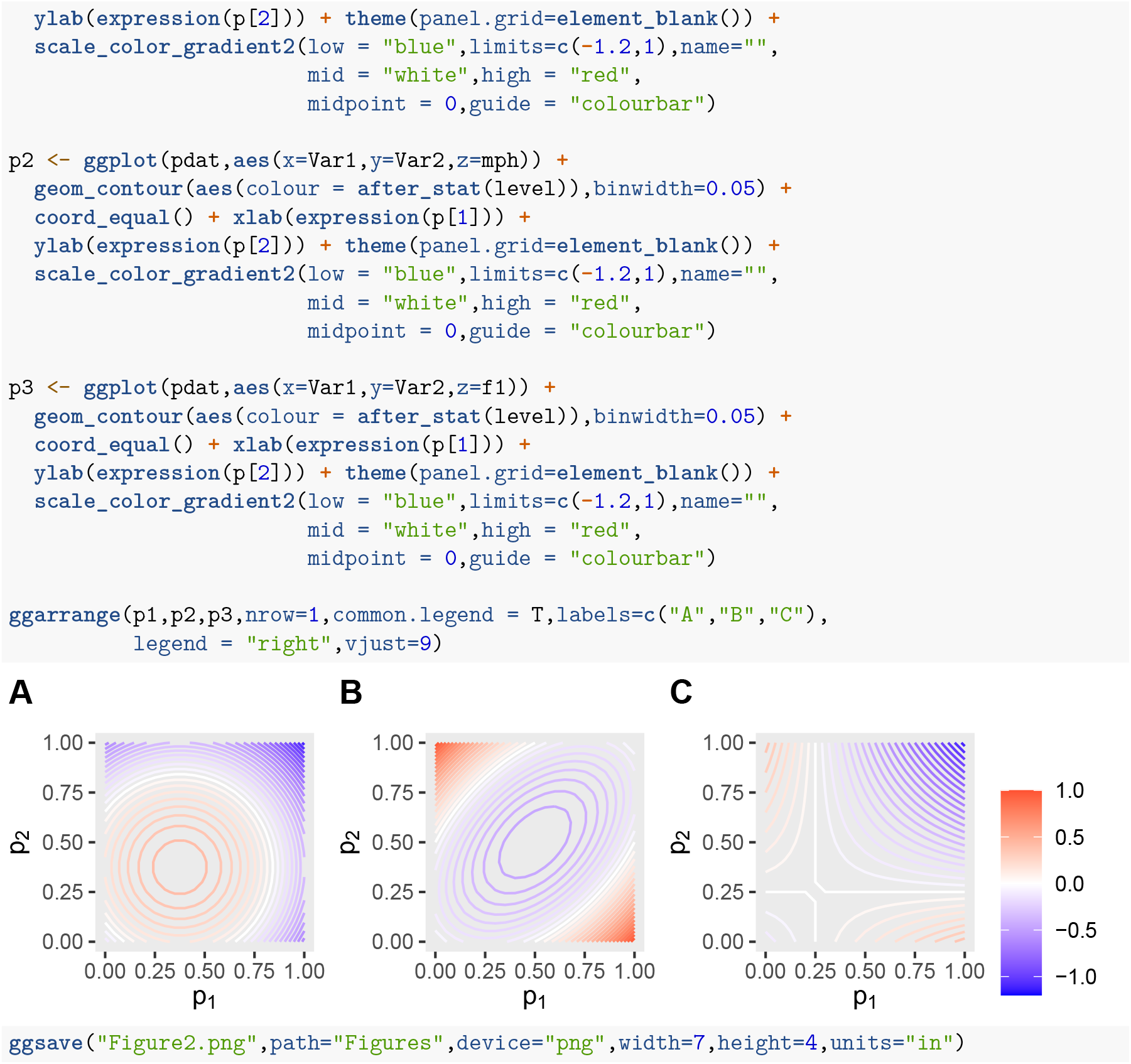

**Figure S1**

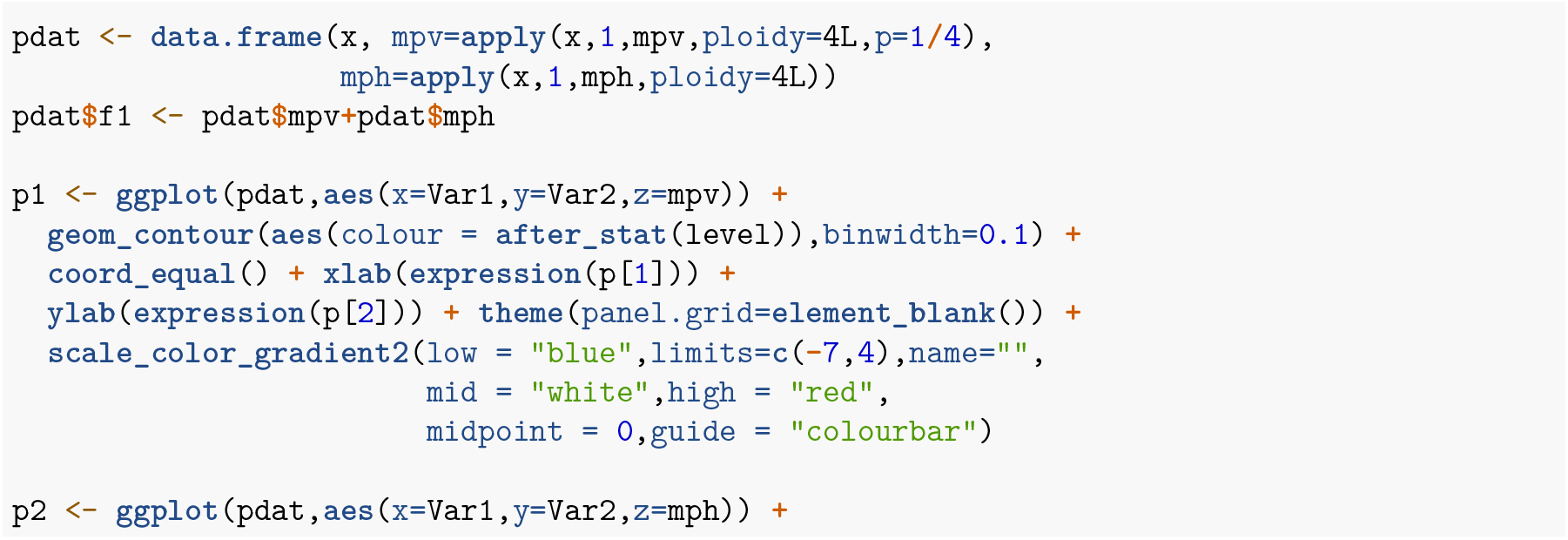
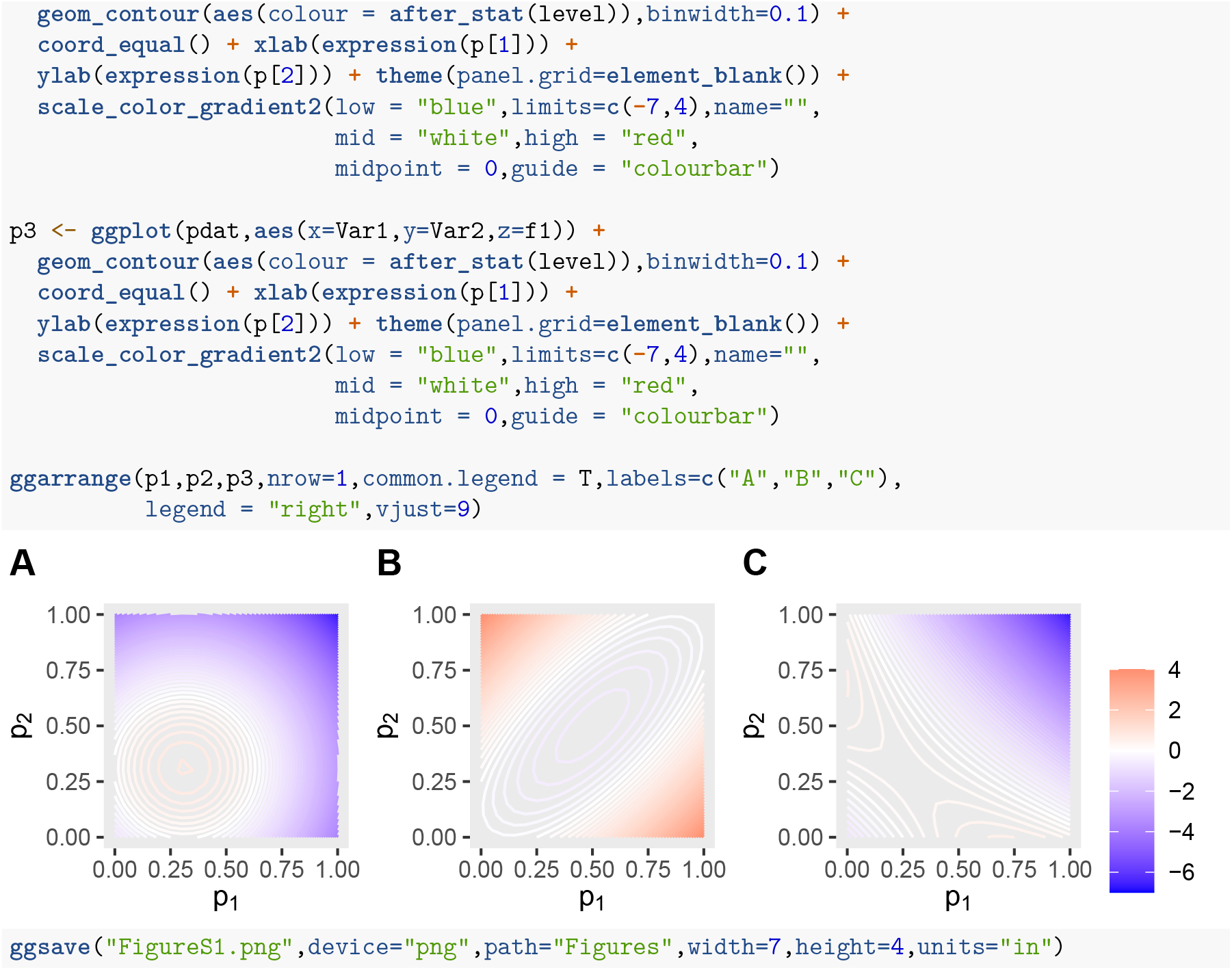

### Prepare simulation data for plotting

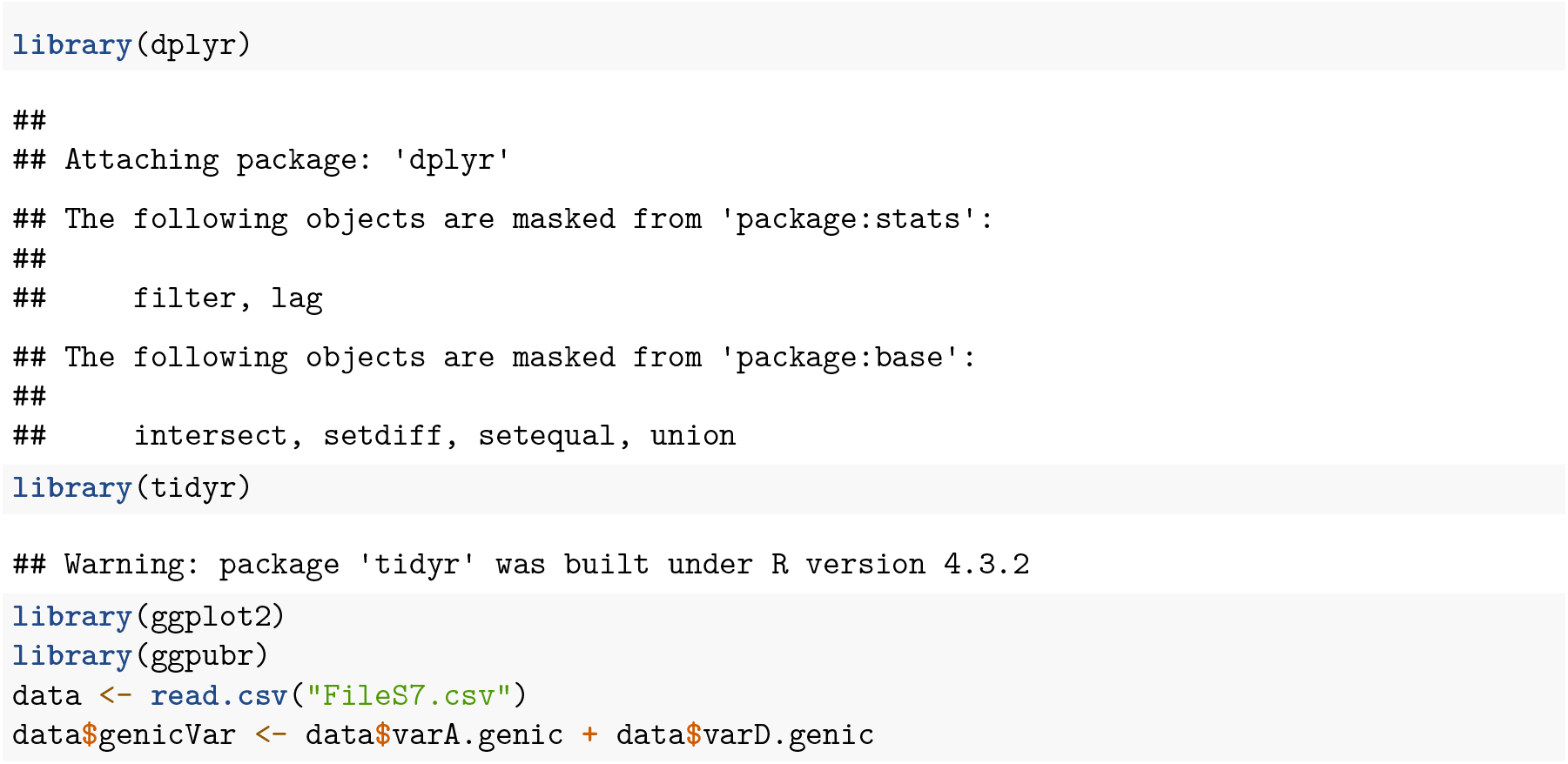
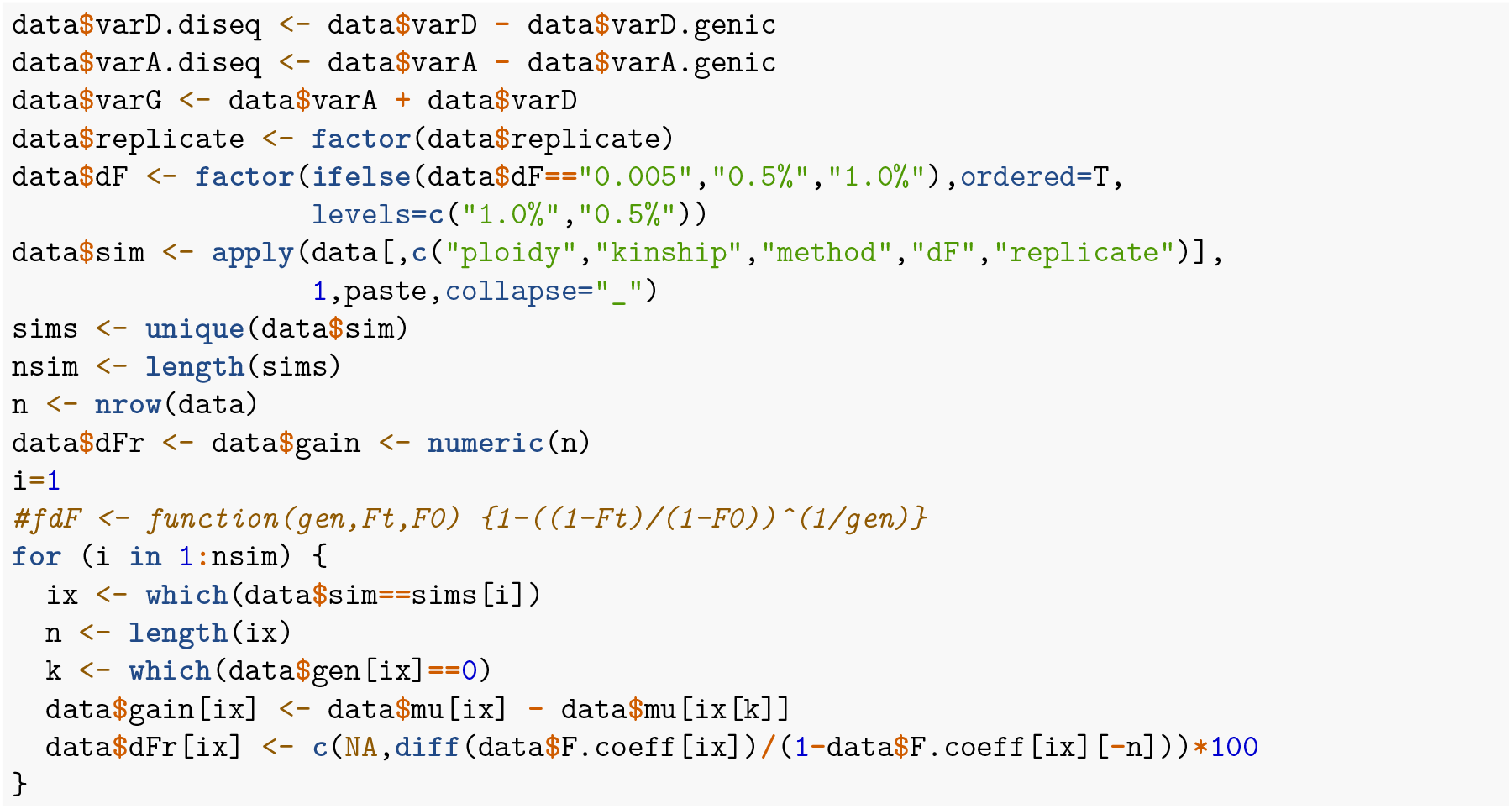

**Figure 3**

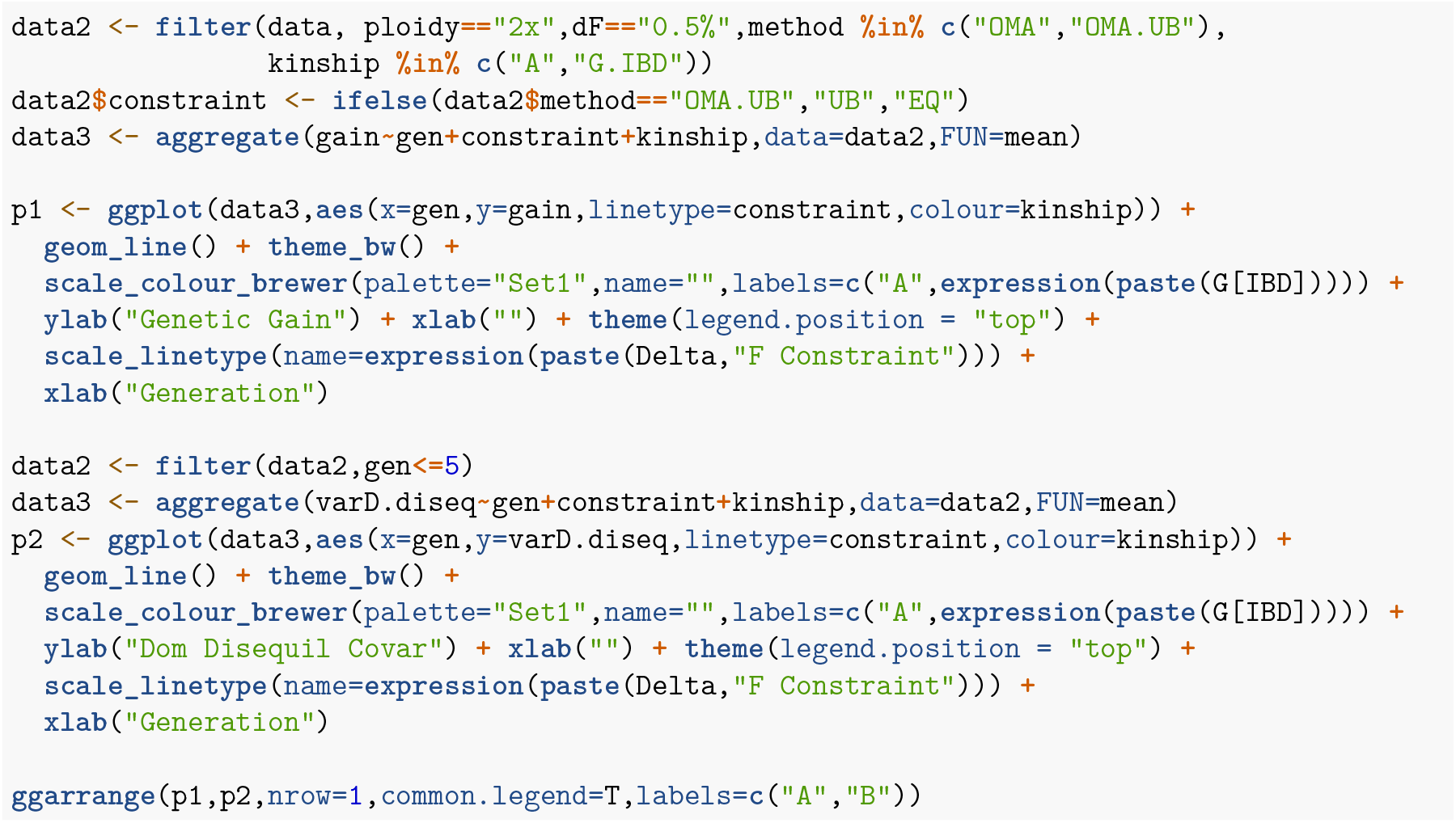
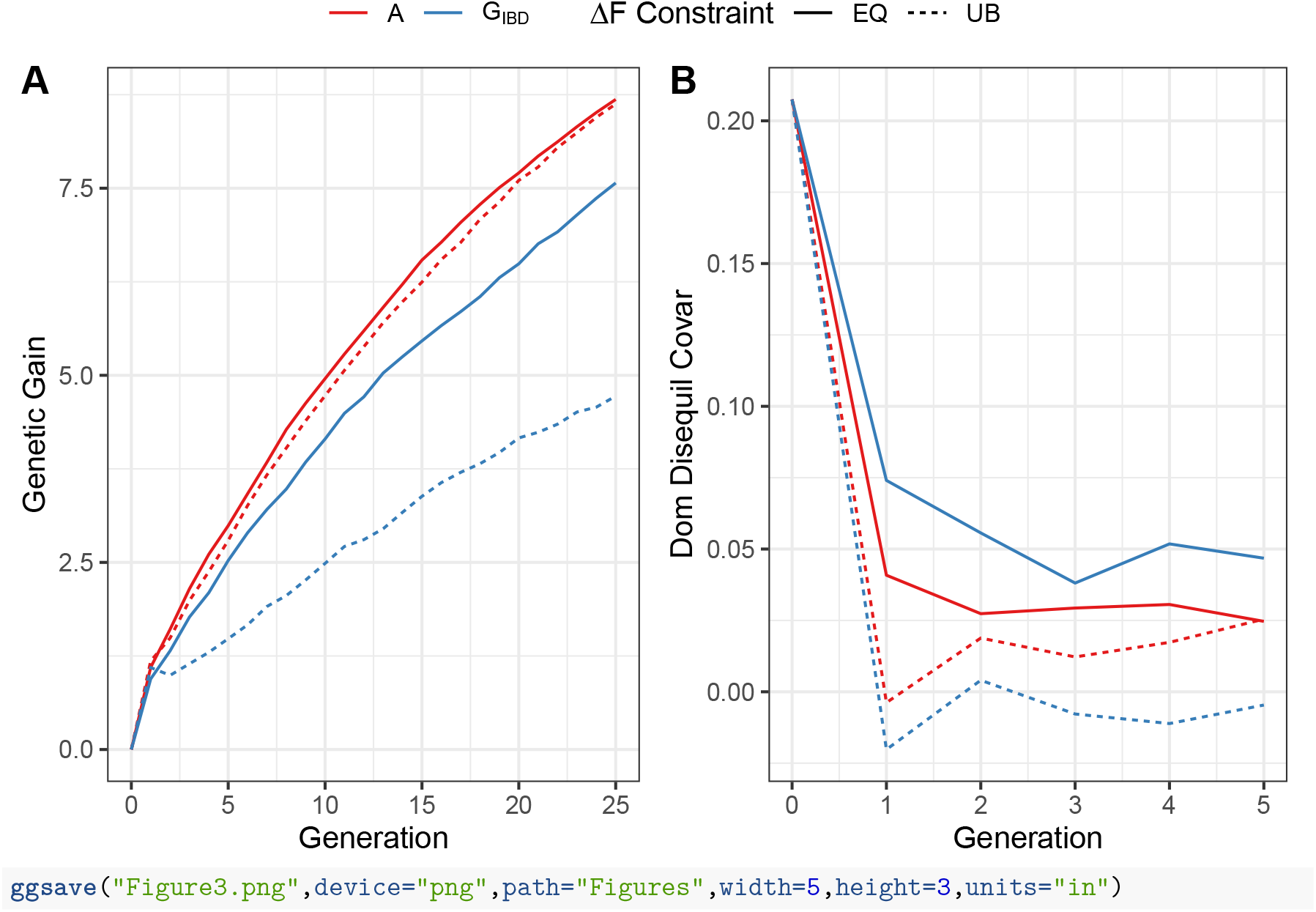

**Figure S2**

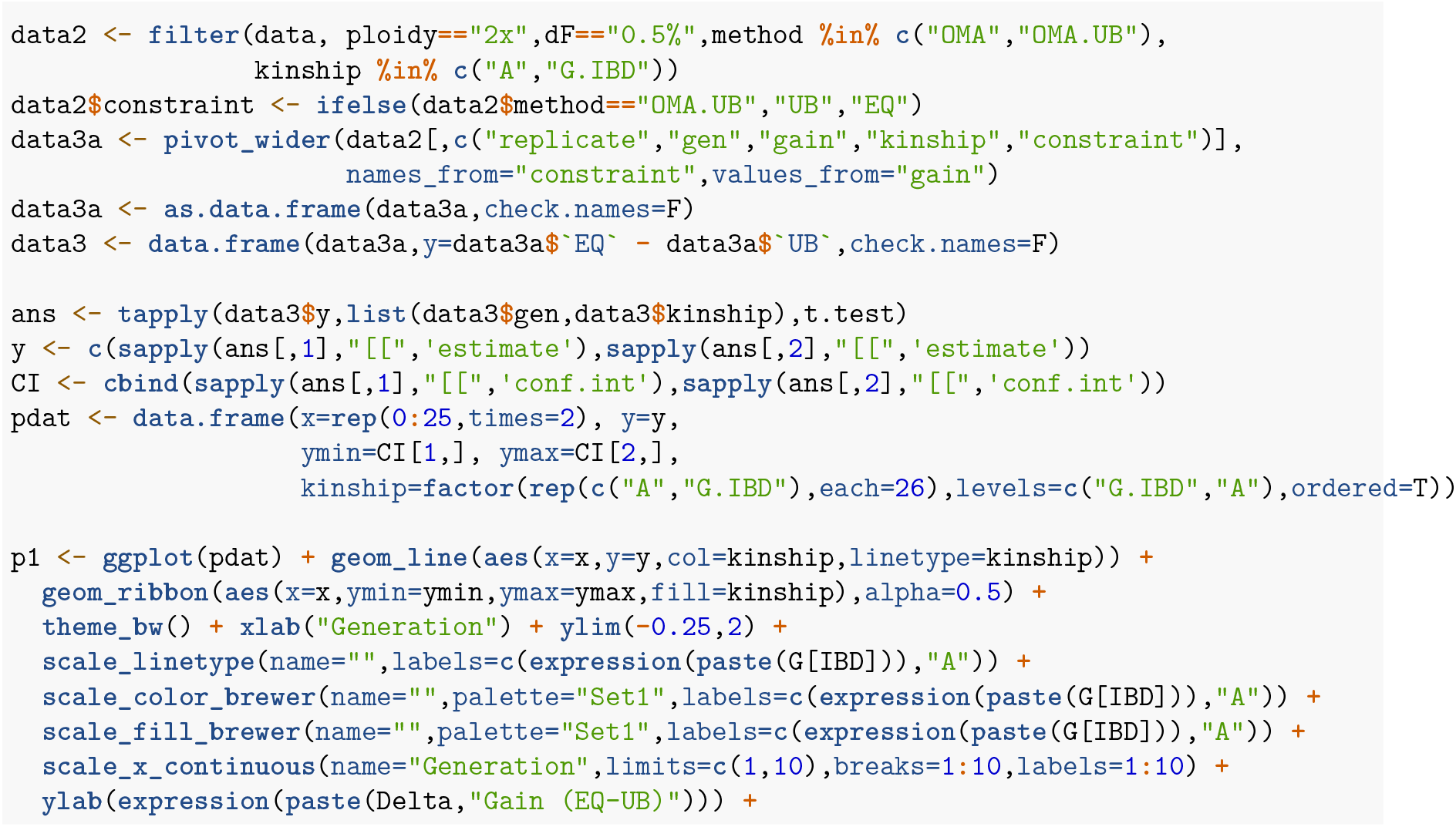
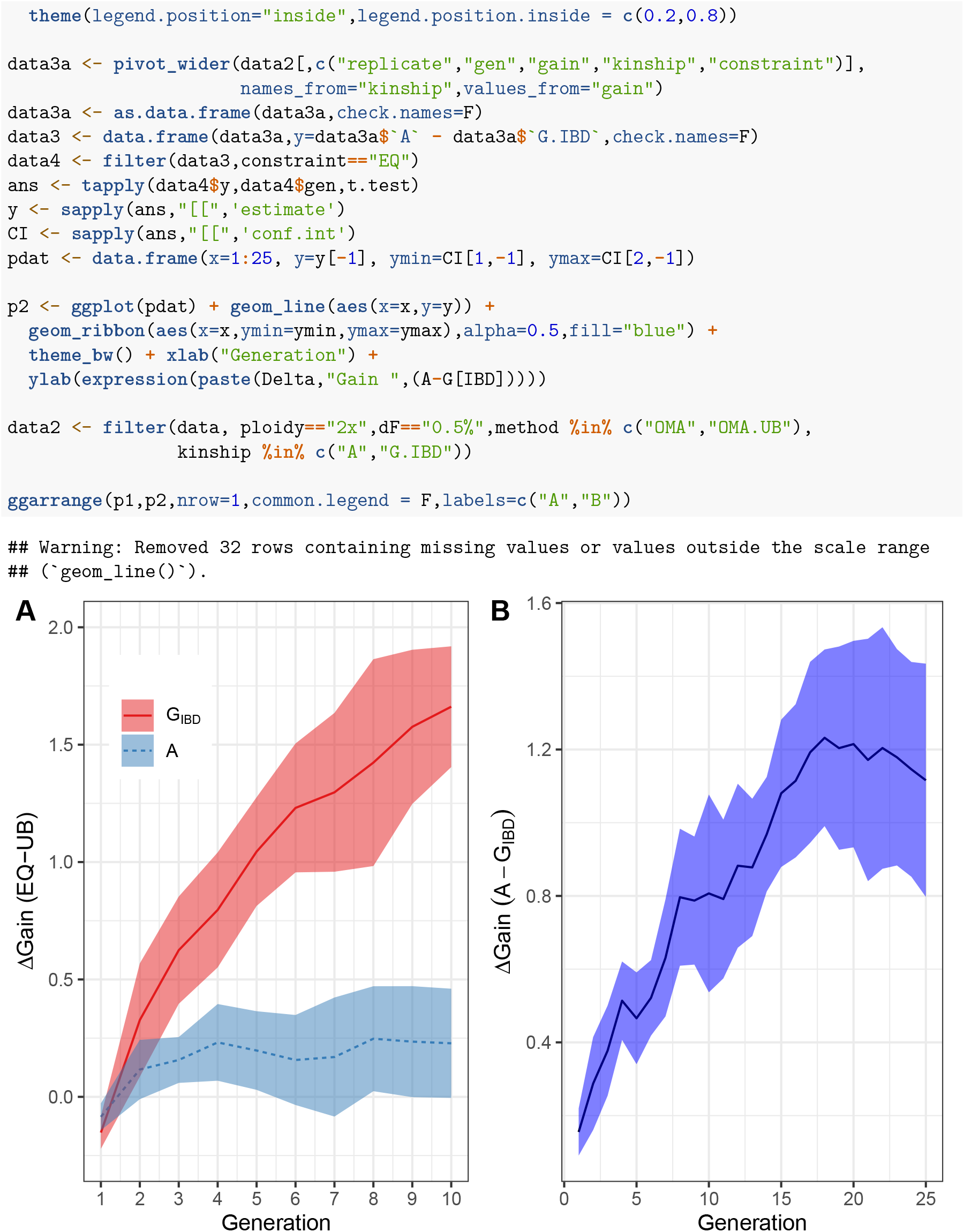
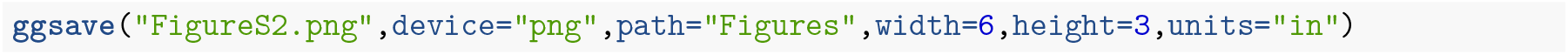

**Figure 4**

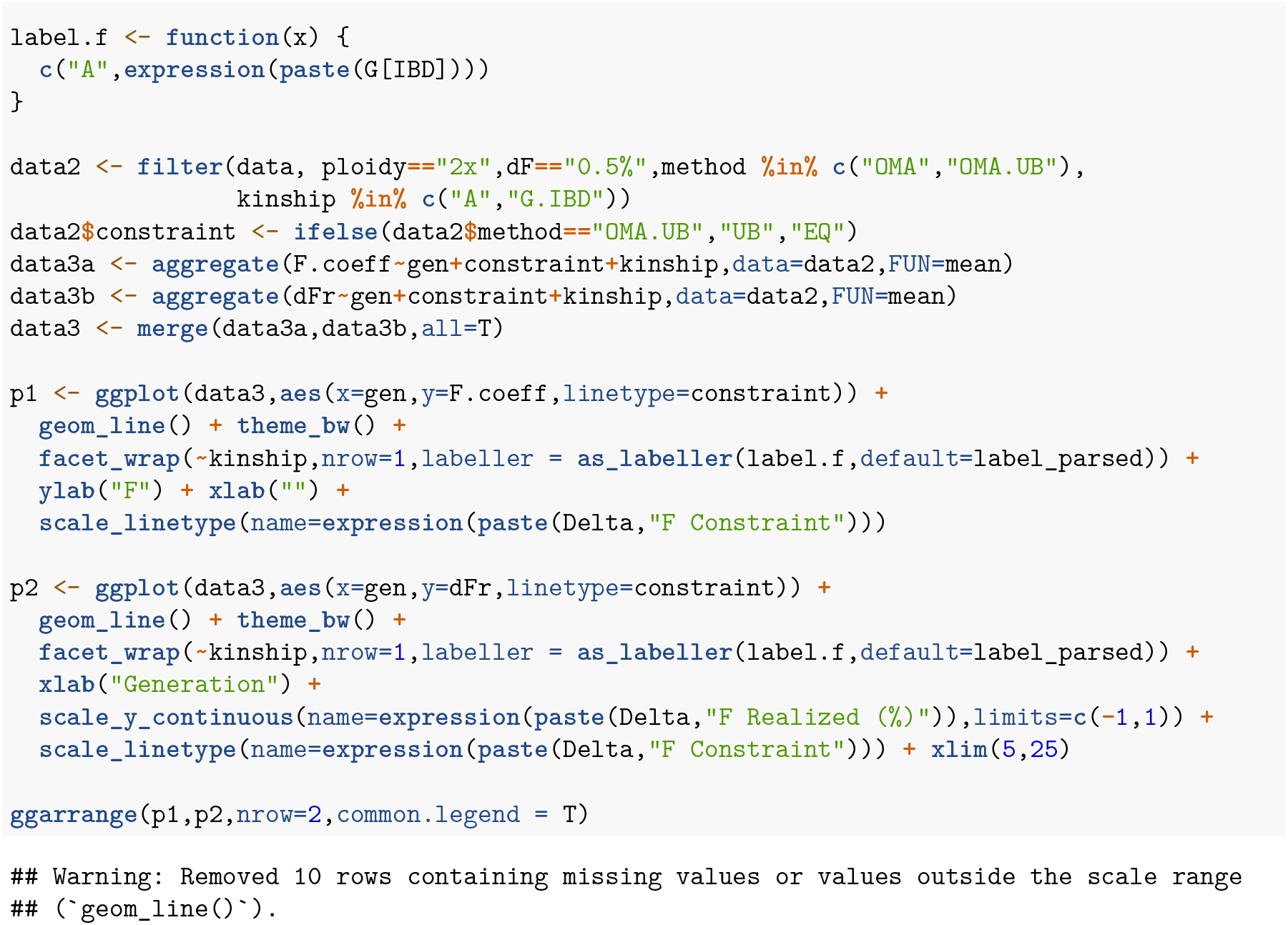
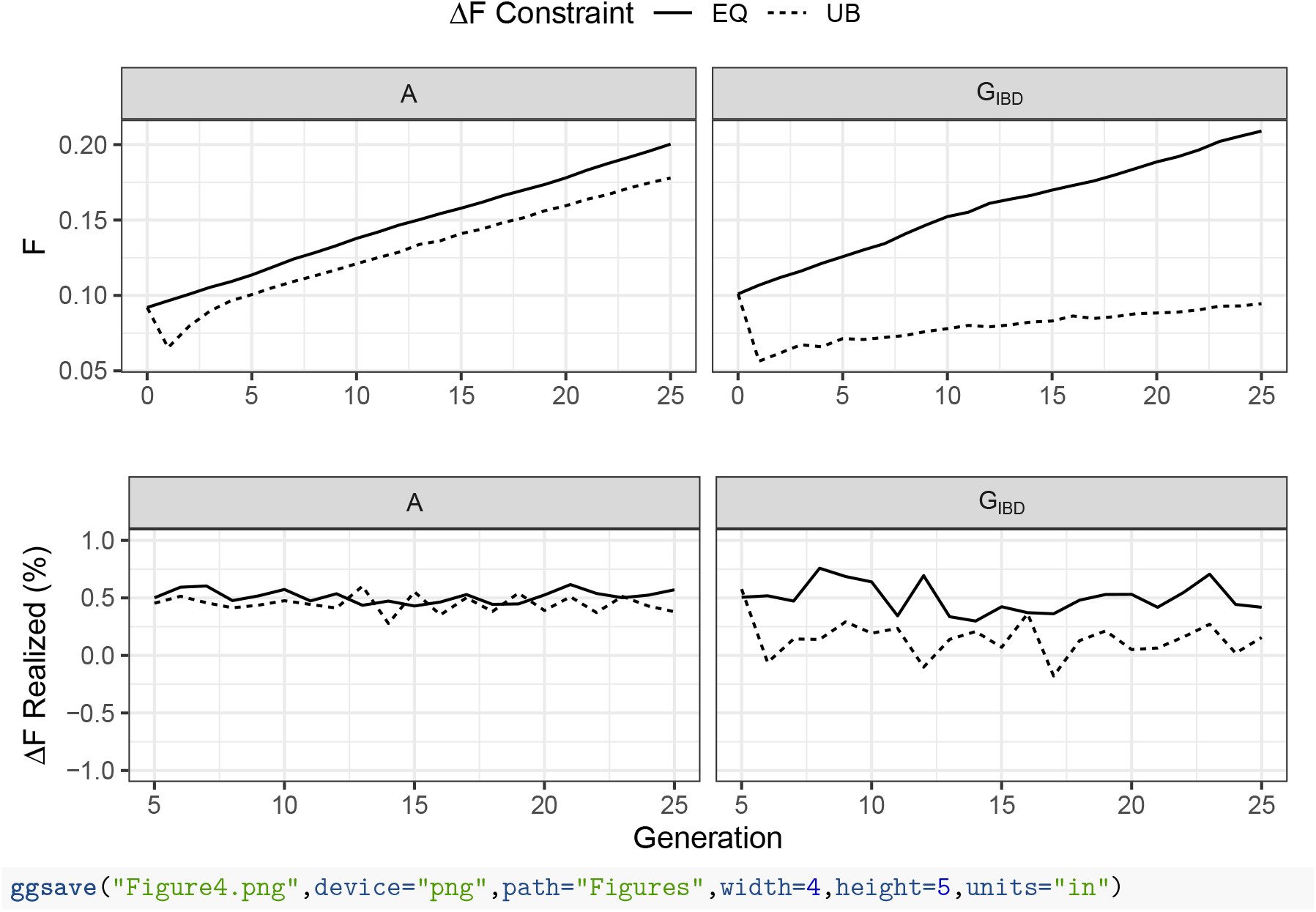

**Table S2**

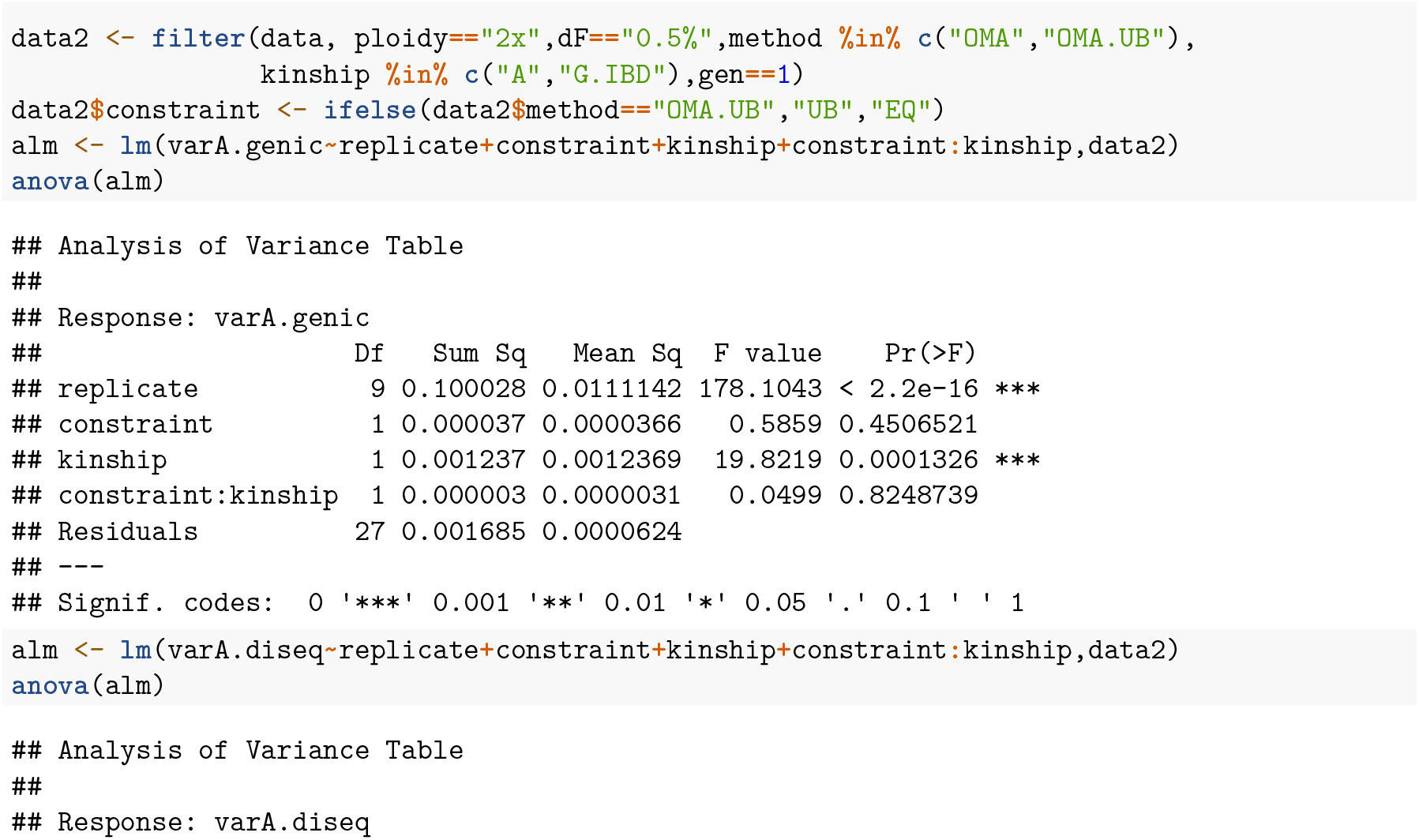
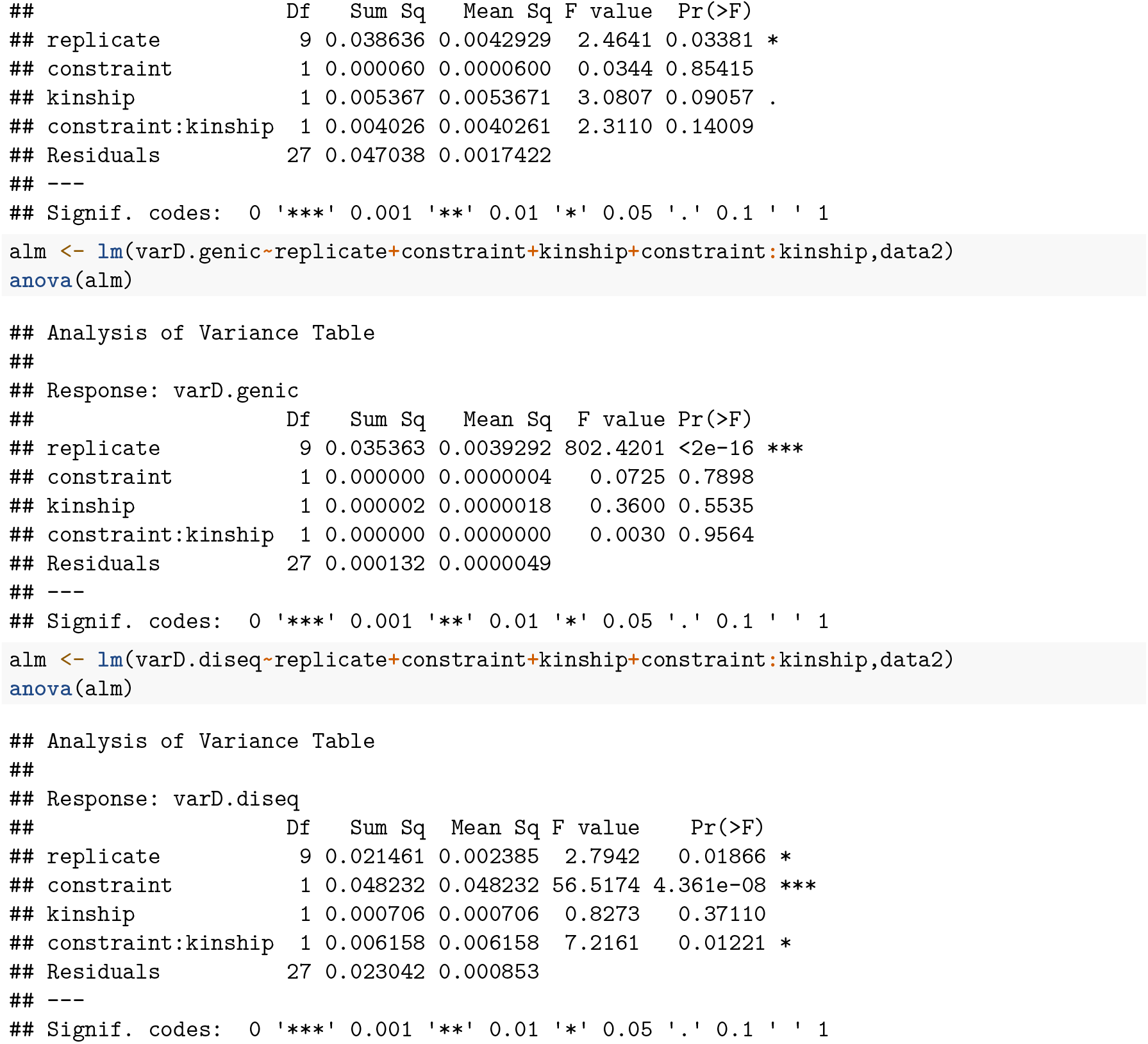

**Figure S3**

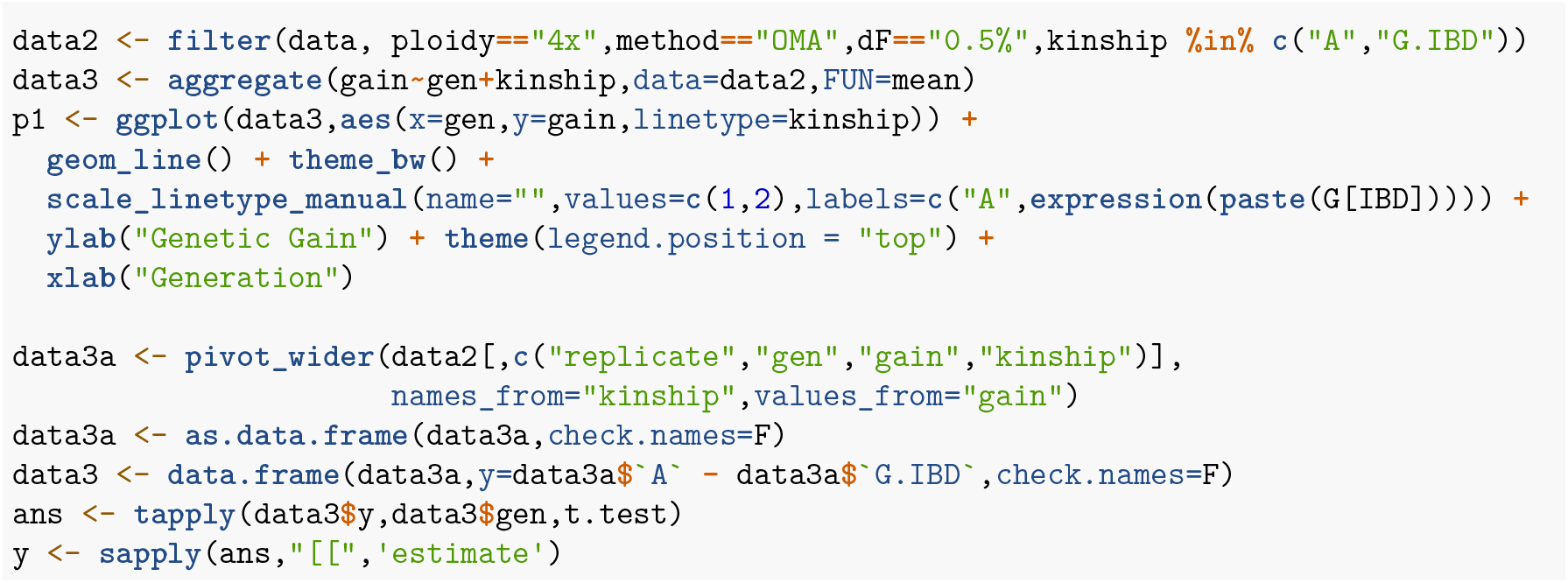
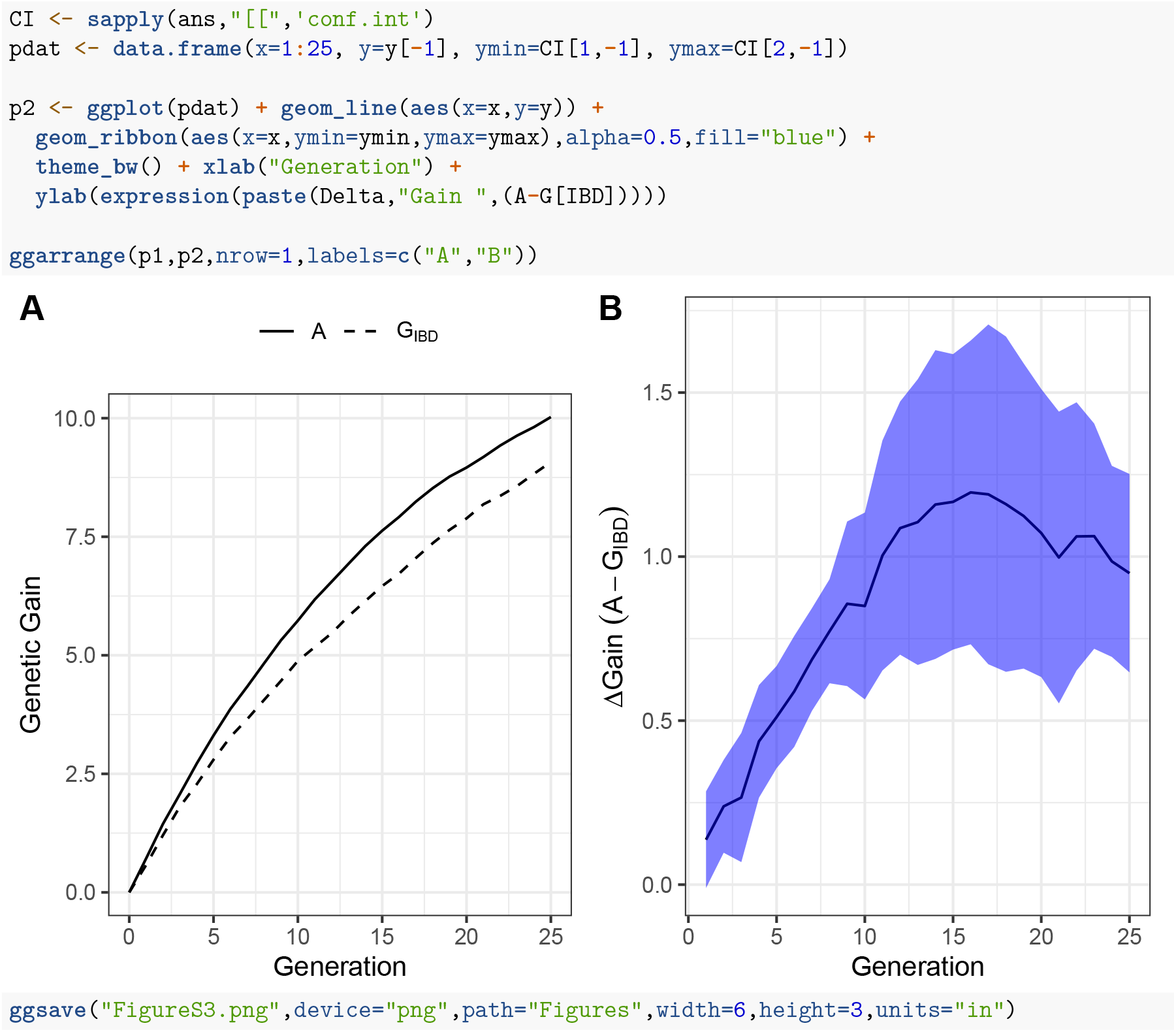

**Figure S4**

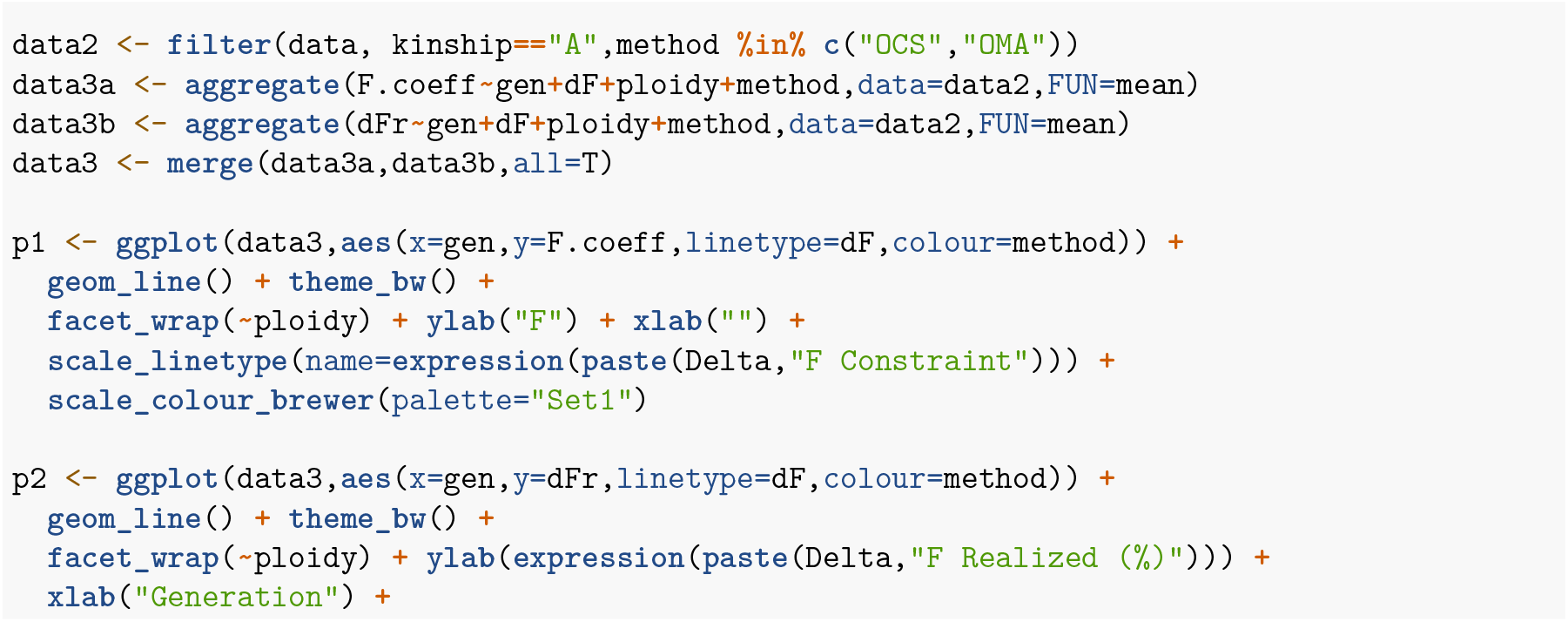
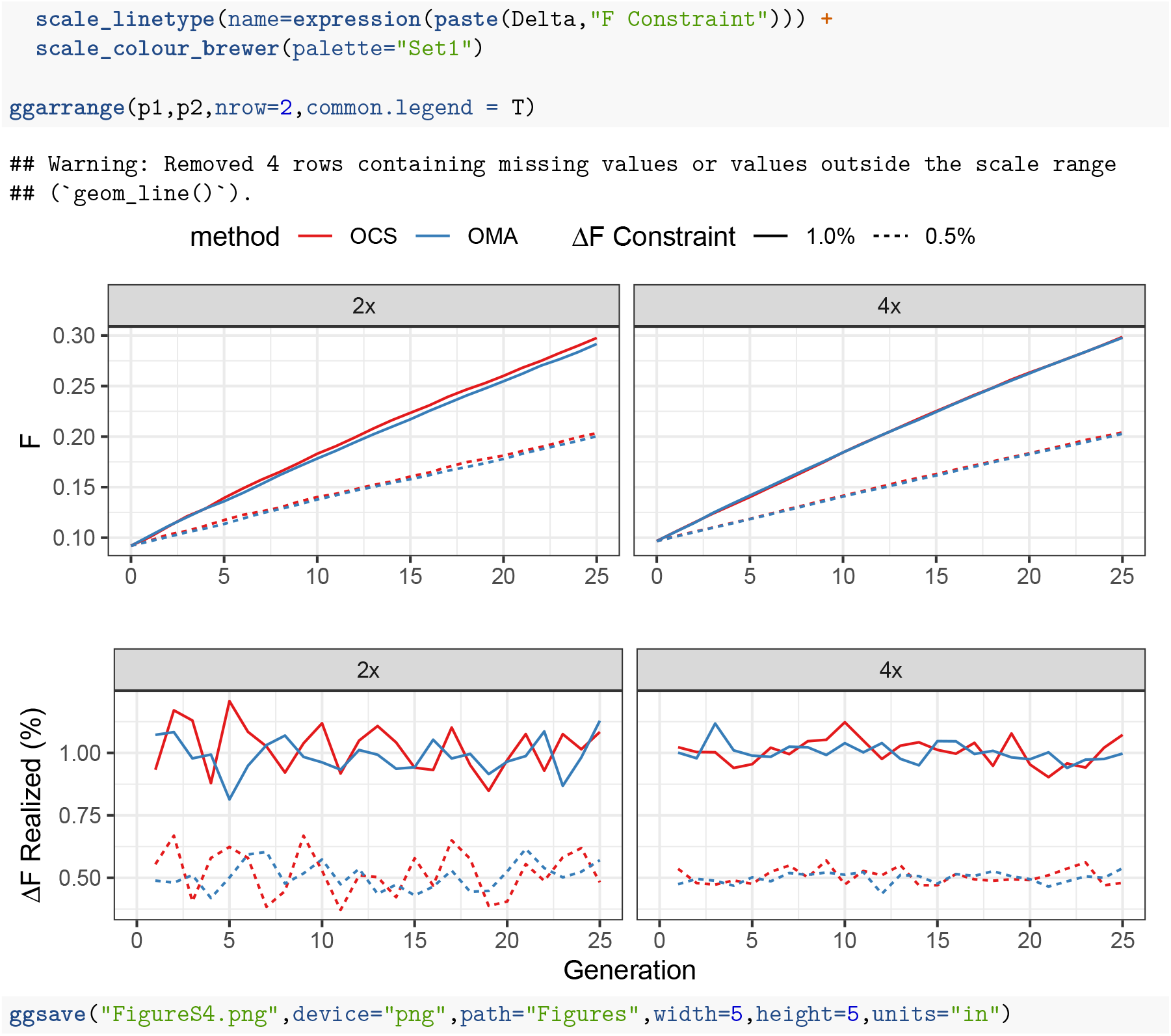

**Figure 5**

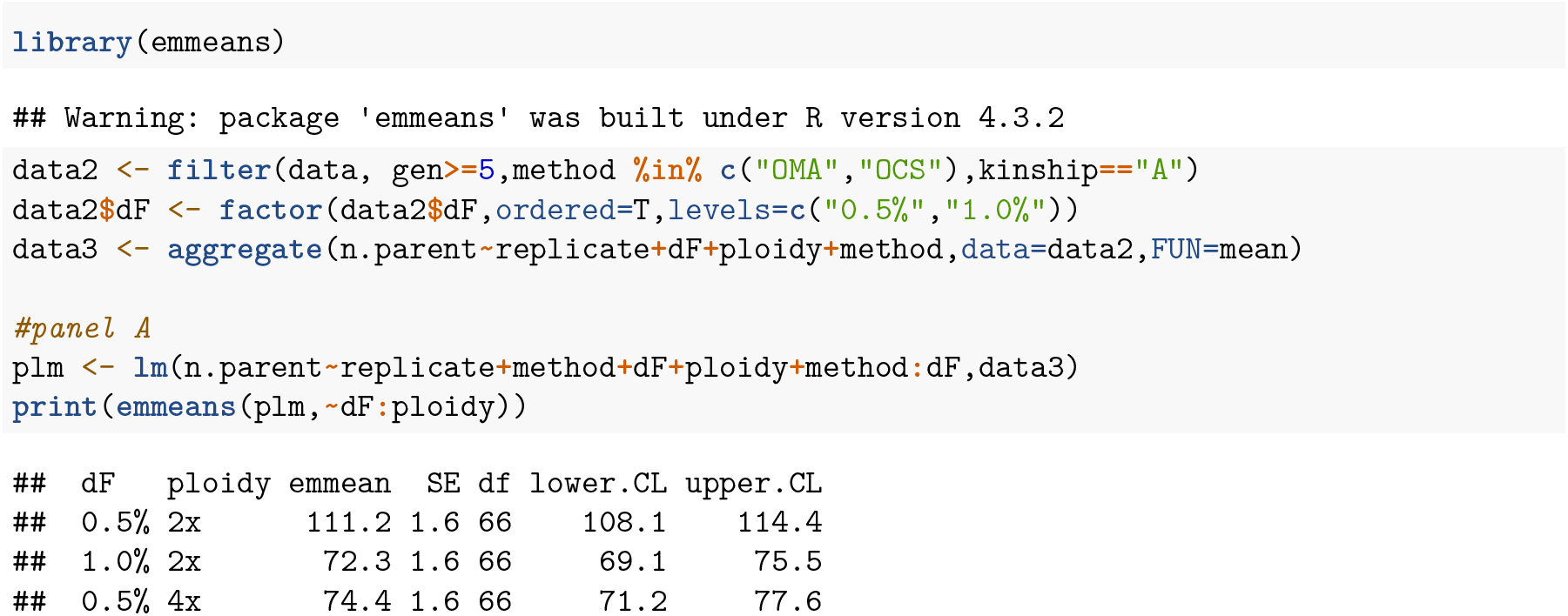
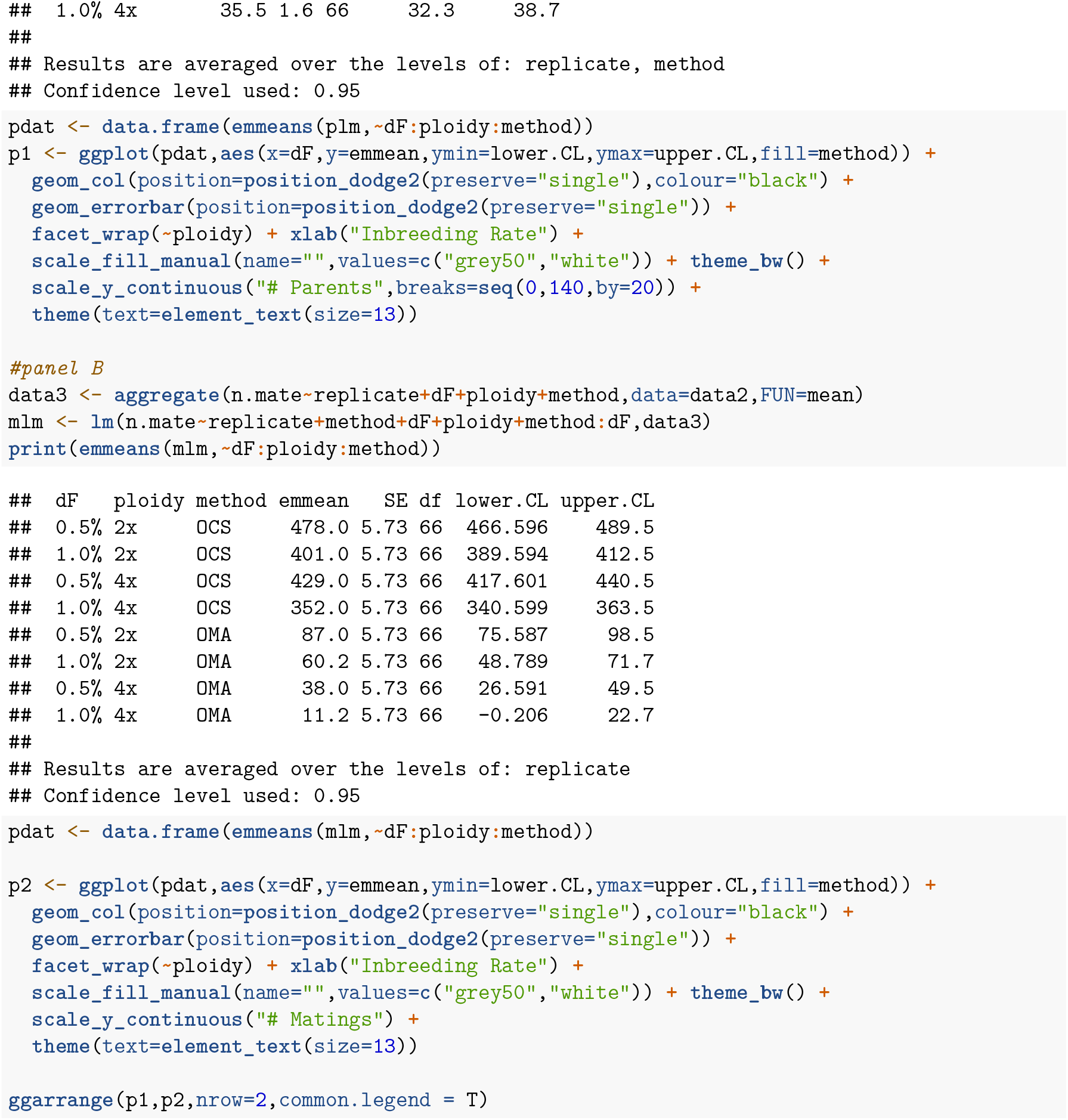
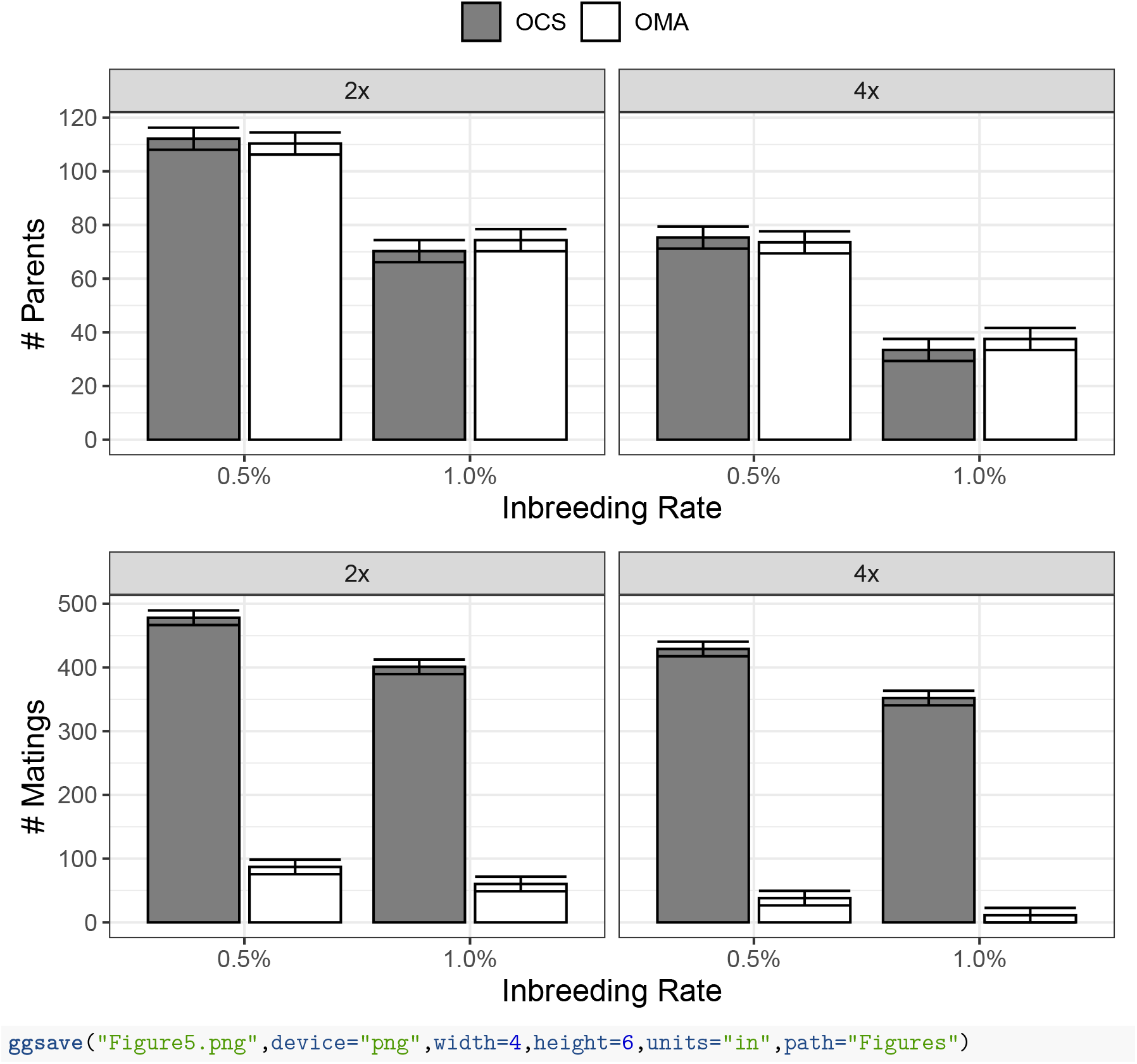

**Figure 6**

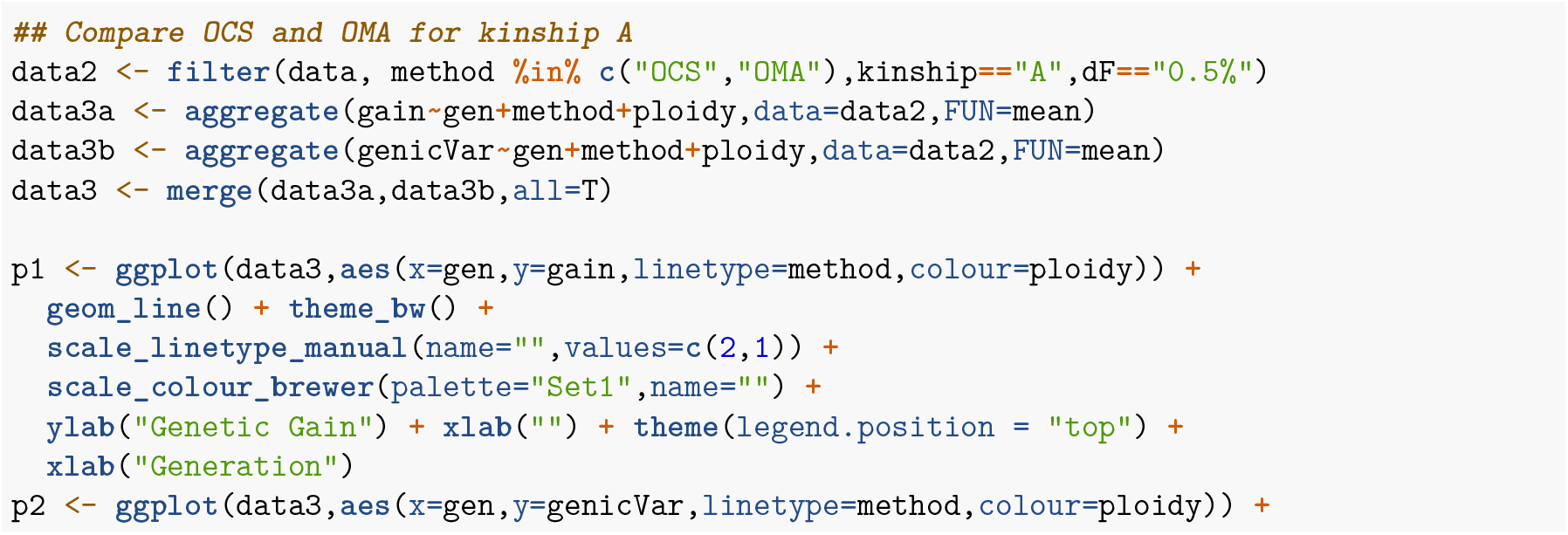
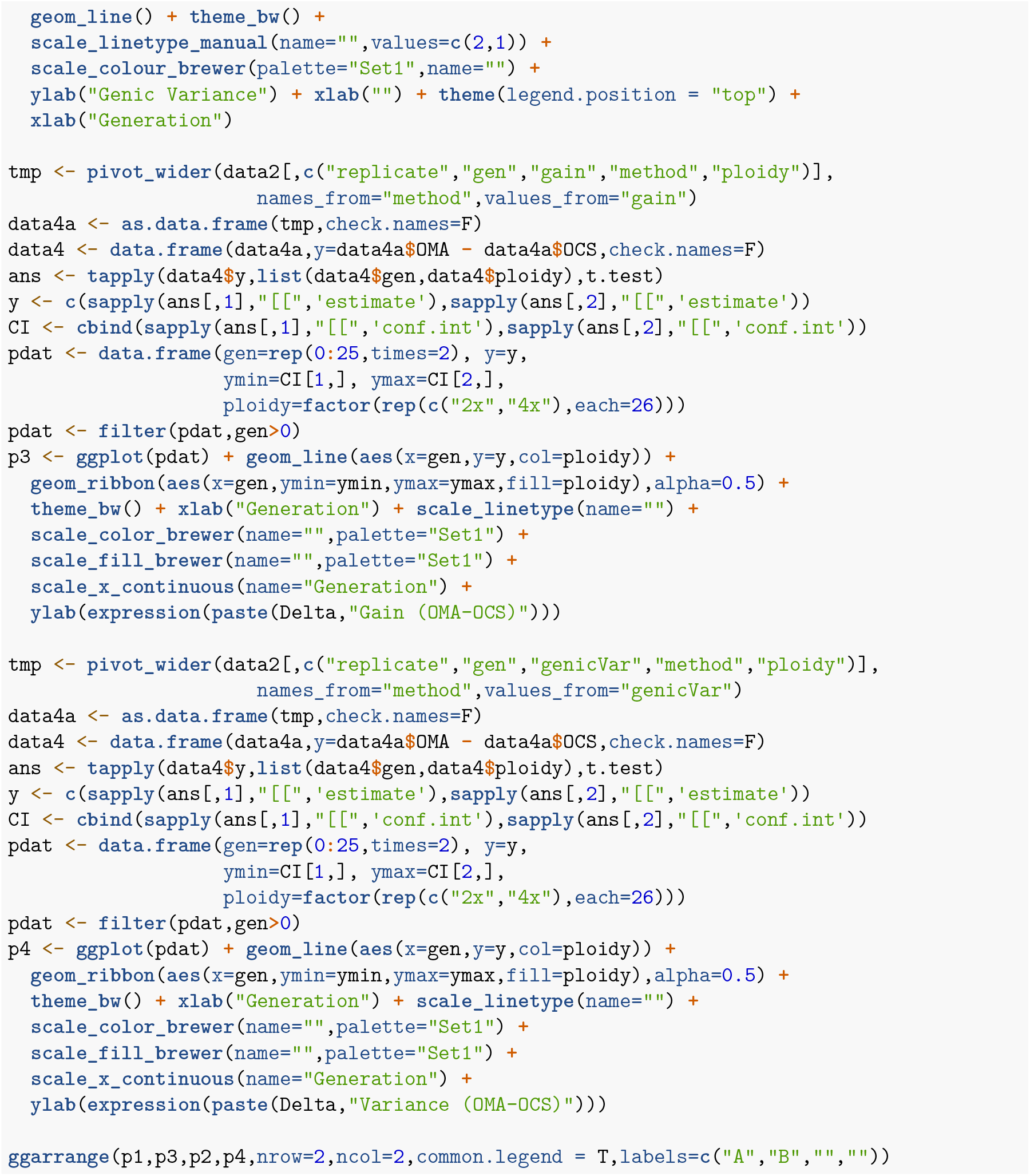
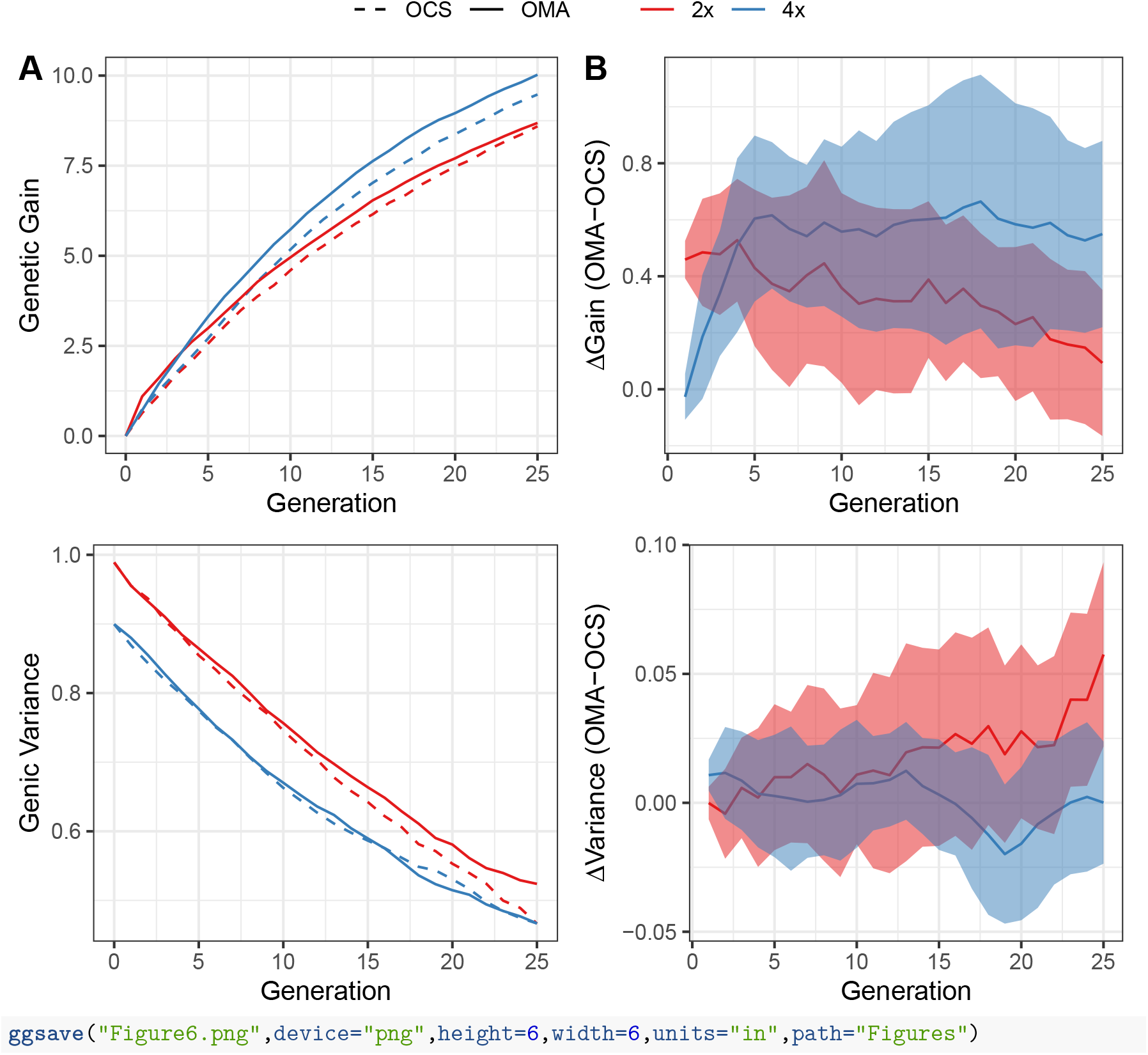

**Figure S5 1% inbreeding**

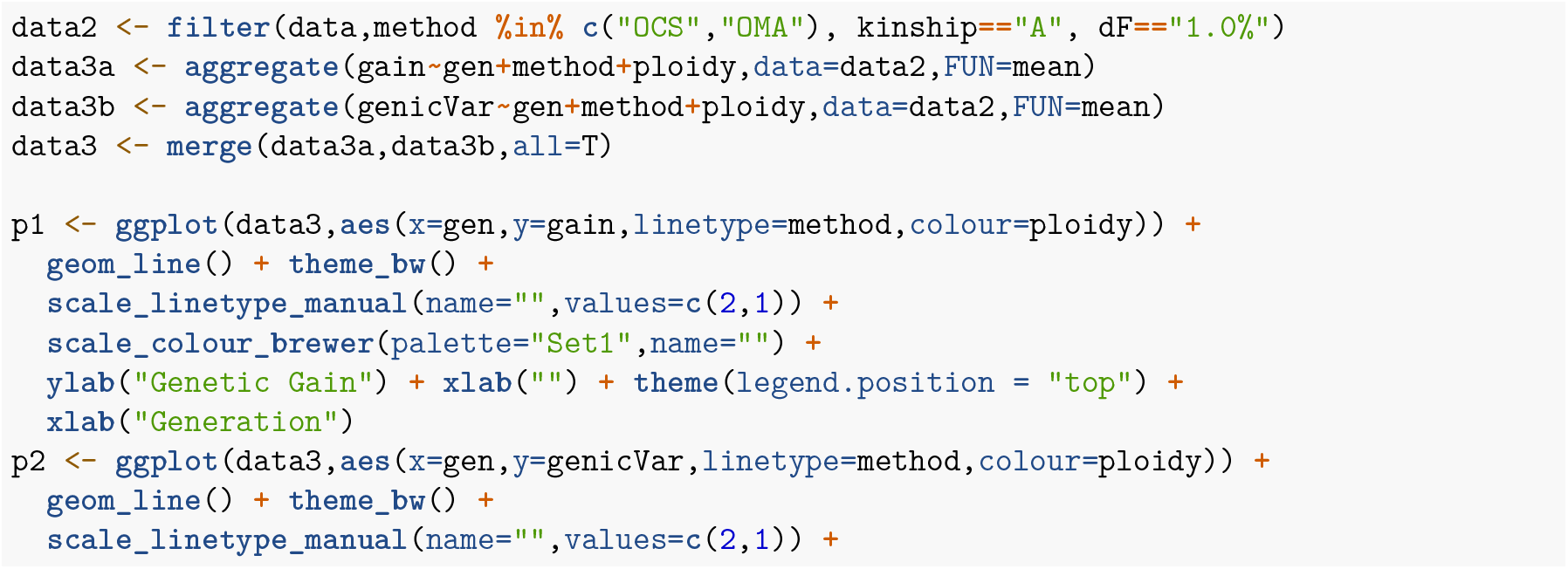
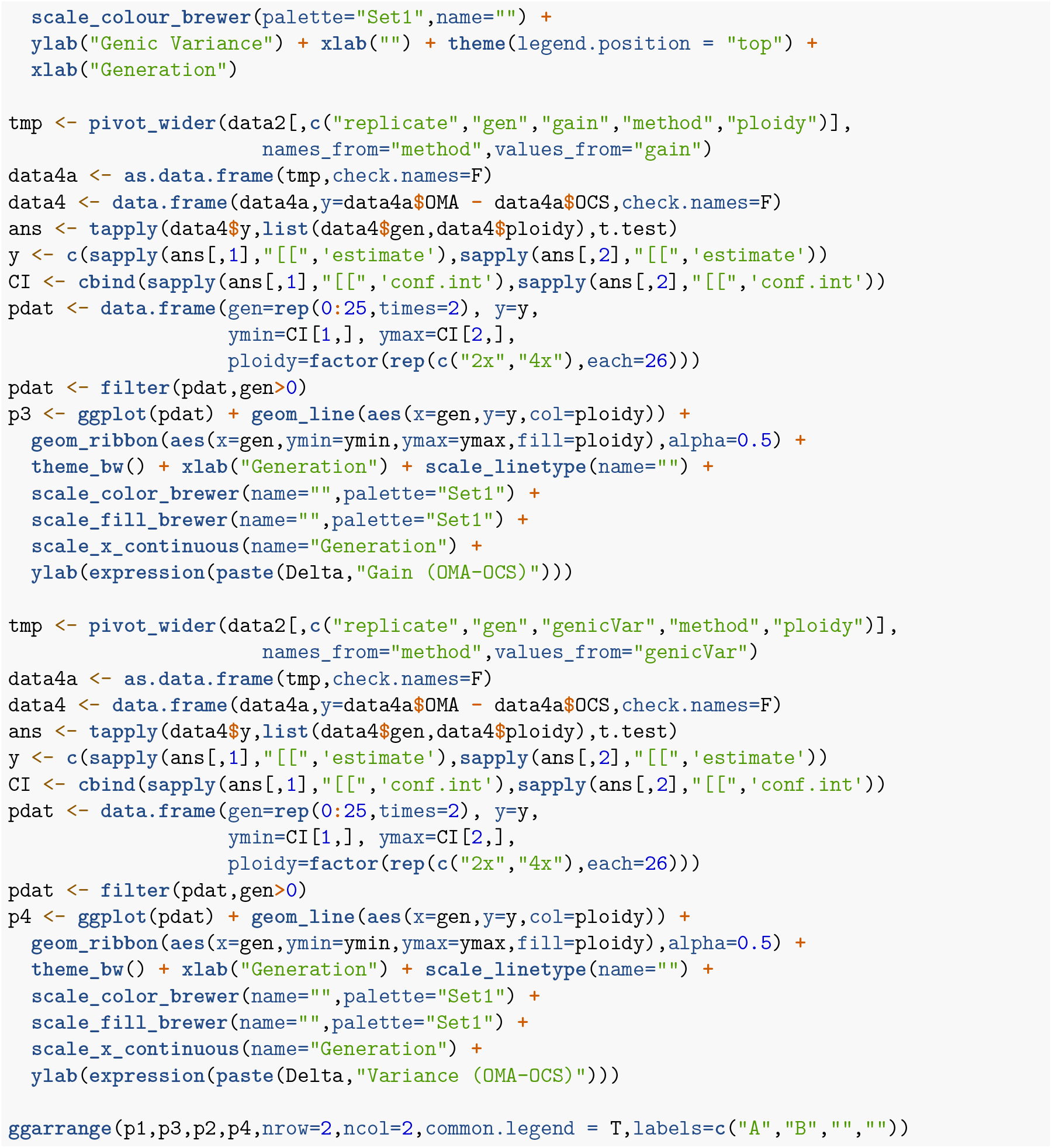
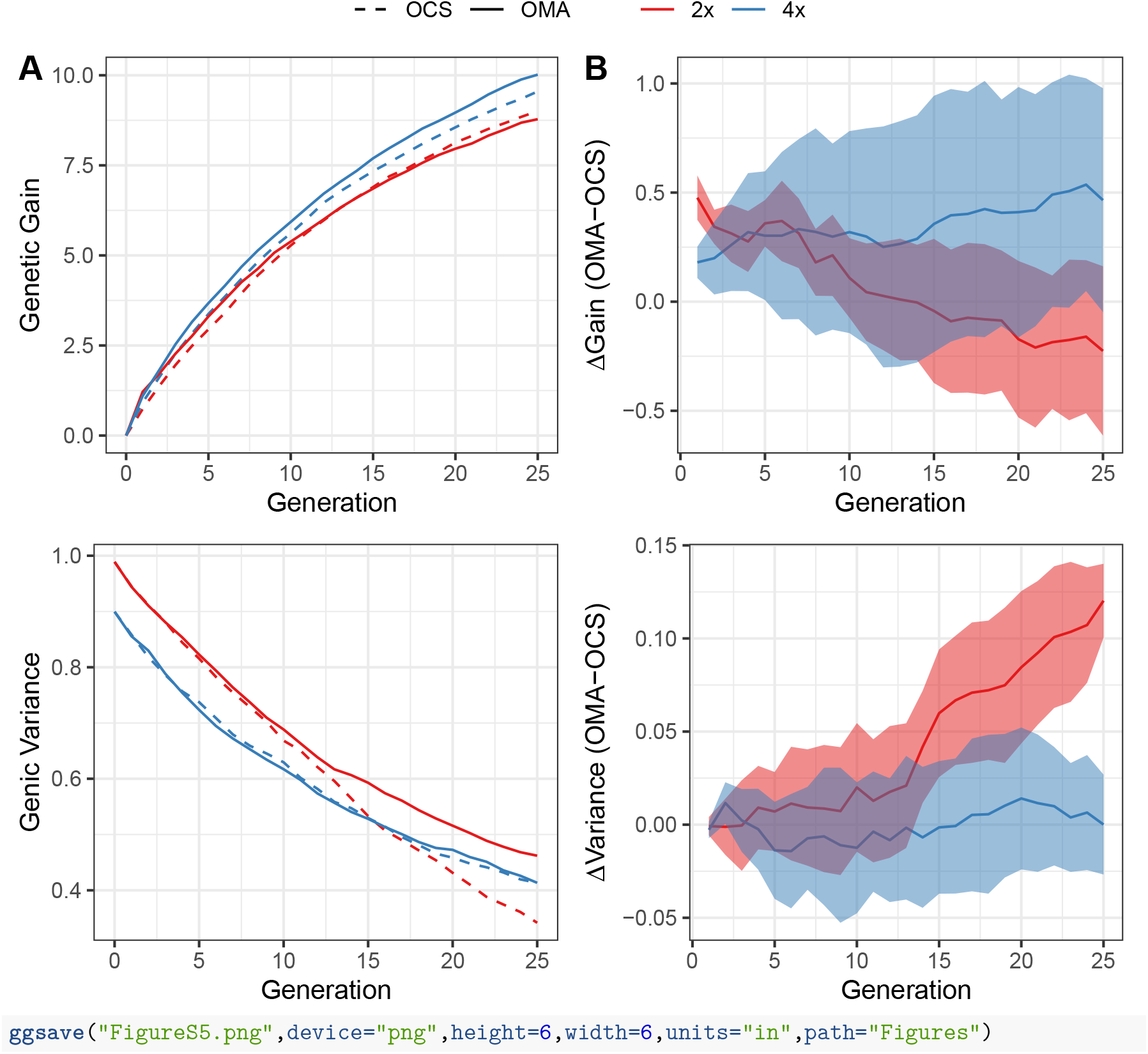

**Figure 7**

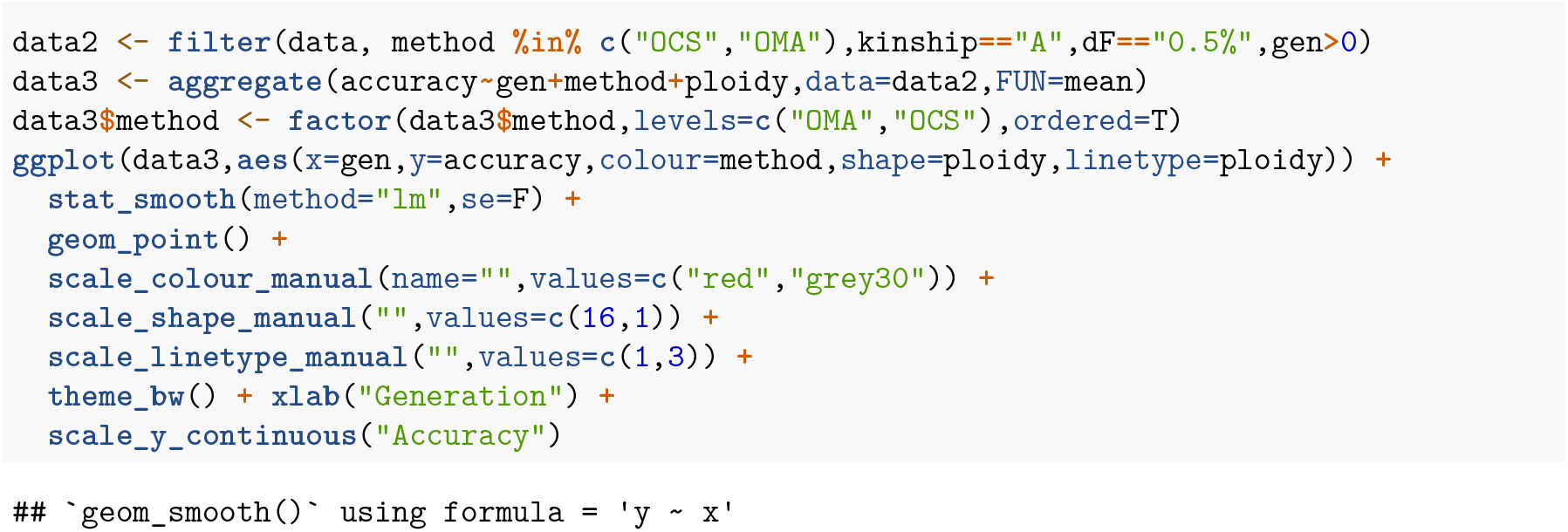
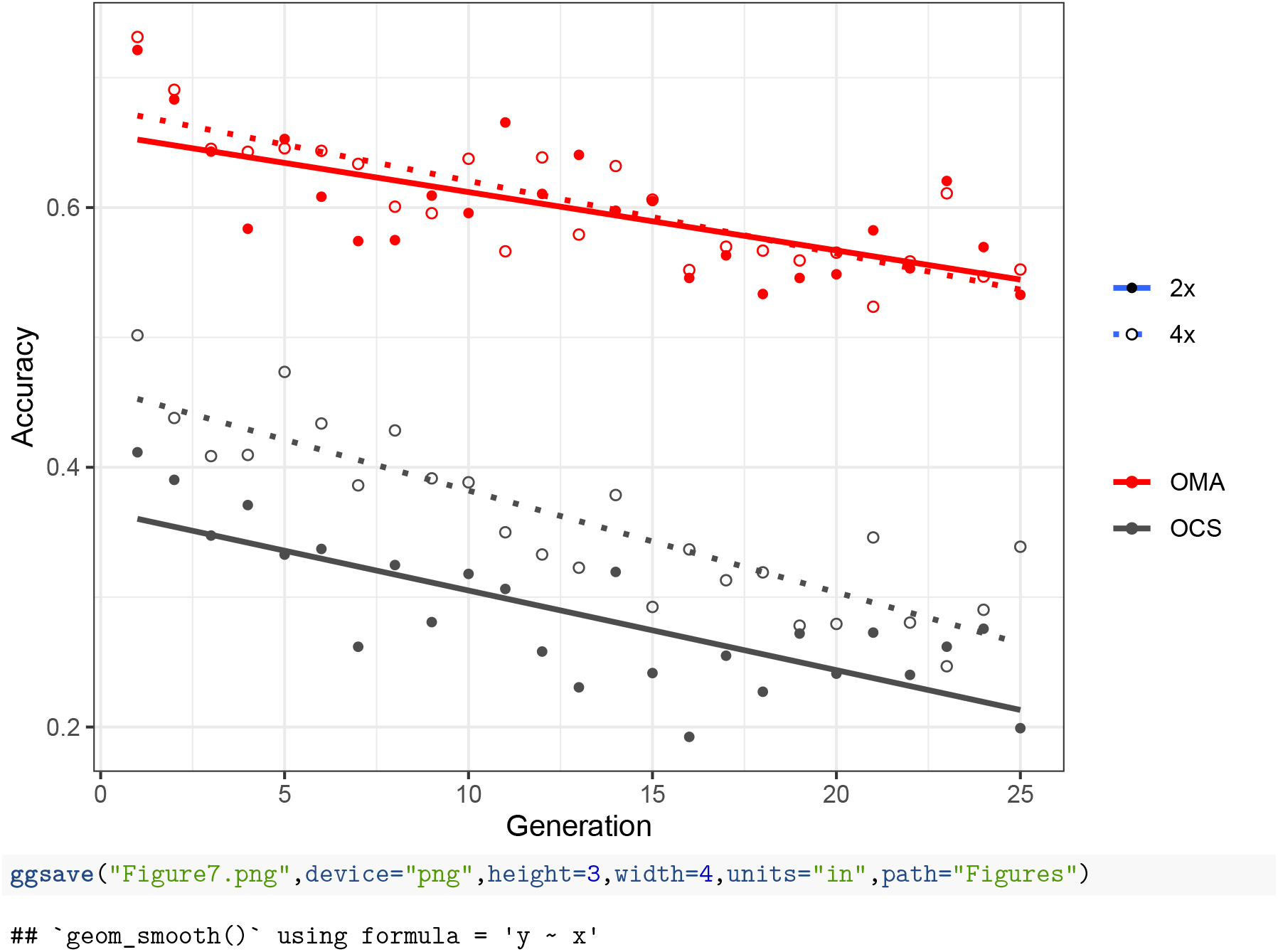

**Figures S6 G matrix**

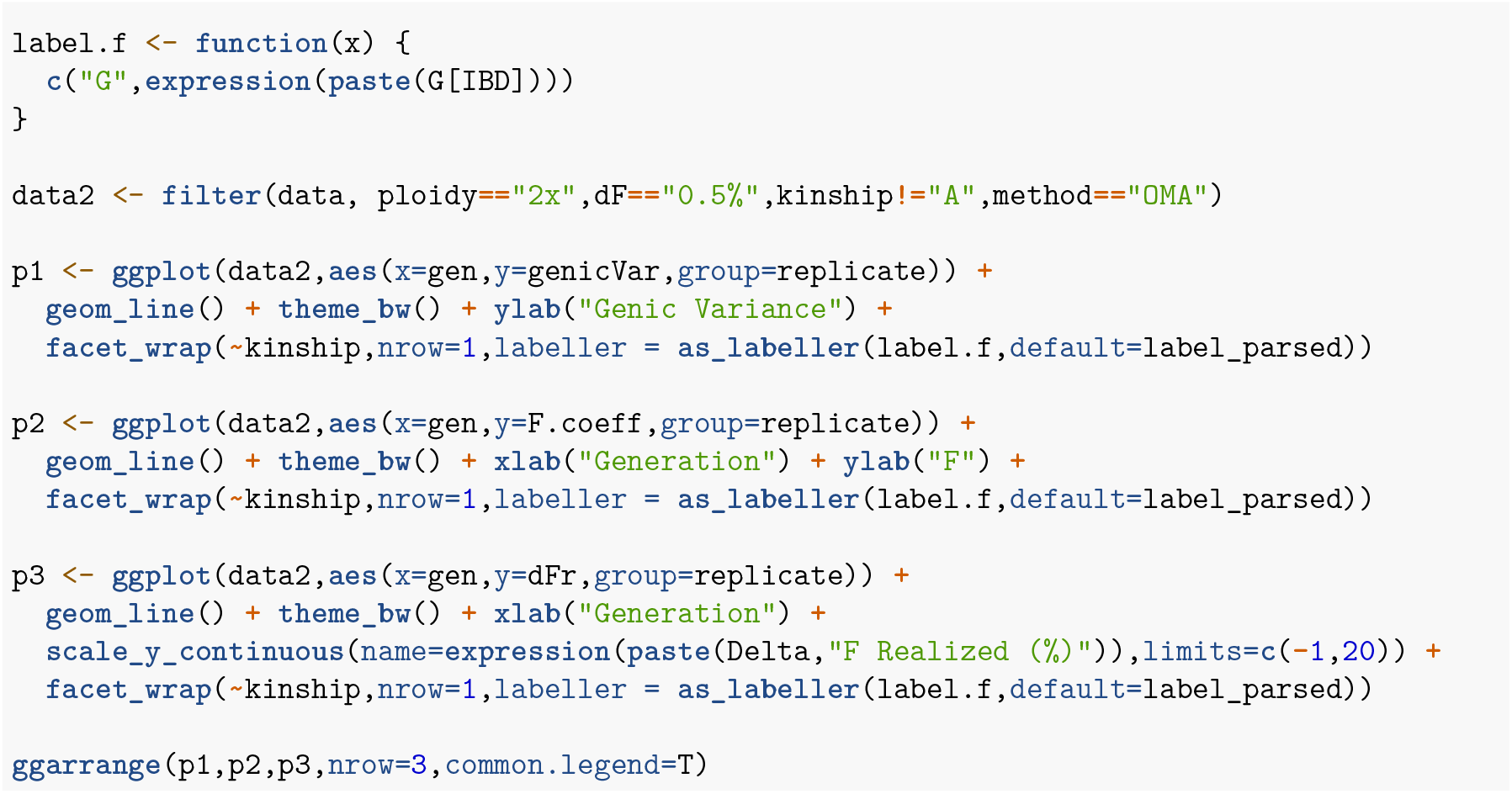
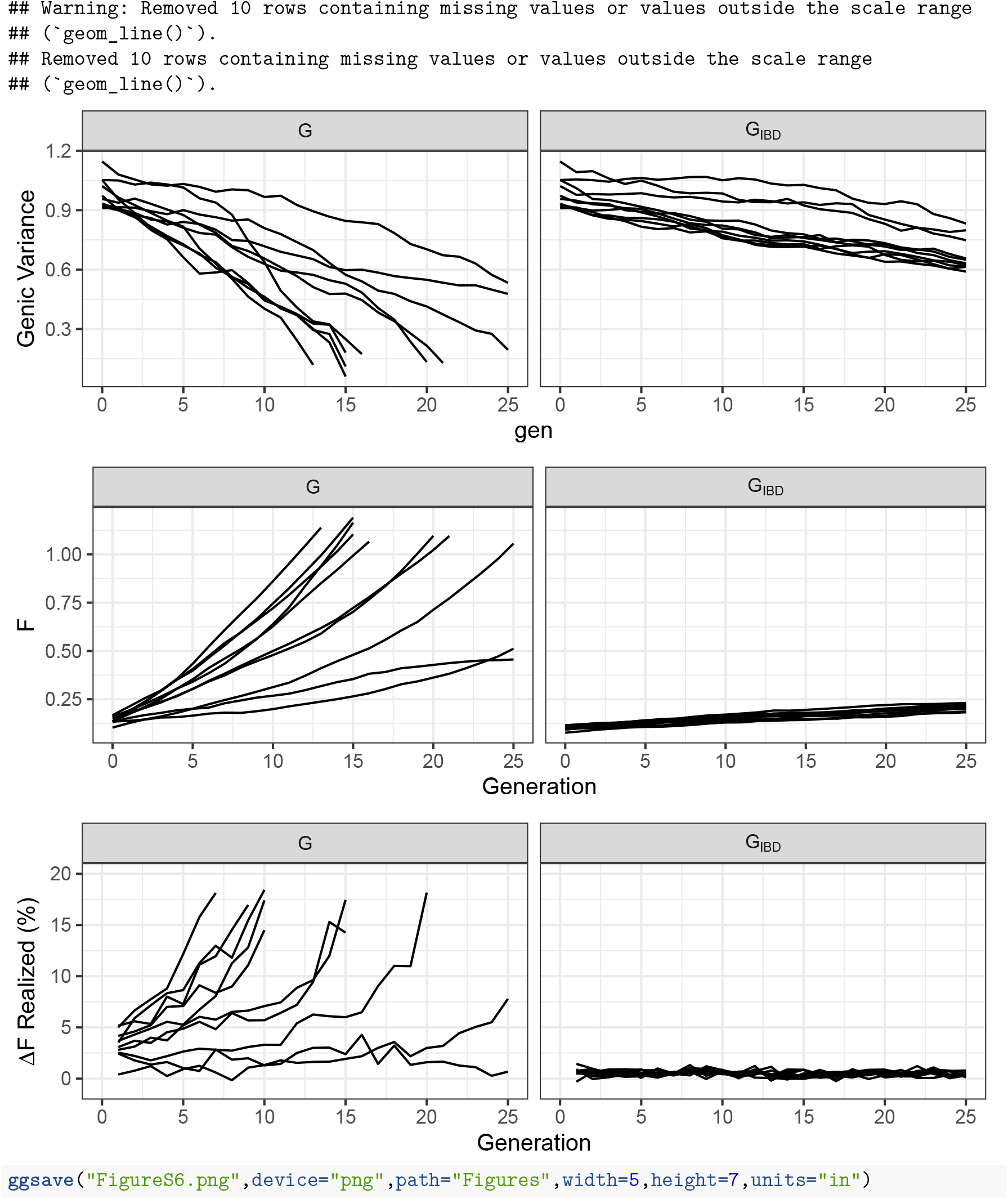

